# Abundance and diversity of phages, microbial taxa and antibiotic resistance genes in the sediments of the river Ganges through metagenomic approach

**DOI:** 10.1101/2020.04.29.067819

**Authors:** Narender Kumar, Amit Kumar Gupta, Sarabjeet Kour Sudan, Deepika Pal, Vinay Randhawa, Girish Sahni, Shanmugam Mayilraj, Manoj Kumar

**Affiliations:** Division of Protein Science & Engineering, Institute of Microbial Technology, Council of Scientific & Industrial Research (CSIR), Chandigarh, 160 036, India; Virology Unit and Bioinformatics Centre, Institute of Microbial Technology, Council of Scientific & Industrial Research (CSIR), Chandigarh, 160 036, India; MTCC-Microbial Type Culture Collection & Gene Bank, Institute of Microbial Technology, Council of Scientific & Industrial Research (CSIR), Chandigarh, 160 036, India

**Keywords:** Phages, Bacteria, Antibiotic Resistance Genes, The Ganges River, Sediment, Metagenomics

## Abstract

In the present study, we have analyzed the metagenomic DNA from the pooled sediment sample of the river Ganges to explore the abundance and diversity of phages, microbial community and antibiotic resistance genes. Utilizing data from Illumina platform, 4174 (∼0.0013%) reads were classified for the 285 different DNA viruses largely dominated by the group of 260 distinctive phages (3602 reads, ∼86.3%). Among all, *Microcystis* (782 hits), *Haemophilus* (403), *Synechococcus* (386), *Pseudomonas* (279), *Enterococcus* (232), *Bacillus* (196), *Rhodococcus* (166), *Caulobacter* (163), *Salmonella* (146), *Enterobacteria* (143), *Mycobacterium* (128), *Propionibacterium* (71), *Erwinia* (70), *Ralstonia* (56) phages shows the highest abundance and account for approximately 90% of the total identified phages. Additionally, we have also identified corresponding host pertaining to these phages. Mainly, *Proteobacteria* (∼69.3%) dominates the microbial population structure. Primarily orders such as *Caulobacterales* (∼28%), *Burkholderiales* (∼13.9%), *Actinomycetales* (∼13.7%), *Pseudomonadales* (∼7.5%) signify the core section. Further, 21869 (∼0.00695%) reads were classified in 20 ARG types (classes) and 240 ARGs (subtypes) among which 4 ARG types namely multidrug resistance (MDR) (12041 reads, ∼55%), bacitracin (3202 reads, ∼15%), macrolide-lincosamide-streptogramin (MLS) (1744 reads, ∼7.98%), and fosmidomycin (990 reads, ∼4.53%) has the highest abundance. Simultaneously, six resistance mechanisms were also recognized with the dominance of antibiotic efflux (72.8%, 15919 reads). The results unveil the distribution of (pro)-phages; microbial community and various ARGs in the Ganges river sediments. Further research on these identified phage(s) could be used in phage-based therapeutics against pathogenic bacteria.

## Introduction

Viruses are key players in human health and diseases [1]. Viruses targeting the bacteria (bacteriophages) are apparently one of the most abundant biological entities on the planet [1-6]. Phages play cardinal role in almost every ecosystem including natural environment, human microbiota etc. [6-9]. They are known for significantly influencing microbial community structure, biogeochemical nutrient cycles [10-12]. They can also act as vehicle for exchange of genetic material between bacteria (transduction), mobilization of antibiotic resistance genes (ARGs) etc. [13]. Phages also have evolutionary impact as they can facilitate recombination event, integration as prophages etc. [10, 14]. Furthermore, they are also effective alternative to antibiotics and can be utilized as therapy [15, 16]. To this regard, proper understanding and exploration of virome is indispensable and critically important to identify viral pathogens and phage community [1, 11, 17-20].

Various traditional molecular assays/techniques were utilized to detect viruses in samples (clinical culture or environmental), which include polymerase chain reaction (PCR), real time PCR (qPCR), reverse transcription polymerase chain reaction (RT-PCR) etc. However, these conventional techniques do not allow wide monitoring and discovery of viral communities. To overcome these limitations, viral metagenomics approach using high throughput sequencing platforms (454, Illumina, PacBio, Nanopore etc.) provided a great hand and instrumental for understanding viral (microbial) world [3, 14, 18]. Various studies have supported the application of metagenomics to investigate the viral ecology, pathogen surveillance and to elucidate viral evolution and diversity [3, 21-27]. Moreover, it is of prime interest to get insight into the viromes specifically phage communities from different environments as it may affect the microbial diversity of the area and could assist in the future research of phage based therapeutics to combat bacterial pathogens [14, 28-34]. We still have inadequate knowledge and understanding of phage diversity in different ecosystems, which remains to be fully explored.

Further, it is also necessary to understand distribution of microbial community for public and environmental health and ARGs to frame proficient strategies against them. The ability of pathogenic bacteria to develop resistance against antibiotics by acquiring mutations or resistance genes is a major concern and threat to human health worldwide [35-38]. Various studies advocate the abundance of distinct antibiotic resistance genes (ARGs) and their reservoirs. There are several environmental sources i.e., soil [39, 40], sediments [41, 42], sewage [43], activated sludge [44], human gut microbiota [45], human feces [46], animal waste [46], drinking water [47], surface water [48, 49], river water [50], wastewater treatment plants (WWTPs) [51, 52], glaciers [53] etc. are known to have diverse type of ARGs and considered to be hotspots for same. Moreover, natural resistome could be a most prominent source of the pathogenic resistance genes [54]. Furthermore, selective pressure of used antibiotics [36, 55] and horizontal gene transfer (HGT) of ARGs mediated via mobile genetic elements (MGEs) such as transposons, plasmids, phages [56, 57] etc. through conjugation, transduction and transformation make the situation even worse [36, 55, 57].

In this study, we are exploring and investigating the pooled sediments of the Ganges River. To best of our knowledge, this is one of the first metavirome works to unravel the Ganges viral habitat. The one of the major goals of present study is phage identification and to assess their diversity. Overall, an inspection and insights into abundance and diversity of phages, microbial community and ARGs from the Ganges River sediment was illustrated.

## Materials and Methods

### Sites and sampling

Sediment samples were collected from the major cities of the river Ganges i.e. from Bijnor, Narora, Kannauj, Kanpur, Allahabad, Allahabad Sangam, Mirzapur and Varanasi (**Figure S1 and Table S1**) in the month of June 2015. Sediment i.e. the upper layer of soil base, which is around 3-4 cm (∼2 inch) below the river water, was collected from these sites. Upstream and downstream locations were selected and GPS coordinates were monitored through GPS 72H by GARMIN (**Table S1**). All the samples were manually collected utilizing the properly decontaminated and clean stainless steel (SS) sampler. Samples were taken around 10-15 feet from the shore of the river and depth of water is around 5-8 feet. From each site, sediment was collected about 200-300 grams each. To avoid off-target contamination standard microbiological precautions were taken. We have collected sample in the aseptic and sterile glass bottles. Each time clean sampler and fresh gloves were used. Whirl pack bags were used for the sample collection. Further, transported to laboratory and stored at 4°C.

### DNA isolation

Sediment samples were processed for the DNA isolation using FastDNA spin kit (MP Biomedicals, LLC) following the manufacturer’s instructions. Briefly, 500 mg of sediment sample from each site is used to isolate DNA. After following the standard protocol and steps, binding matrix was gently resuspended into 50-100 µl of DES (DNase/pyrogen free water). Next, eluted DNA after centrifugation was stored at 4°C into the clean catch tubes. Finally, 50 µl of eluted DNA from each sample were pooled and concentrated in rotary vacuum evaporator up to half of its total volume. Next, agarose gel electrophoresis and Nanodrop-UV spectrophotometer were used for the quantity and purity check. Quantity of the pooled DNA sample was about 45ng/uL.

### Library preparation and Illumina sequencing

Illumina library was prepared using the NEXTFlex DNA library protocol and preparation guide at the Genotypic Technology’s Genomics facility. Very briefly, S220 system from Covaris Inc., USA was used to shear DNA and generate ∼200-500 bp fragments. Further, Agilent Bioanalyzer was utilized to see fragment size distribution. Next, HighPrep beads (MagBio Genomics, Inc, USA) used to clean fragmented DNA and further end-repair, A-tailing and adaptors ligation was performed through NEXTFlex DNA Sequencing kit following the manufacturer’s protocol. Adaptor ligated fragments were cleaned and subjected to PCR using primers from NEXTFlex DNA Sequencing kit. Qubit fluorometer and the Agilent Bioanalyzer were used for the quantification and size distribution of the prepared library respectively according to the manufacturer’s protocol. Finally, Illumina NextSeq sequencing platform is utilized for the sequencing of prepared library.

### Library preparation and Nanopore sequencing

DNA fragments of size >5kb was extracted from low melting point agarose gel. Gel purified DNA (∼300 ng, Qubit, Invitrogen) was used for the library preparation. Further, barcoded using PCR reaction (LongAmpTaq 2x New England Biolabs, USA), then 1.2x AmPure beads (Beckmann-Coulter, USA) was used to clean, and end-repaired through NEBnext ultra II end repair kit (New England Biolabs, USA). Next, 1x AmPure beads used to clean end-repaired DNA. Then, NEB blunt/TA ligase (New England Biolabs, USA) utilized to perform adapter ligations (HPA and HPT) for 15 minutes each. Further, 50 ul of MyOneTM streptavidin C1 beads (Invitrogen, USA) was used to clean library mix and eluted in 25ul of elution buffer. In total, ∼198 ng of sequencing library was used for the sequencing employing MinION Mk1b (Oxford Nanopore Technologies, UK) with SpotON flow cell ID FAB49007 (R9.4) on MinKNOW 1.1.21 in a 48hr sequencing protocol. Metrichor V.2.43.1 was used for the base calling.

### Metagenomic sequence data processing and analysis for phage identification

Illumina sequencing data from the Ganges sediment was subjected to the quality control to remove low quality reads (quality score<30) and filtering using NGSQC toolkit [58]. Correspondingly, metagenomic-sequencing data (FAST5) from Nanopore (MinION) sequencing platform was converted and extracted into more readable and useful FASTA and FASTQ formats through implementing poretools [59]. Finally, all quality filtered data from Illumina and 2D read dataset from Nanopore was utilized for the viruses/phage profiling and further analysis.

For this, a local database (VIRdb) was constructed with lineage information by utilizing all viral/phage reference sequences (9594). For the identification of probable viral reads and to reduce the search space, first BWA software package [60] with MEM algorithm is employed for illumina data (with default settings) and with predefined (-x) ont2d setting for the nanopore sequencing data. For same, viral reference sequence database (VIRdb) was indexed using bwa index function. Further, probable viral mapped reads from both the sequencing data were extracted from sequence alignment map (SAM)/binary alignment map (BAM) files using the SAMtools view function [61]. For virome classification and distribution analysis, probable viral reads were subjected to the blastn mapping [62, 63] with the stringent measures of 90 percent identity and 50 percent coverage constrain for the illumina data and for the nanopore data 70 percent identity and 50 percent coverage criteria was used with max target 1 (best hit only) to avoid the chance of getting false positive hits or to gain more confident virome distribution. The complete workflow used for the virome profiling and analysis in the study is illustrated in **Figure 1**. Virome distribution was shown using the krona plot [64] for interactive visualization.

**Figure 1.**
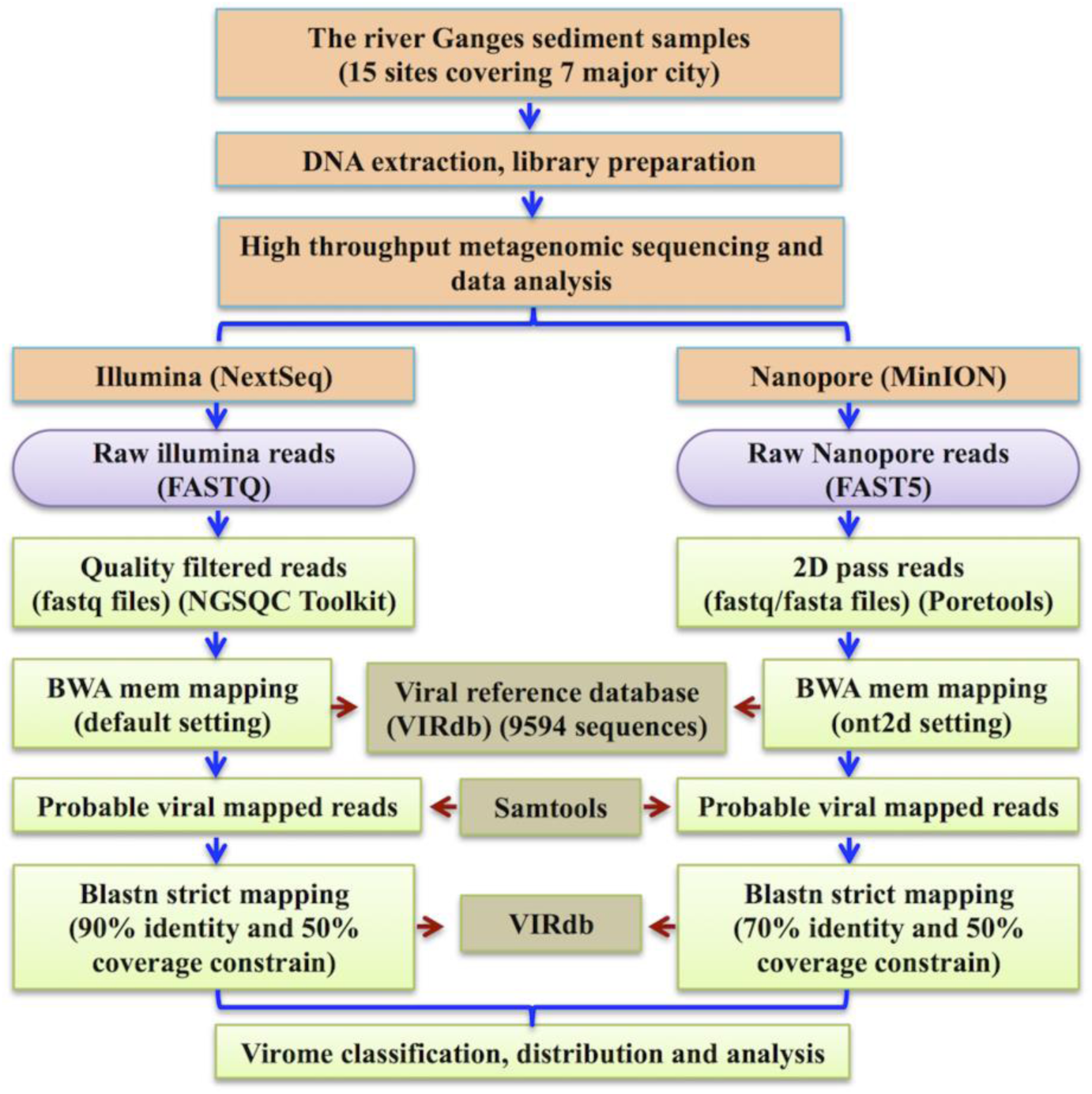
Analysis workflow for the metagenomic identification of viruses/phages

### Microbiome profiling and diversity from illumina metagenomic data

In order to identify and quantify the microbial community in the Ganges river sediment, MetaPhlAn2; a taxon-specific marker gene database was utilized [65]. The abundance at the different levels i.e. species, genus, family, order, class, phyla was reported. GraPhlAn software is used for the compact circular representation of the taxonomic structure [66].

### Illumina metagenomic analysis using the Structured ARG (SARG) database

For the analysis of ARGs from the illumina metagenomic data the Structured ARG database (SARG) was utilized to investigate the occurrence and abundance of ARGs using ARG-OAP [67], an online analysis pipeline, which combines Usearch [68] (ublastx_stage_one) and Blastx (ublastx_stage_two on Galaxy system) for ARG detection. SARG integrates ARG sequences from two most commonly used reference databases i.e. Antibiotic Resistance Genes Database (ARDB) [69] and the Comprehensive Antibiotic Resistance Database (CARD) [70] that are in a well-organized structure (ARG types and subtypes). To utilize ARG-OAP, firstly, an integrated database of ARGs (SARG.udb) was constructed using ARDB (7827 sequences) and CARD (6640 sequences; protein homolog model (4258), protein variant model (192) and protein wild type model (2190) after removing redundancies.

For ublastx_stage_one analysis, quality-filtered data (clean metagenomic sequences) were partitioned into smaller files (63 files for both the pairs, forward and reverse) using split command with 10000000 lines each due to memory constrain (i.e., 4 Gb limit) and limitation. Subsequently, each pair files were used for Ublastx_stage_one analysis to extract potential ARG reads from data. After this, two files (extracted.fa and meta_data_online.txt) were used for the stage two analyses. Extracted.fa contains potential ARG reads and Metafile include estimated detailed statistics.

In ublastx_stage_two analysis, extracted potential ARG-like sequences from stage one analysis were used for the on-line processing (ARG-OAP) on galaxy web server. Identification, annotation and classification of ARG-like sequences have been performed. A sequence fragment is identified as an ARG-like sequence, if it passes the stringent criteria of sequence identity (≥80%), E-value cut-off (1e-07) and alignment length (≥25 aa). Additionally, abundance of each identified ARG type and subtype were calculated.

ARG abundance was calculated using the portion of type or subtypes of ARG-like sequences in total metagenomic sequences. To avoid the biasness, ARGs abundance in the data can be normalized to the ARG reference sequence length and number of 16S rRNA genes using the equation as also described in previous studies [67] i.e.

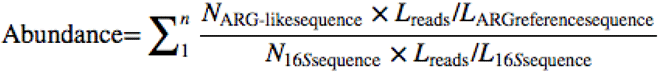

Where, N_ARG-like sequence_ denotes number of ARGs like sequences; L_reads_ define the length of the reads; L_ARG reference sequence_ represents nucleotide sequence length of the corresponding ARG reference sequence; N_16S sequence_ depicts number of the 16S rRNA gene sequences and L_16S sequence_ is the full length of 16S rRNA gene.

The complete approach used for the resistome profiling and analysis is represented in **Figure 2**. Relative abundance and distribution of different ARG types (classes) and subtypes (resistance genes) were depicted using various graphs and Krona tool [64] for interactive visualization.

**Figure 2.**
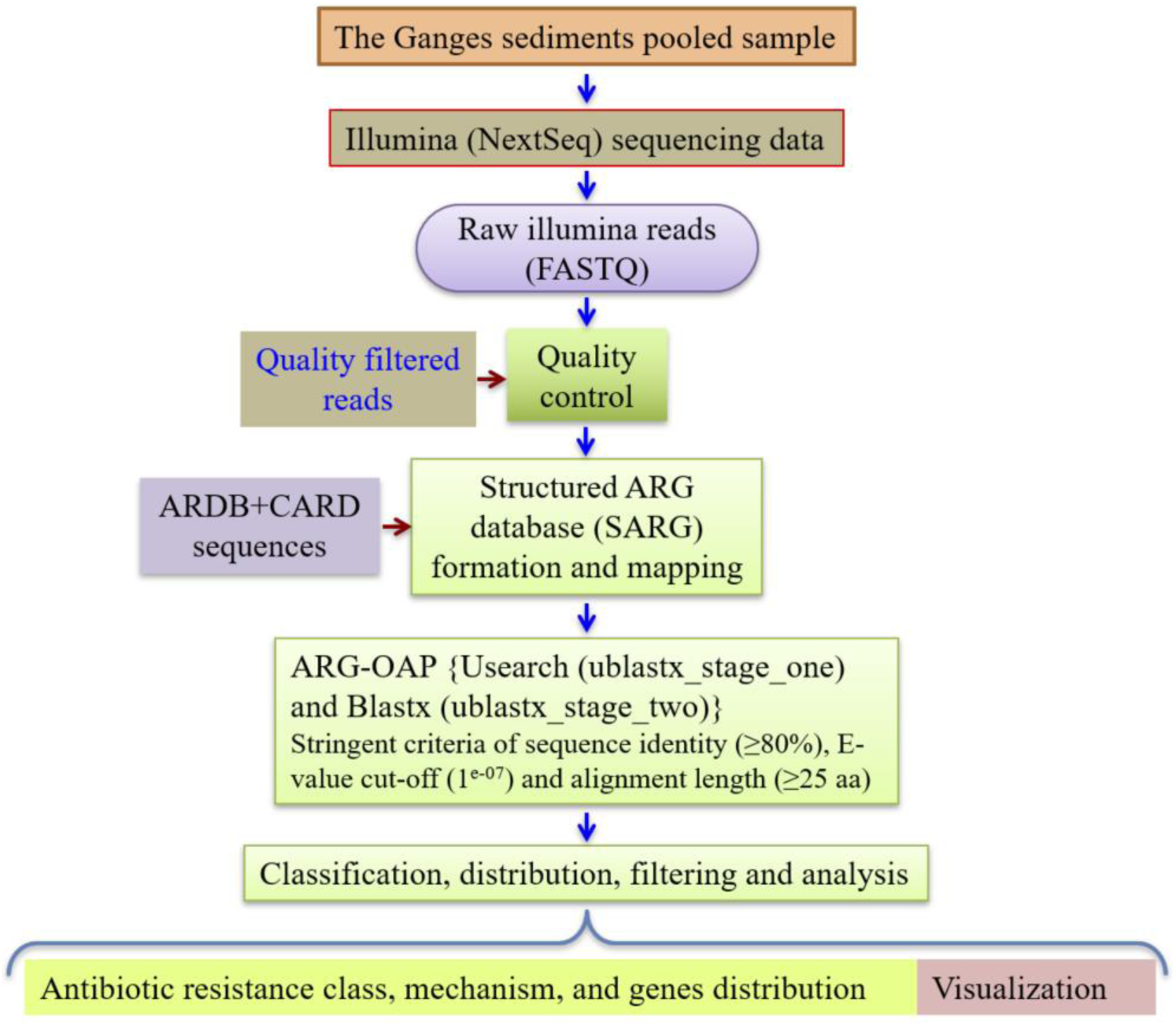
Workflow for resistome profiling

### ARG resistance mechanisms distribution

We have also analyzed the different resistance mechanism categories of the identified ARG subtypes. These categories were defined and classified according to the CARD database [70] and literature information. Abundance of each mechanism category was reported based on the collective occurrences of individual ARG subtypes belonging to particular class.

### Accession Number

Illumina and nanopore sequencing raw data are submitted in European Nucleotide Archive (ENA) with the accession number ERP109128.

## Results

### Illumina metagenomic sequencing and quality control

The Ganges sediment sample is processed with the illumine NextSeq sequencing platform and quality control and filtering of raw data is performed using NGSQC toolkit. All low-quality reads (Q<30) were removed. Also, adapter and primer trimming were performed. Overall, high quality filtered reads (314887862) were further used for the virome/phage exploration. Besides, profiling of microbial taxa and antibiotic resistance genes (ARGs) is also performed.

### Phage distribution and diversity

We have uncovered the distribution and abundance of different phages through Illumina data analysis. Inclusively, 4174 (∼0.0013%) reads were categorized for 285 distinct viruses from the Ganges river sediments (**Figure 3, Table S2**). Mainly, the presence of double-stranded DNA (dsDNA) viruses (4039), which account for approx. 97% of the total identified viral assemblages, is notified. Additionally, unclassified bacterial viruses, retro-transcribing viruses, ssDNA viruses, ssRNA viruses, dsRNA viruses and satellites were also found. Among all, dominance of distinct phages (3602, ∼86.30%) is discovered and analysis reveals the preponderance of order *Caudovirales* (dsDNA group I bacteriophages) that account for ∼85.82% (3582) of total viral habitat (**Figure S2**). The abundance profile of the order *Caudovirales* further described at the level of families i.e. *Myoviridae* (2347, ∼56.23%) followed by *Siphoviridae* (1139, ∼27.3%) and at last *Podoviridae* (88, ∼2.11%) (**Table S2**). Additionally, viruses from the family *Baculoviridae* (390, 9.34%) and *Poxviridae* (108, 2.59%) are also found in abundance (**Figure 4 and Table S2**).

**Figure 3.**
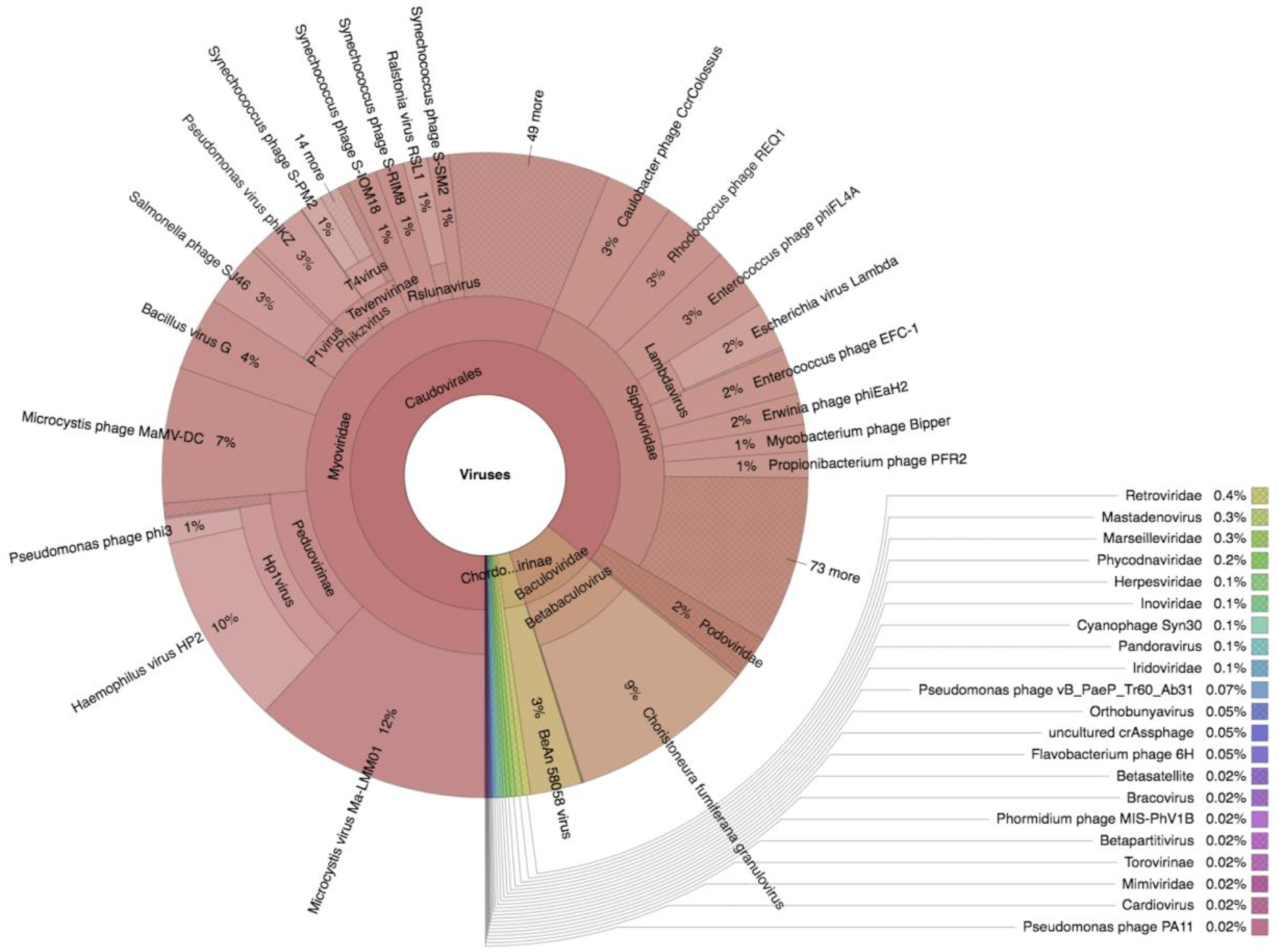
Krona plot showing virome spectrum identified from the Ganges river sediments using illumina sequencing

**Figure 4.**
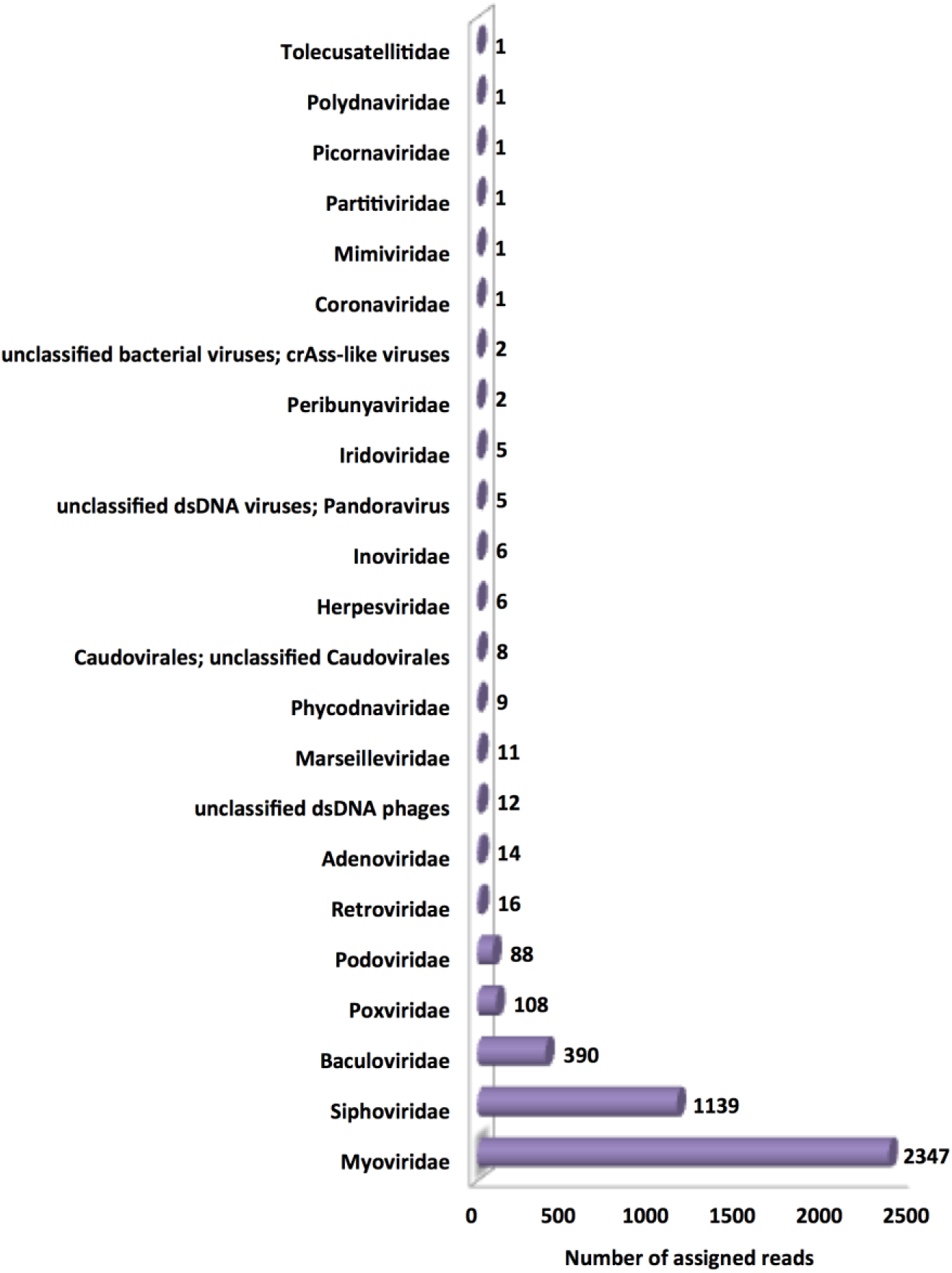
Family/group wise viral abundance profile identified from the Ganges river sediments employing illumina sequencing

Furthermore, we also present the abundance profile and existence of different bacteriophages at species level, which shows and recognized to have evident antibacterial pursuit against the pathogenic and deadly bacteria. Some of the most abundantly designated and found are *Microcystis phage Ma-LMM01* (498, ∼11.93%), *Haemophilus phage HP2* (403, ∼9.66%), *Microcystis phage MaMV-DC* (284, ∼6.8%), *Bacillus phage G* (155, ∼3.71%), *Salmonella phage SJ46* (130, ∼3.11%), *Pseudomonas phage phiKZ* (120, ∼2.87%), *Synechococcus phage S-IOM18* (61, ∼1.46%), *Synechococcus phage S-RIM8 A.HR1* (60, ∼1.44%), *Pseudomonas phage phi3* (52, ∼1.25%), *Ralstonia phage RSL1* (50, ∼1.2%), *Synechococcus phage S-PM2* (46, ∼1.1%) and *Synechococcus phage S-SM2* (45, ∼1.08%) members of the *Myoviridae* family; likewise, *Caulobacter phage CcrColossus* (143, ∼3.43%), *Rhodococcus phage REQ1* (140, ∼3.35%), *Enterococcus phage phiFL4A* (130, ∼3.11%), *Enterobacteria phage lambda* (99, ∼2.37%), *Enterococcus phage EFC-1* (95, ∼2.28%), *Erwinia phage phiEaH2* (65, ∼1.56%), *Mycobacterium phage Bipper* (56, ∼1.34%), *Propionibacterium phage PFR2* (55, ∼1.32%) belongs to the family *Siphoviridae*. Correspondingly, *Choristoneura fumiferana granulovirus* (386, ∼9.25%) member of family *Baculoviridae* and *BeAn58058 virus* (108, ∼2.59%) from *Poxviridae* family is also identified (**Figure 5 and Table S2**). **Figure 5** depicts the relatively abundant viral/phage species according to designated reads.

**Figure 5.**
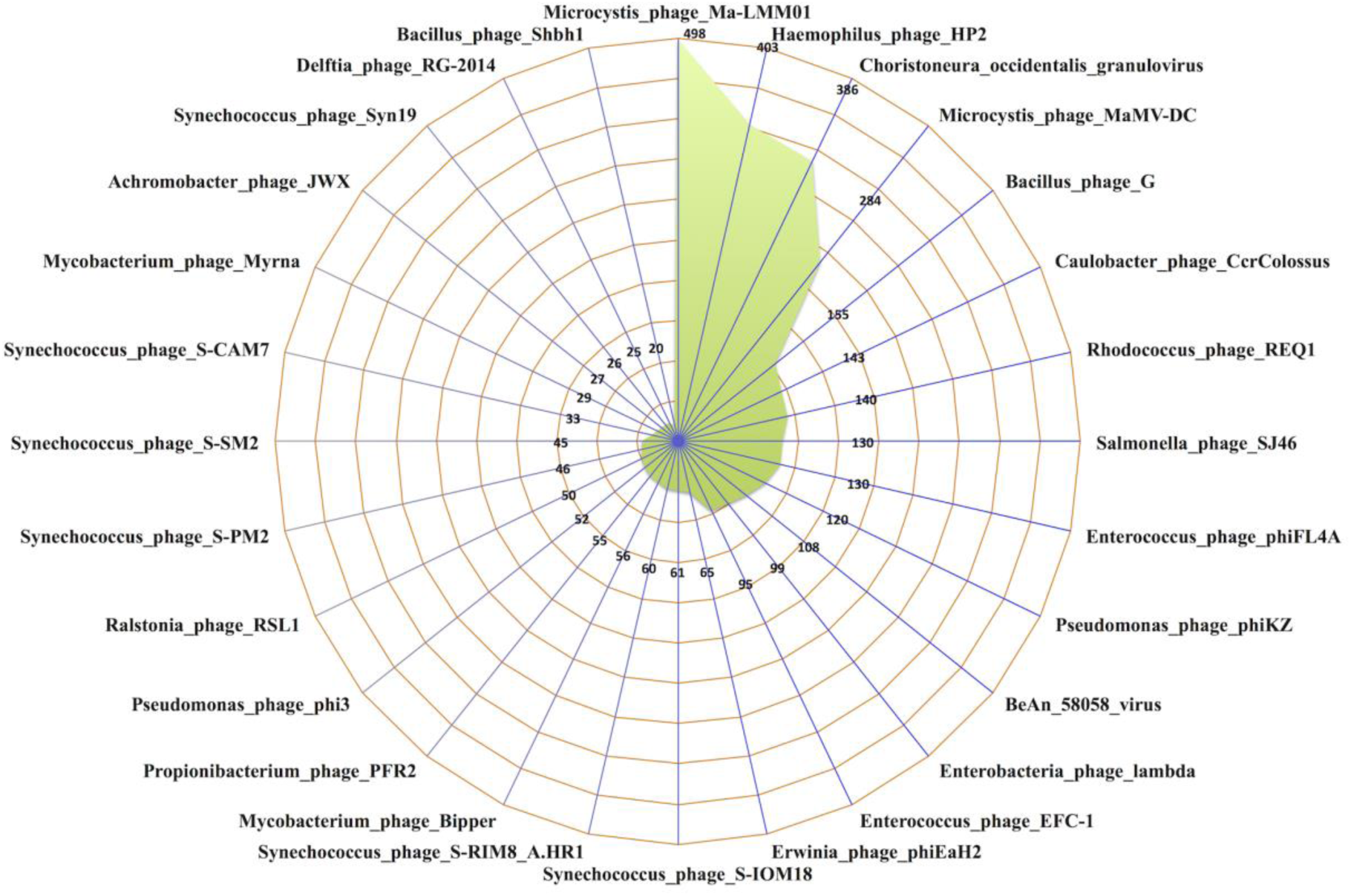
Radar graph depicting the 28 most abundant (≥20 reads) viral/phage species with number of designated reads

Among all, phages from different bacterial group/genus, i.e., *Microcystis* (782 hits, 21.71%), *Haemophilus* (403, 11.19%), *Synechococcus* (386, 10.72%), *Pseudomonas* (279, 7.75%), *Enterococcus* (232, 6.44%), *Bacillus* (196, 5.44%), *Rhodococcus* (166, 4.61%), *Caulobacter* (163, 4.53%), *Salmonella* (146, 4.05%), *Enterobacteria* (143, 3.97%), *Mycobacterium* (128, 3.55%), *Propionibacterium* (71, 1.97%), *Erwinia* (70, 1.94%), *Ralstonia* (56, 1.55%) shows the highest abundance and account for approximately 90% of the total identified phages (**Table S3**).

### Microbial community structure in Ganges sediment

Relative taxonomic abundance shows the presence of bacteria (∼94.4%) and archaea (∼5.6%) in the Ganges sediments. Overall, 9 phyla (7 bacterial and 2 archaeal), 17 classes (14+3), 38 orders (33+5), 72 families (65+7), 119 genera (107+12), 131 species (117+14) were revealed (**Table S4**). The relative abundance and estimated read distribution of microbial taxa are provided in the **Table S4**. *Proteobacteria* (∼69.3%) and *Actinobacteria* (∼15%) are primarily dominating the microbial community structure. Mainly, orders i.e. *Caulobacterales* (∼28%), *Burkholderiales* (∼13.9%), *Actinomycetales* (∼13.7%), *Pseudomonadales* (∼7.5%), *Rhodocyclales* (∼7%) form around 70% of the microbiota and the families i.e. *Caulobacteraceae, Oxalobacteraceae, Micrococcaceae, Rhodocyclaceae, Pseudomonadaceae* represent the major component of microbiome of the Ganges sediment (**Table S4**). Taxonomic structure is represented using circular graph with relative abundance and reads distribution (**Figure 6**).

**Figure 6.**
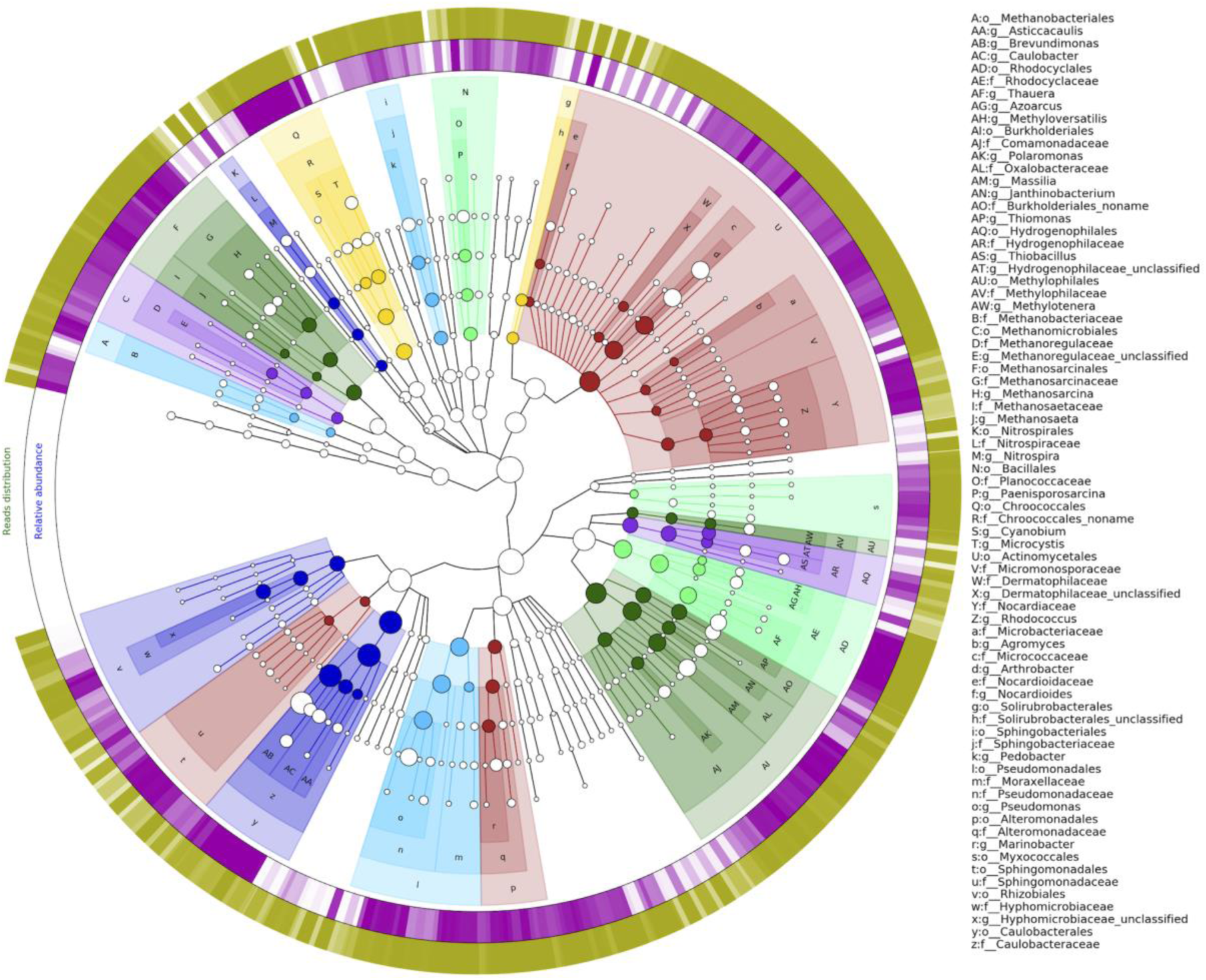
A taxonomy tree representing microbial ecology from the sediment of the Ganges. Relatively abundant taxa (>0.5%) at different levels (orders, families and genus) were marked in different abbreviated form. Outer most green ring represents the read distribution and purple ring shows relative abundance and color intensity signify the relative percentage.

### Existence and abundance of antibiotic resistance genes (ARGs)

Furthermore, occurrence and distribution of ARGs is evaluated from the Ganges sediment. ARG analysis utilizing Structured ARG (SARG) database revealed abundance (the portion of ARG-like sequences in total metagenomic sequences, denoted using the unit -ppm, i.e. one read in one million reads) profile of different ARG types and subtypes. Overall, 21869 (∼0.00695%) illumina reads were categorized as ARG (**Figure 7 and Table S5**). In total, 20 ARG types (classes) and 240 ARG subtypes (genes) were detected (**Tables S5 and S6**).

**Figure 7.**
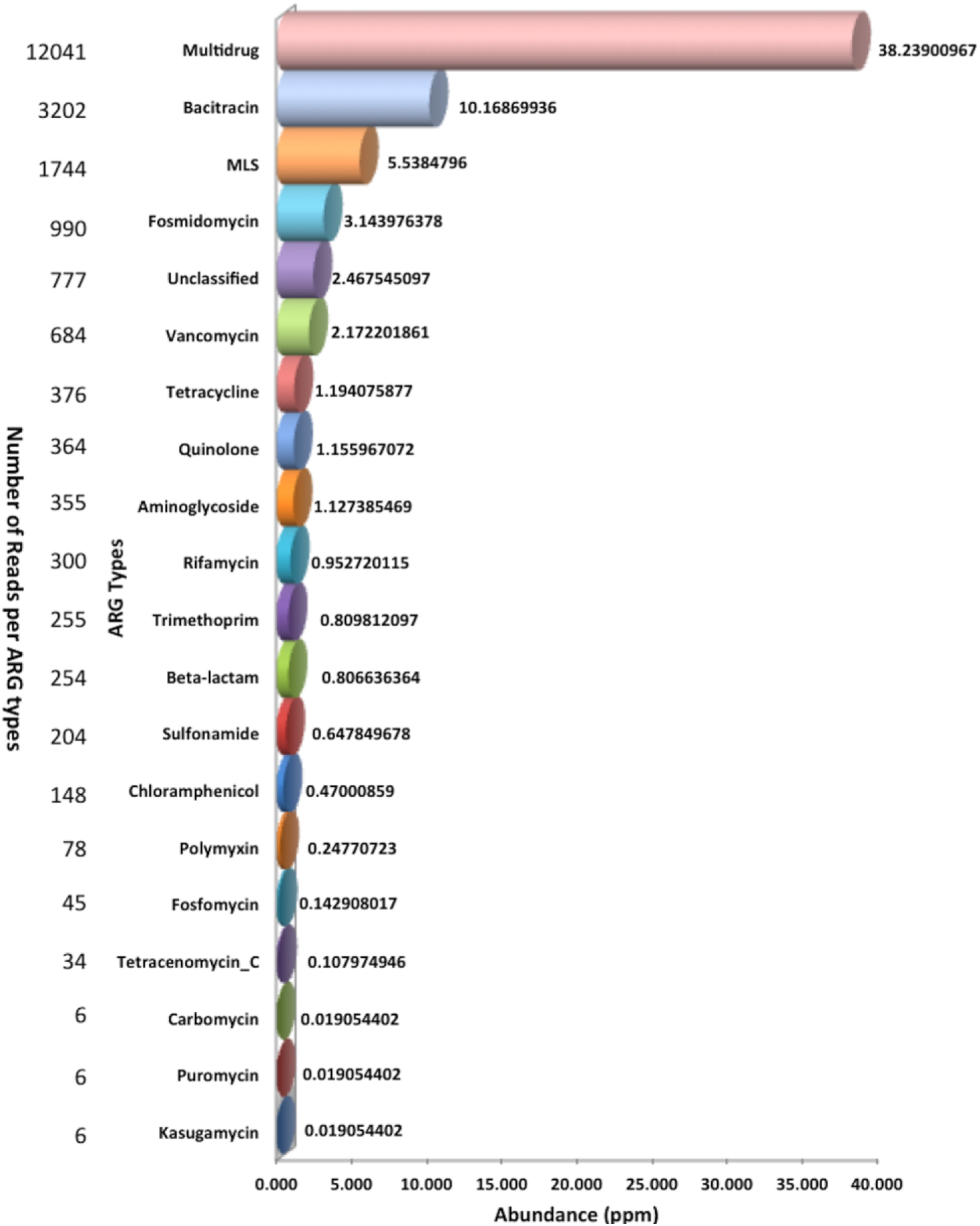
Distribution of ARG types in the Ganges river sediment sample

### ARG types and subtypes

ARG type distribution in the sample is wide-ranging from 38.24 ppm (Multidrug resistance genes) to 0.01 ppm (puromycin and carbomycin resistance genes) (**Figure 7**). The most abundant ARG types are multidrug (MDR) (12041 reads, ∼55 %, ∼38.24 ppm), bacitracin (3202 reads, 15%, ∼10.17 ppm), macrolide-lincosamide-streptogramin (MLS) (1744 reads, 7.98 %, ∼5.54 ppm), and fosmidomycin (990 reads, 4.53%, ∼3.14 ppm) resistance genes compose around 82% of total ARG reads. Apart from most abundant ARG types unclassified, vancomycin, tetracycline, quinolone and aminoglycoside are also rich in distribution (≥1 ppm) than the other ARG types approximately account for another ∼12% of total ARG reads. Likewise, different ARG subtypes (genes) are also assessed. *bacA* gene that confer resistance to bacitracin (3145, ∼14.38%), MDR gene *mexF* (2441, ∼11.16%), *acrB* gene, a bacterial multidrug efflux transporter (1634, ∼7.5%), *macB* gene confer resistance to MLS class drugs (1421, ∼6.5%) and the multidrug_ABC_transporter (1067, ∼4.9%), and *mdtB* (968, ∼4.4%) belong to MDR are the most abundant ARGs in the Ganges sediment (**Table S7**). Complete catalogue of 240 ARGs with frequency is provided in **Table S7**.

MDR is the most abundant resistance type detected in the Ganges sediments sample (**Figure 7 and Table S5**). Overall, 59 unique subtypes of MDR were detected which constitute around 55% of total ARG reads (**Table S6**). Among the 56 subtypes, the representative 38 MDR subtypes (Number of reads >10) were shown in **Figure S3**. Including the distribution profile of major subtypes (ppm>1) i.e. *mexF* (∼7.75 ppm, 2441 reads), *acrB* (∼5.19 ppm, 1634), multidrug_ABC_transporter (∼3.39 ppm, 1067), *mdtB* (∼3.07 ppm, 968), multidrug_transporter (∼2.70 ppm, 850), *mexW* (∼2.13 ppm, 671), *ceoB* (∼1.96 ppm, 616), *mdtC* (∼1.84 ppm, 579), *smeE* (∼1.08 ppm, 341) and so on. Likewise, bacitracin (∼10.17 ppm, 3202 reads) is the second most abundant ARG type (**Figure 7 and Table S5**). Overall, it creates 14.65% of entire ARG types approximately. There are two subtypes, i.e., *bacA* and *bcrA* were found in this category. *bacA* (∼9.99 ppm, 3145 reads) is having largest proportion among all subtypes. It makes around 14.4% of all ARG subtypes and is most abundant ARG subtype detected in the study (**Table S7**). Additionally, *bcrA* (∼0.18 ppm, 57) is also discovered. MLS is the third most abundant ARG type (∼5.54 ppm, 1744 reads) (**Figure 7 and Table S5**). In total, 28 subtypes were detected that account for around 8% of total ARGs. *macB* (∼4.5 ppm, 1421 reads) is the most prominent, which make around 83 percent of total MLS identified. Simultaneously, fosmidomycin (∼3.14 ppm) were also revealed. In total 990 reads were recorded, which account for approx. 4.52% of all ARG types (**Figure 7 and Table S5**). There are two subtypes detected in this type i.e. *rosA* and *rosB. rosB* (∼ 2.51 ppm) is more abundant subtype with 791 reads. Likewise, in *rosA* (∼0.63 ppm), 199 reads were categorized (**Table S6**).

Apart from above mentioned top most abundant ARG types, vancomycin resistance genes (VRGs) were also found in relevant amount. Overall, 13 unique subtypes of vancomycin were identified. Among these, *vanR* (∼1.42 ppm) is the most abundant VRG with 447 reads (65% of VRGs). Likewise, second abundant VRG is *vanS* (∼0.49 ppm), account for 23% of all VRGs and comprises of 154 reads (**Table S6**). Tetracycline resistance genes (TRGs) are also one of the important classes of ARGs. We have identified 376 reads pertaining to 20 TRG subtypes constitute around 1.72% of total ARG reads (**Table S6**). *tetP* (∼0.44 ppm), *otrA* (∼0.18 ppm), *tetM* (∼0.11 ppm) and *tet35* (∼0.11 ppm) are most abundant TRGs with 139, 58, 36 and 35 reads, respectively (**Table S6**). Another class of ARGs i.e. quinolone (∼1.16 ppm) is also discovered. There are 364 reads quantified in this category, which make approximately 1.65 percent of total ARG reads. There are three subtypes were identified namely *norB, qepA* and *qnrS*. Among these *qepA* (∼0.60 ppm) is most abundant with 190 reads. Likewise, 172 reads were demarcated for *norB* (∼0.55 ppm) and least 2 read for *qnrS* (**Table S6**). Aminoglycoside (∼1.13 ppm) resistance is also revealed that consists of 355 reads that is around 1.62% of total ARGs. In total, 22 unique aminoglycoside subtypes were uncovered (**Table S6**). Among all, *aadA* has the highest abundance (∼0.31 ppm) with 98 reads. Beta-lactam resistance genes (BRG) are also one of the key classes of ARGs. We have identified 254 reads corresponds to 59 subtypes represent around 1.17% of total ARG reads (**Tables S6**). Among all, *THIN-B* (∼0.07 ppm), *OXA-209* (∼0.05 ppm) and *OXA-36* (∼0.04 ppm) are most abundant BRGs with 24, 16 and 13 reads, respectively. Additionally, some ARGs are categorized in unclassified category. We have found 4 subtypes in this class i.e. transcriptional regulatory protein *CpxR* (∼1.48 ppm, 459 reads), putative response regulator *ArlR* (∼0.69 ppm, 218), cAMP-regulatory protein (∼0.18 ppm, 57 reads), and histidine kinase (∼0.13 ppm, 43) (**Table S6**).

### ARG resistance mechanisms

Different resistance mechanisms from the Ganges sediments were identified. Overall, 238 ARGs were categorized in the six different mechanisms based on the CARD database and literatures i.e. 99 subtypes from antibiotic inactivation, 83 correspond to antibiotic efflux, 27 belongs to antibiotic target alteration, 14 from antibiotic target protection, 11 from antibiotic target replacement, 4 subtypes from reduced permeability to antibiotic (**Table S8**). Two subtypes are unclassified and specified as “others”. Notably, dominance of resistance mechanism is differing in terms of ARG abundance as compare to subtype numbers. The most dominant mechanisms are antibiotic efflux (72.8%, 15919 reads), antibiotic target alteration (18.3%, 4006) and antibiotic inactivation (5.1%, 1123) according to relative abundance of ARGs (**Figure S4**). Approximately 96% of the total ARG subtypes are covered under these three mechanisms.

### Nanopore metagenomic sequencing data statistics and phage profiling

We have also made attempt for phage profiling using nanopore sequencing technology. Raw nanopore reads (FAST5 files) were extracted into FASTA and FASTQ files using poretools. In total, 48605 reads from the Ganges sediment sample were utilized for the analysis. Maximum read size is 9273 bp and average read size is ∼1000 bp.

Virome profiling from Nanopore data reveals the presence of double-stranded DNA (dsDNA) viruses. Inclusively, 60 (∼0.12%) reads were assigned for the viruses from the Ganges river sediments (**Table S9**). Among all, the most abundant identified order is *Caudovirales*, which represent dsDNA group I bacteriophages that account for ∼98.33% (59 hits) of total designated viral habitat. The order *Caudovirales*, further categorized into two families named according to their percent share i.e. *Myoviridae* (46, ∼76.66%), followed by *Siphoviridae* (13 reads, ∼21.67% of total viral content). Additionally, read belong to *Poxviridae* family (1, ∼1.7%) is also found (**Table S9 and Figure S5**).

Further, analysis shows the existence of distinct bacteriophages namely *Pseudomonas phage OBP* (45 reads, ∼75%), *Microcystis phage Ma-LMM01* (1, ∼1.7%) belong to *Myoviridae* family, and *Enterobacteria phage lambda* (*Escherichia virus lambda*) (9, ∼15%), *Pseudomonas phage B3* (1), *Caulobacter phage* (1, ∼1.7%), *Burkholderia phage phiE125* (1, ∼1.7%), and *Erwinia phage PhiEaH1* (1, ∼1.7%) members of the family *Siphoviridae*, which are known to have certain bactericidal activity against respective pathogens. Correspondingly, *BeAn 58058 virus* (1, ∼1.7%) from *Poxviridae* family is also identified (**Figure S5** and **Table S9**).

Moreover, we have also explored the identified genomic regions of these phages to characterize and identify the particular gene products. Mainly from the *Pseudomonas* phage OBP, occurrence of putative virion structural proteins, hypothetical proteins, T4-like DNA polymerase, putative virion-associated RNA polymerase beta’ subunit, putative chaperonin GroEL, putative UvsX and putative terminase large subunit was observed. Likewise, tail component, tail:host specificity protein, *NinE* protein, *NinI* protein, *xis* (*Excisionase*) and integration protein from *Enterobacteria phage Lambda* was identified. Similarly, presence of some gene products i.e. portal protein (*Pseudomonas phage B3*), *tnpB* (*Burkholderia phage phiE125*), putative RtcB-like protein (*Caulobacter phage*), phoH (*Erwinia phage PhiEaH1*) and Kelch repeat and BTB domain-containing protein (*BeAn 58058 virus*) were also marked. These diverse sets of phages could be utilized and considered for the development of combat strategies against disease causing bacteria.

## Discussion

For the first time, in 1896 British bacteriologist Ernest Hankin demonstrated the antibacterial property of the Ganges water against the bacteria *Vibrio cholera*. Approximately after two decades (1915), Frederick William Twort illustrated that viruses might be the factor behind it. Independently, in 1917, Felix d’Herelle a microbiologist at Pasteur Institute particularized and discovered the phages as agent and proposed the term “bacteriophage”, Greek meaning “bacteria eater” [71-73]. The Ganges River is known and well regarded for its healing properties. It becomes lifeline for millions of people in India. Bacteriophages were discovered and introduced to the world through studies on the Ganges River [72, 74].

Viruses including phages (viral parasites of bacteria) are the most abundant biological bodies on the earth ecosystem. Phages play an important role and have wide applications from medical sciences to food safety [75, 76]. Bacteriophages are known for their highly specific bactericidal activity without influencing the natural flora. This ability of phages makes them as prospective tool to treat diseases. They are also known to effect and shape structure of bacterial community [10]. Applications of bacteriophages or phage cocktails for the treatment of multiple diseases like diabetic foot ulcer, leg ulcer, chronic otitis and burn wound infection are shown in various studies and clinical trials [75, 77-79]. Also, prompt escalation in the antibiotic resistance bacteria has recommenced focus on the phage therapy [76].

Antibiotic resistance is a global burden and major challenge towards the public health security. Diverse environments such as human gut microbiota, human feces, animal waste, hospital wastewater, soil, sewage, activated sludge, wastewater treatment plants (WWTPs), glaciers, rivers, sediments etc. are known to have and considered to be a reservoir of ARGs[35, 80, 81]. Recently, Department of Biotechnology (DBT), Government of India and the Research Council UK (RCUK) commissioned report “*Scoping Report on Anti-microbial Resistance in India*” compiled the antimicrobial Resistance (AMR) and microbial studies from India and also shade light on the threat of AMR, its causes and sources prepared by the Centre for Disease Dynamics and Economic Policy (CDDEP). This report also points out factors contributing to AMR i.e. mass-bathing in the Ganges during pilgrimages, inappropriate use of antibiotics (Humans, Animals), untreated wastewater effluent discharge from pharmaceutical industry etc. This report also suggests improving the waste handling and treatment facilities in pilgrimage area to better protect ARG dissemination. It further highlights that AMR research studies in India are lacking in all areas including environment despite of crucial challenges and recommended the urgent need to fill the gap through more research on different aspects of AMR from Humans, animals and environment [35].

There are various studies, which advocate the importance and focused on the exploration of virome and phages from diverse sources [3, 14, 82-87]. However, to best of our knowledge, there is no metagenomic study reported that explore phage diversity from the Ganges River since the discovery of phages. Nevertheless, very recently, a study described the abundance of microbial taxa and ARGs from the five riverbanks (Assi, Bhadaini, Harishchandra, Dr. Rajendra Prasad (Dr. RP) and Rajghat) of the Ganges at Varanasi city [88]. Reddy and Dubey have shown the dominance of *Proteobacteria* and *Actinobacteria* phyla in sediment. Also, specified most abundant ARG types i.e. Beta-lactam, multidrug and aminoglycoside in sediment [88]. Lately, Samson *et al.* described the influence of confluence of Yamuna River on bacterial taxonomy of the Ganges [89]. A study by Ahammad *et al.* describes the identification of ARGs from the upper Ganges and the Yamuna River. They analyzed the samples from seven sites in the Rishikesh-Haridwar region from the Ganges and five sites on the Yamuna River in Delhi associated with seasonal Pilgrimages in May/June. This study reports the presence and abundance of beta-lactam (i.e. *bla*NDM-1 and *bla*OXA-48) and tetracycline (i.e. *tet*M, *tet*W, *tet*Q) ARG types [90], which is also in coherence with the finding of this study. Another study Devarajan *et al.* provide the occurrence of ARGs mainly beta-lactam (*bla*NDM-1, *bla*SHV, *bla*CTX-M etc.) from the Cauvery River Basin (CRB), Tiruchirappalli, Tamil Nadu, India [50]. Likewise, Marathe *et al.* highlighted the effect of untreated wastewater discharge on the resistome from Mutha River, Pune, India. They have identified the 175 ARGs including the *bla*NDM-1, *bla*OXA-48, *tet*X, *bla*OXA-58, mcr-1 etc. using shotgun metagenomics approach [41]. Furthermore, study from Diwan *et al.* reported the presence of ARG types (beta-lactams, quinolone etc.) from seven regular and three mass-bathing sampling points from the river Kshipra, Ujjain, India [91]. These findings from different studies irrespective of distinct source rivers represent commonality and reported to have the presence of mainly beta-lactam and tetracycline resistance genes. Our study also elucidates the occurrence of these ARG classes with many others.

This study emphasizes on the metagenomic exploration of phages, microbial community and ARGs from the Ganges River sediments. We have collected the sediment samples from upstream and downstream locations of seven major cities along the middle stretch of the river Ganges i.e. from Bijnor, Narora, Kannauj, Kanpur, Allahabad, Allahabad Sangam, Mirzapur and Varanasi during the month of June 2015 and analyzed the pooled sediment sample to provide a considerably quick profiling of the Ganges habitat. However, further studies need to be done to explore spatial distribution at individual sites. We have mainly utilized Illumina (NextSeq) sequencing platform for the study and also made efforts to explore nanopore sequencing platform (MinION) for phage profiling. However, there is a requirement for improvement and optimization of nanopore sequencing and further analysis. Illumina metagenomic analysis shows the presence of different phages with the dominance of order *Caudovirales*. Our study shows the prevalence of *Myoviridae, Siphoviridae* and *Baculoviridae* viral families. Moreover, phages belong to the *Microcystis, Haemophilus, Synechococcus, Pseudomonas, Enterococcus, Bacillus, Rhodococcus, Caulobacter, Salmonella, Enterobacteria, Mycobacterium* group/genus are present in abundance. Also, we were able to find *Myoviridae* as most dominant family and existence of phages mainly for *Pseudomonas, Enterobacteria, Microcystis* utilizing nanopore sequencing. Furthermore, we show the relative abundance of microbial community that mainly dominated by the *Proteobacteria* and *Actinobacteria.* Additionally, *Cyanobacteria, Bacteroidetes, Firmicutes* are also found in abundance. Very recent report published by Reddy and Dubey also substantiates our findings [88]. Simultaneously, Archaeal phyla *Euryarchaeota* and *Thaumarchaeota* also identified and reported. Moreover, ARGs profile is also revealed, which comprise 20 distinct ARG types and 240 ARG subtypes. We found the rich abundance of MDR, bacitracin, MLS and fosmidomycin. Similarly, ARG subtypes *bacA, mexF, acrB, macB* are most abundant in the Ganges sediment. Correspondingly, antibiotic efflux, antibiotic target alteration and antibiotic inactivation are the most prevalent resistance mechanism identified. Overall, our work provides abundance and distribution of various phages, microbial taxa and ARGs from the Ganges sediment.

## Conclusion

To best of our knowledge, this is the first metagenomic report, which documents the phage profile from the sediment of the Ganges River. Our study shows the dominance of bacteriophages (i.e. order *Caudovirales* or different families like *Myoviridae and Siphoviridae*) in overall viral composition. Simultaneously, microbial community analysis and ARG profiling is also performed. Taxonomic analysis disclosed prevalence of distinct phyla i.e. *Proteobacteria, Actinobacteria, Cyanobacteria, Bacteroidetes* and *Firmicutes.* Additionally, ARG profiling shows dominance of MDR and bacitracin. This also specifies the need for effective management and treatment of waste (i.e. fecal, sewage, industrial, hospital etc.). This type of surveillance studies is imperative for sustainable improvement and efficiently developing control measures. Moreover, we also propose indubitably that such type of studies will assist in the exploration of highly diverse kingdom of viruses on earth, to monitor the environmental quality and to expand the size of viral sequence resources for further functional characterizations. There are also immense opportunities in the field of phage-based therapeutics and solutions to fight against disease conditions caused due to bacterial pathogens. Nonetheless, it is also necessary to explore specific viral or phage communities that uncovered from metagenomic studies through advancing the isolation methods, functional characterization and interpreting their role in distinctive situations.

## Author Contributions

Conceived and supervised the experimental study, G.S. and S.M.; Collections of samples, extraction of community DNA and processing for metagenomic sequencing, N.K., S.K.S. and D.P.; supervised the computational analysis, M.K.; computational NGS data analysis and interpretation, A.K.G. and M.K.; virome/phage and microbial community analysis, A.K.G. and M.K.; antibiotic resistance genes profiling, A.K.G., V.R. and M.K.; Manuscript writing, A.K.G., N.K., S.M. and M.K.

## Funding

This research work was supported by the Council of Scientific & Industrial Research (CSIR)-Institute of microbial technology (IMTECH); Ministry of Water Resources, Development & Ganga rejuvenation project entitled “Water quality monitoring of the Ganga river from Gomukh to Hooghly (National Mission for Clean Ganga–NMCG”, Grant number GAP-0147-20) and Department of biotechnology (DBT) (Grant number GAP0001).

### Acknowledgments

We appreciate technical help from Mr. Malkit Singh. S.K.S. and A.K.G. are supported by DST-INSPIRE fellowship sponsored by Department of Science & Technology (DST), Government of India. D.P. is supported by an UGC-JRF fellowship sponsored by University Grants Commission, Government of India. N.K. is supported by CSIR-SRF, Council of Scientific & Industrial Research (CSIR), Government of India. V.R. is supported by National Post-Doctoral Fellowship (N-PDF) funded by Science and Engineering Research Board (SERB), Department of Science & Technology (DST), Government of India.

## Supplementary Materials

Figure S1: Map showing sample collection sites on the Ganges river, Figure S2: Circular plot showing distribution of order Caudovirales with relative abundance uncovered from the Ganges sediments, Figure S3: Abundance profile and reads distribution of representative 38 MDR subtypes. Pie chart (inset) shows number of reads belongs to each subtype, Figure S4: Antibiotic resistance mechanism distribution. Red bar shows relative abundance (%) and blue portion depict number of subtypes belongs to particular mechanism, Figure S5: Circular plot depicting virus profile identified using Nanopore sequencing technology, Table S1: Sample collection sites on the Ganges river, Table S2: List of all the identified viruses/phages from the Ganges river sediments using the Illumina sequencing along with relative abundance percentage, Table S3: Relative distribution of reads belongs to identified phages from different bacterial group/genus, Table S4: Diversity and abundance of microbial community in the Ganges sediment, Table S5: Table showing number of reads and relative abundance assigned to 20 different ARG classes (types) discovered in the Ganges sediments, Table S6: Alphabetical list of all the identified ARG classes (types) and genes (subtypes) along with corresponding abundance and number of reads, Table S7: Abundance wise catalogue of antibiotic resistance genes, Table S8: Distribution and abundance of the antibiotic resistance mechanisms, Table S9: Identified viruses/phages from the river Ganges sediment employing Nanopore sequencing.

