## Supplementary Materials for "Abundance and diversity of phages, microbial taxa and antibiotic resistance genes in the sediments of the river Ganges through metagenomic approach"

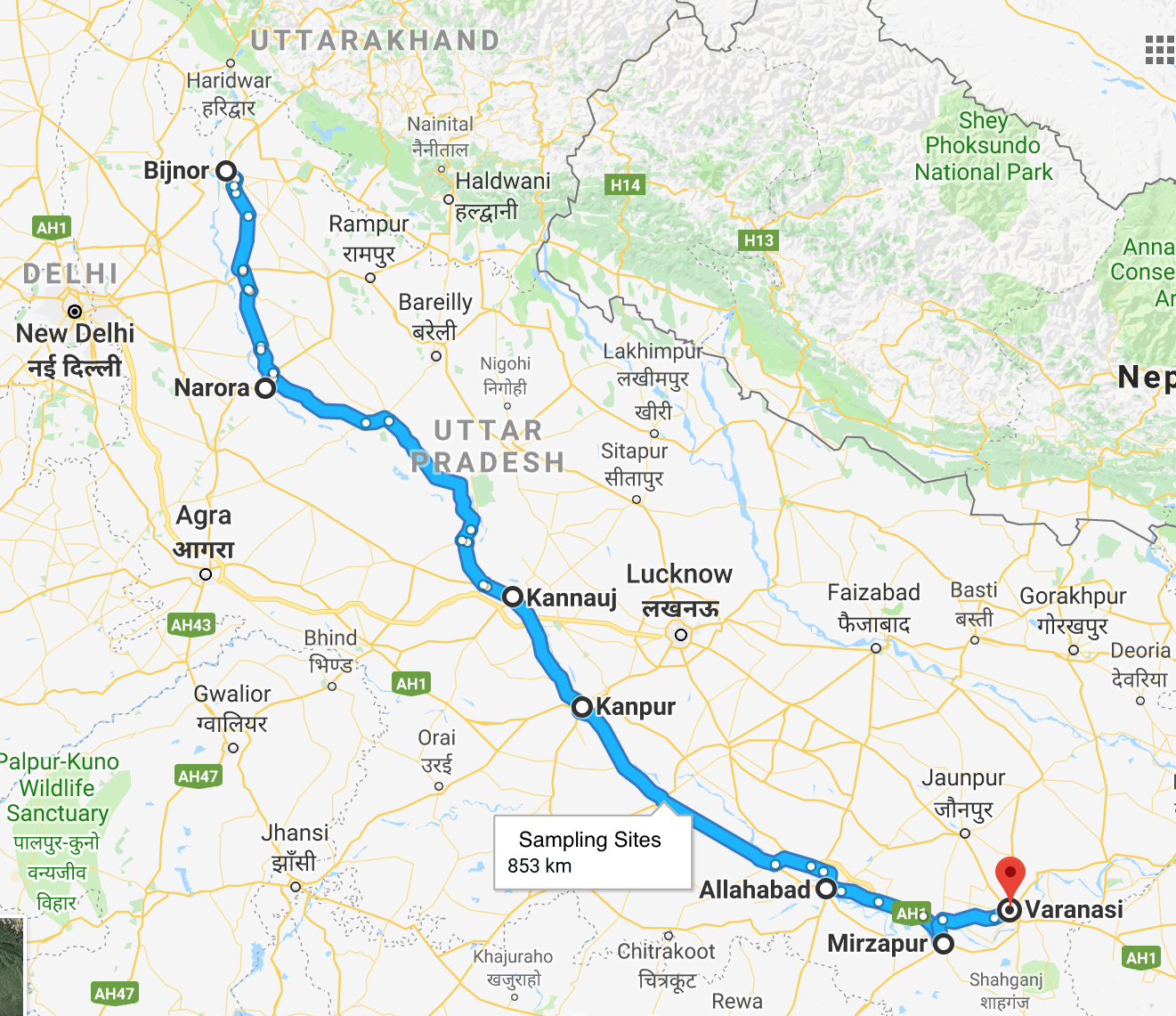

**Figure S1**. Map showing sample collection sites on the Ganges river

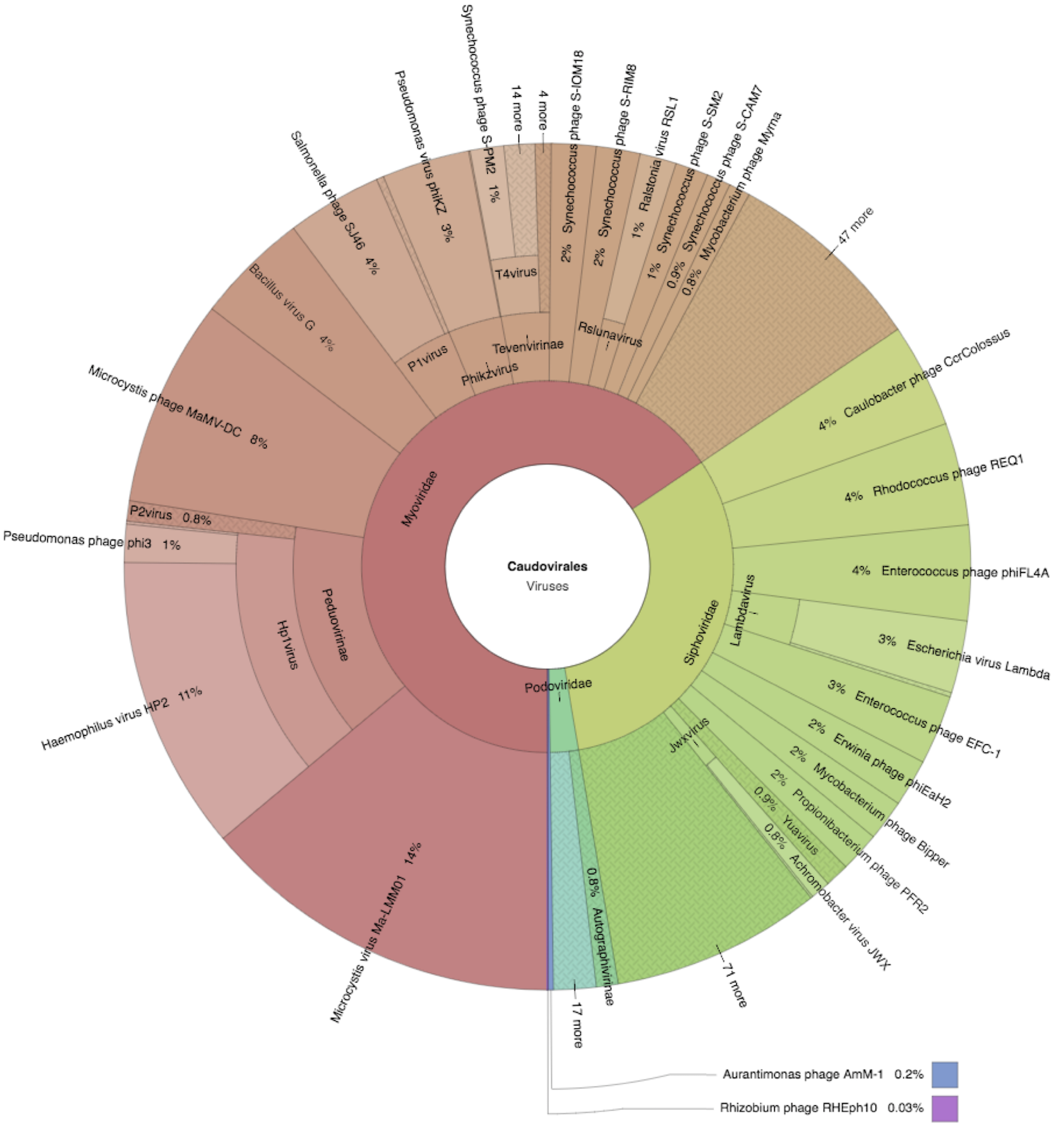

**Figure S2**. Circular plot showing distribution of order *Caudovirales* with relative abundance uncovered from the Ganges sediments

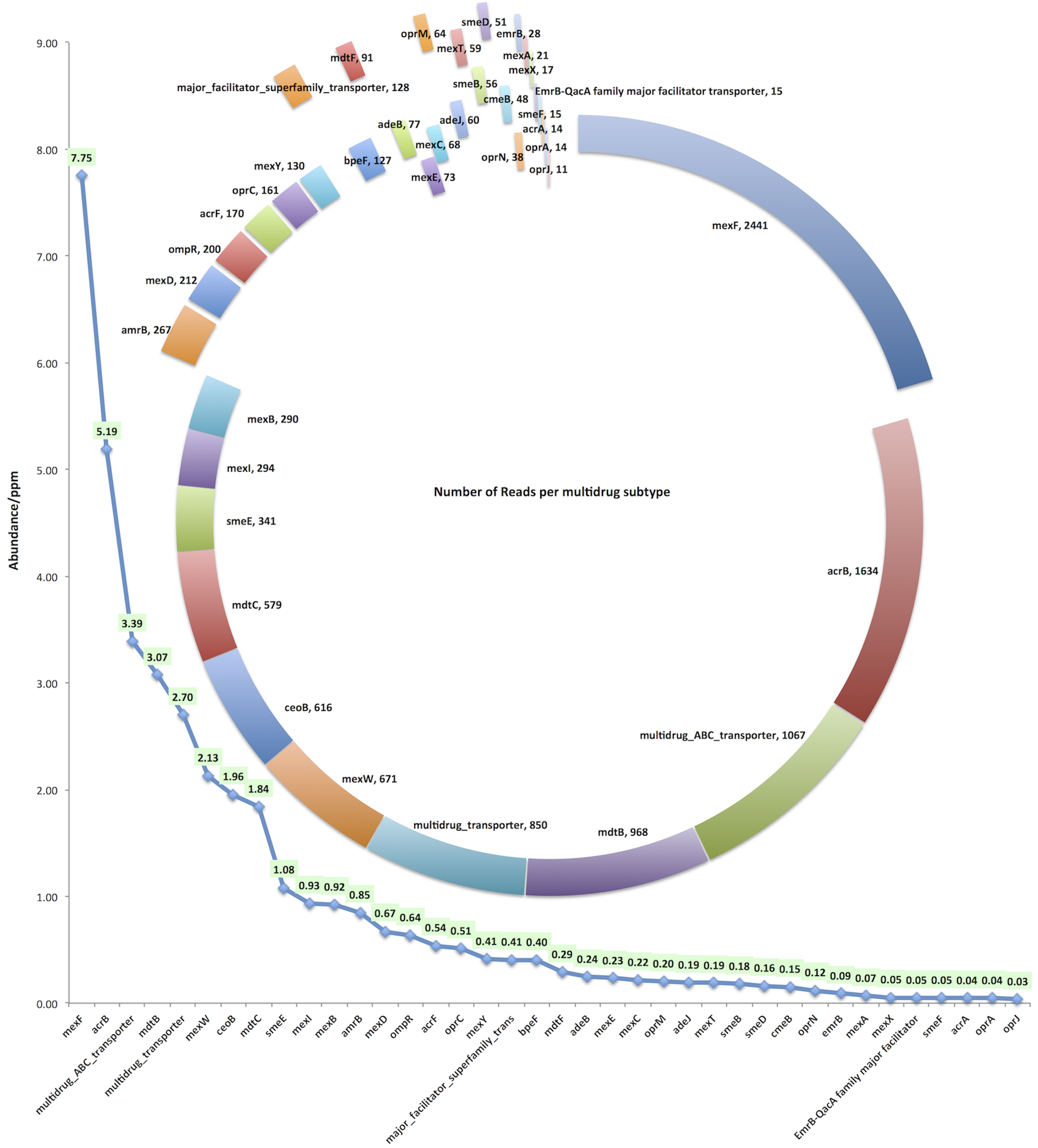

**Figure S3**. Abundance profile and reads distribution of representative 38 MDR subtypes. Pie chart (inset) shows number of reads belongs to each subtype

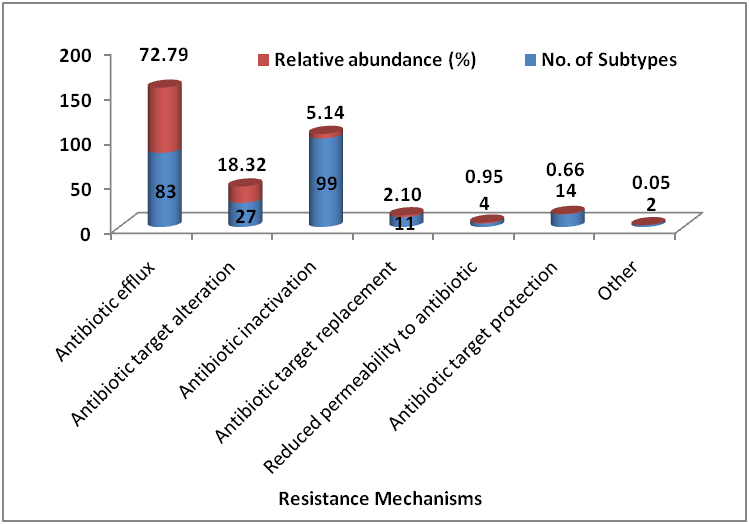

**Figure S4**. Antibiotic resistance mechanism distribution. Red bar shows relative abundance (%) and blue portion depict number of subtypes belongs to particular mechanism.

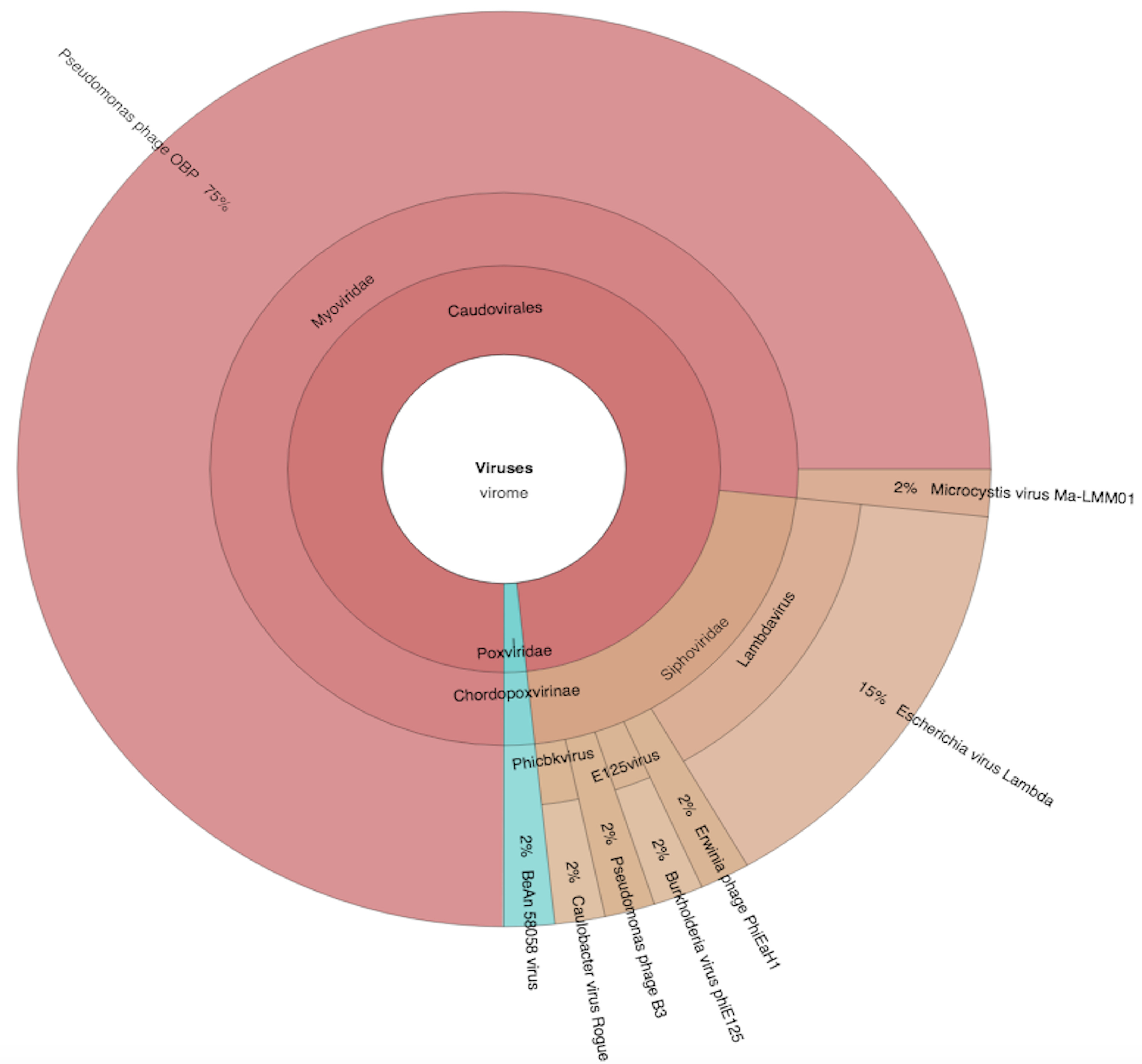

**Figure S5.** Circular plot depicting virus profile identified using Nanopore sequencing technology

**Table S1**. Sample collection sites on the Ganges river

| **S. No.** | **Sampling Location** | **Longitude** | **Latitude** |
| --- | --- | --- | --- |
| 1 | Bijnor-Upstream | N 29°22’28” | E 78°02’29” |
| 2 | Bijnor-Downstream | N 29°17’06” | E 7806’06” |
| 3 | Narora-Upstream | N 28°41’54” | E 78°17’32” |
| 4 | Narora-Downstream | N 28°09’21” | E 78°24’37” |
| 5 | Kannauj-Upstream | N 29°08’98” | E 79°53’19” |
| 6 | Kannauj-Downstream | N 27°64’ | E 79°59’24” |
| 7 | Kanpur-Upstream | N 27°08’98” | E 79°53’19” |
| 8 | Kanpur-Downstream | N 26°24’55” | E 80°26’66” |
| 9 | Allahabad-Sangam | N 25°25’45” | E 81°52’97” |
| 10 | Allahabad-Upstream | N 26°24’55” | E 80°26’66” |
| 11 | Allahabad-Downstream | N 26°24’42” | E 80°26’26” |
| 12 | Mirzapur-Upstream | N 25°10’29” | E 82°29’46” |
| 13 | Mirzapur-Downstream | N 25°09’47” | E 82°33’32” |
| 14 | Varanasi-Upstream | N 25°09’36” | E 82°00’29” |
| 15 | Varanasi-Downstream | N 25°16’26” | E 83°00’97” |

**Table S2.** List of all the identified viruses/phages from the Ganges river sediments using the Illumina sequencing along with relative abundance percentage

| No. | Accession Number | Virus/Phage Name | Number of assisgned reads | Relative abundance (%) | Viral family/ Category |
| --- | --- | --- | --- | --- | --- |
| 1 | NC_008562.1 | Microcystis phage Ma-LMM01 | 498 | 11.93 | Myoviridae |
| 2 | NC_003315.1 | Haemophilus phage HP2 | 403 | 9.66 | Myoviridae |
| 3 | NC_008168.1 | Choristoneura occidentalis granulovirus | 386 | 9.25 | Baculoviridae |
| 4 | NC_029002.1 | Microcystis phage MaMV-DC | 284 | 6.8 | Myoviridae |
| 5 | NC_023719.1 | Bacillus phage G | 155 | 3.71 | Myoviridae |
| 6 | NC_019406.1 | Caulobacter phage CcrColossus | 143 | 3.43 | Siphoviridae |
| 7 | NC_016655.1 | Rhodococcus phage REQ1 | 140 | 3.35 | Siphoviridae |
| 8 | NC_031129.1 | Salmonella phage SJ46 | 130 | 3.11 | Myoviridae |
| 9 | NC_013644.1 | Enterococcus phage phiFL4A | 130 | 3.11 | Siphoviridae |
| 10 | NC_004629.1 | Pseudomonas phage phiKZ | 120 | 2.87 | Myoviridae |
| 11 | NC_032111.1 | BeAn 58058 virus | 108 | 2.59 | Poxviridae |
| 12 | NC_001416.1 | Enterobacteria phage lambda | 99 | 2.37 | Siphoviridae |
| 13 | NC_025453.1 | Enterococcus phage EFC-1 | 95 | 2.28 | Siphoviridae |
| 14 | NC_019929.1 | Erwinia phage phiEaH2 | 65 | 1.56 | Siphoviridae |
| 15 | NC_021536.1 | Synechococcus phage S-IOM18 | 61 | 1.46 | Myoviridae |
| 16 | NC_020486.1 | Synechococcus phage S-RIM8 A.HR1 | 60 | 1.44 | Myoviridae |
| 17 | NC_031253.1 | Mycobacterium phage Bipper | 56 | 1.34 | Siphoviridae |
| 18 | NC_031108.1 | Propionibacterium phage PFR2 | 55 | 1.32 | Siphoviridae |
| 19 | NC_030940.1 | Pseudomonas phage phi3 | 52 | 1.25 | Myoviridae |
| 20 | NC_010811.2 | Ralstonia phage RSL1 | 50 | 1.2 | Myoviridae |
| 21 | NC_006820.1 | Synechococcus phage S-PM2 | 46 | 1.1 | Myoviridae |
| 22 | NC_015279.1 | Synechococcus phage S-SM2 | 45 | 1.08 | Myoviridae |
| 23 | NC_031927.1 | Synechococcus phage S-CAM7 | 33 | 0.79 | Myoviridae |
| 24 | NC_011273.1 | Mycobacterium phage Myrna | 29 | 0.69 | Myoviridae |
| 25 | NC_028768.1 | Achromobacter phage JWX | 27 | 0.65 | Siphoviridae |
| 26 | NC_015286.1 | Synechococcus phage Syn19 | 26 | 0.62 | Myoviridae |
| 27 | NC_027348.2 | Delftia phage RG-2014 | 25 | 0.6 | Podoviridae |
| 28 | NC_030925.1 | Bacillus phage Shbh1 | 20 | 0.48 | Myoviridae |
| 29 | NC_016650.1 | Rhodococcus phage RGL3 | 18 | 0.43 | Siphoviridae |
| 30 | NC_031935.1 | Synechococcus phage S-WAM2 | 15 | 0.36 | Myoviridae |
| 31 | NC_022518.1 | Human endogenous retrovirus K113 | 15 | 0.36 | Retroviridae |
| 32 | NC_031242.1 | Cyanophage S-RIM50 | 15 | 0.36 | Myoviridae |
| 33 | NC_013085.1 | Synechococcus phage S-RSM4 | 14 | 0.34 | Myoviridae |
| 34 | NC_029058.1 | Lactobacillus phage LfeInf | 14 | 0.34 | Myoviridae |
| 35 | NC_001405.1 | Human adenovirus C | 14 | 0.34 | Adenoviridae |
| 36 | NC_031944.1 | Synechococcus phage S-WAM1 | 13 | 0.31 | Myoviridae |
| 37 | NC_020859.1 | Synechococcus phage S-RIM2 R1 1999 | 13 | 0.31 | Myoviridae |
| 38 | NC_019507.1 | Campylobacter phage CP21 | 13 | 0.31 | Myoviridae |
| 39 | NC_007809.1 | Pseudomonas phage M6 | 12 | 0.29 | Siphoviridae |
| 40 | NC_030230.1 | Tokyovirus A1 nearly | 11 | 0.26 | Marseilleviridae |
| 41 | NC_026607.2 | Salmonella phage STP4-a | 11 | 0.26 | Myoviridae |
| 42 | NC_022096.1 | Pseudomonas phage PaBG | 11 | 0.26 | Myoviridae |
| 43 | NC_019411.1 | Caulobacter phage CcrSwift | 11 | 0.26 | Siphoviridae |
| 44 | NC_023006.1 | Pseudomonas phage PPpW-3 | 10 | 0.24 | Myoviridae |
| 45 | NC_028860.1 | Mycobacterium phage Smeadley | 10 | 0.24 | Siphoviridae |
| 46 | NC_026014.1 | Enterobacteria phage P88 | 10 | 0.24 | Myoviridae |
| 47 | NC_005856.1 | Enterobacteria phage P1 | 10 | 0.24 | Myoviridae |
| 48 | NC_016765.1 | Pseudomonas phage vB PaeS PMG1 | 9 | 0.22 | Siphoviridae |
| 49 | NC_010116.1 | Pseudomonas phage YuA | 9 | 0.22 | Siphoviridae |
| 50 | NC_019408.1 | Caulobacter phage CcrRogue | 9 | 0.22 | Siphoviridae |
| 51 | NC_031922.1 | Synechococcus phage S-CAM9 | 8 | 0.19 | Myoviridae |
| 52 | NC_031080.1 | Mycobacterium phage Tonenili | 8 | 0.19 | Myoviridae |
| 53 | NC_013651.1 | Vibrio phage N4 | 7 | 0.17 | Podoviridae |
| 54 | NC_020851.1 | Synechococcus phage S-SKS1 | 7 | 0.17 | Siphoviridae |
| 55 | NC_000871.1 | Streptococcus phage Sfi19 | 7 | 0.17 | Siphoviridae |
| 56 | NC_016652.1 | Rhodococcus phage REQ2 | 7 | 0.17 | Siphoviridae |
| 57 | NC_018282.1 | Pseudomonas phage MP1412 | 7 | 0.17 | Siphoviridae |
| 58 | NC_021312.1 | Phaeocystis globosa virus | 7 | 0.17 | Phycodnaviridae |
| 59 | NC_009603.1 | Microbacterium phage Min1 | 7 | 0.17 | Siphoviridae |
| 60 | NC_007581.1 | Clostridium phage c-st | 7 | 0.17 | Myoviridae |
| 61 | NC_027334.1 | Aurantimonas phage AmM-1 | 7 | 0.17 | Caudovirales; unclassified Caudovirales |
| 62 | NC_015250.1 | Acinetobacter phage 133 | 7 | 0.17 | Myoviridae |
| 63 | NC_029119.1 | Staphylococcus phage SPbeta-like | 6 | 0.14 | Siphoviridae |
| 64 | NC_022749.1 | Shigella phage SfIV | 6 | 0.14 | Myoviridae |
| 65 | NC_028657.1 | Pseudomonas phage YMC11/02/R656 | 6 | 0.14 | Siphoviridae |
| 66 | NC_021299.1 | Mycobacterium phage PegLeg | 6 | 0.14 | Siphoviridae |
| 67 | NC_030696.1 | Gordonia phage Smoothie | 6 | 0.14 | Siphoviridae |
| 68 | NC_019516.1 | Cyanophage S-TIM5 | 6 | 0.14 | Myoviridae |
| 69 | NC_031235.1 | Cyanophage S-RIM32 | 6 | 0.14 | Myoviridae |
| 70 | NC_019401.1 | Cronobacter phage vB CsaM GAP32 | 6 | 0.14 | Myoviridae |
| 71 | NC_007149.1 | Vibriophage VP4 | 5 | 0.12 | Podoviridae |
| 72 | NC_016766.1 | Synechococcus phage S-CBS4 | 5 | 0.12 | Siphoviridae |
| 73 | NC_028879.1 | Pseudomonas phage PaMx42 | 5 | 0.12 | Siphoviridae |
| 74 | NC_028980.1 | Pseudomonas phage PAE1 | 5 | 0.12 | Siphoviridae |
| 75 | NC_026609.1 | Lactobacillus phage Ldl1 | 5 | 0.12 | Siphoviridae |
| 76 | NC_020204.1 | Klebsiella phage JD001 | 5 | 0.12 | Myoviridae |
| 77 | NC_003287.2 | Enterobacteria phage M13 | 5 | 0.12 | Inoviridae |
| 78 | NC_021072.1 | Cyanophage Syn30 | 5 | 0.12 | unclassified dsDNA phages |
| 79 | NC_015265.1 | Burkholderia phage KS5 | 5 | 0.12 | Myoviridae |
| 80 | NC_001884.1 | Bacillus phage SPBc2 | 5 | 0.12 | Siphoviridae |
| 81 | NC_023688.1 | Aeromonas phage PX29 | 5 | 0.12 | Myoviridae |
| 82 | NC_015159.1 | Vibrio phage ICP3 | 4 | 0.1 | Podoviridae |
| 83 | NC_008296.2 | Synechococcus phage syn9 | 4 | 0.1 | Myoviridae |
| 84 | NC_015282.1 | Synechococcus phage S-SM1 | 4 | 0.1 | Myoviridae |
| 85 | NC_025456.1 | Synechococcus phage S-CBP1 | 4 | 0.1 | Podoviridae |
| 86 | NC_027132.1 | Synechococcus phage ACG-2014i | 4 | 0.1 | Myoviridae |
| 87 | NC_013645.1 | Streptococcus phage Abc2 | 4 | 0.1 | Siphoviridae |
| 88 | NC_002185.1 | Streptococcus phage 7201 | 4 | 0.1 | Siphoviridae |
| 89 | NC_009382.1 | Ralstonia phage phiRSA1 | 4 | 0.1 | Myoviridae |
| 90 | NC_016762.1 | Pseudomonas phage phi297 | 4 | 0.1 | Siphoviridae |
| 91 | NC_006552.1 | Pseudomonas phage F116 | 4 | 0.1 | Podoviridae |
| 92 | NC_026440.1 | Pandoravirus inopinatum | 4 | 0.1 | unclassified dsDNA viruses; Pandoravirus |
| 93 | NC_031029.1 | Gordonia phage Cucurbita | 4 | 0.1 | Siphoviridae |
| 94 | NC_019520.1 | Escherichia phage phiKT | 4 | 0.1 | Podoviridae |
| 95 | NC_028695.2 | Enterobacter phage phiEap-2 | 4 | 0.1 | Siphoviridae |
| 96 | NC_023717.1 | Cronobacter phage CR9 | 4 | 0.1 | Myoviridae |
| 97 | NC_019527.1 | Aeromonas phage vB AsaM-56 | 4 | 0.1 | Myoviridae |
| 98 | NC_009542.2 | Aeromonas phage phiO18P | 4 | 0.1 | Myoviridae |
| 99 | NC_019538.1 | Aeromonas phage CC2 | 4 | 0.1 | Myoviridae |
| 100 | NC_015289.1 | Synechococcus phage S-SSM5 | 3 | 0.07 | Myoviridae |
| 101 | NC_031906.1 | Synechococcus phage S-CAM3 | 3 | 0.07 | Myoviridae |
| 102 | NC_020837.1 | Synechococcus phage S-CAM1 | 3 | 0.07 | Myoviridae |
| 103 | NC_021339.1 | Streptomyces phage Zemlya | 3 | 0.07 | Siphoviridae |
| 104 | NC_023575.1 | Pseudomonas phage vB_PaeP_Tr60_Ab31 | 3 | 0.07 | unclassified dsDNA phages |
| 105 | NC_030909.1 | Pseudomonas phage YMC11/07/P54_PAE_BP | 3 | 0.07 | Siphoviridae |
| 106 | NC_028809.1 | Pseudomonas phage PaMx74 | 3 | 0.07 | Siphoviridae |
| 107 | NC_031058.1 | Pseudomonas phage NP1 | 3 | 0.07 | Siphoviridae |
| 108 | NC_027621.1 | Propionibacterium phage Wizzo | 3 | 0.07 | Siphoviridae |
| 109 | NC_027294.1 | Propionibacterium phage PHL150M00 | 3 | 0.07 | Siphoviridae |
| 110 | NC_028936.1 | Mycobacterium phage Enkosi | 3 | 0.07 | Siphoviridae |
| 111 | NC_028878.1 | Mycobacterium phage Archie | 3 | 0.07 | Siphoviridae |
| 112 | NC_027993.1 | Escherichia phage K1G | 3 | 0.07 | Siphoviridae |
| 113 | NC_031107.1 | Erwinia phage vB_EamM_Asesino | 3 | 0.07 | Myoviridae |
| 114 | NC_020875.1 | Cyanophage S-SSM4 | 3 | 0.07 | Myoviridae |
| 115 | NC_003309.1 | Burkholderia phage phiE125 | 3 | 0.07 | Siphoviridae |
| 116 | NC_005882.1 | Burkholderia cenocepacia phage BcepMu | 3 | 0.07 | Myoviridae |
| 117 | NC_004821.1 | Bacillus prophage phBC6A52 | 3 | 0.07 | Podoviridae |
| 118 | NC_027374.1 | Bacillus phage Moonbeam | 3 | 0.07 | Myoviridae |
| 119 | NC_023848.1 | Anopheles minimus irodovirus | 3 | 0.07 | Iridoviridae |
| 120 | NC_028834.1 | Achromobacter phage 83-24 | 3 | 0.07 | Siphoviridae |
| 121 | NC_004745.1 | Yersinia phage L-413C | 2 | 0.05 | Myoviridae |
| 122 | NC_015157.1 | Vibrio phage ICP1 | 2 | 0.05 | Myoviridae |
| 123 | NC_024711.1 | Uncultured crAssphage | 2 | 0.05 | unclassified bacterial viruses; crAss-like viruses |
| 124 | NC_002794.1 | Tupaiid herpesvirus 1 | 2 | 0.05 | Herpesviridae |
| 125 | NC_015210.1 | Tsukamurella phage TPA2 | 2 | 0.05 | Siphoviridae |
| 126 | NC_015465.1 | Synechococcus phage S-CBS3 | 2 | 0.05 | Siphoviridae |
| 127 | NC_029031.1 | Synechococcus phage S-CBP42 | 2 | 0.05 | Podoviridae |
| 128 | NC_025464.1 | Synechococcus phage S-CBP4 | 2 | 0.05 | Podoviridae |
| 129 | NC_026924.1 | Synechococcus phage ACG-2014g | 2 | 0.05 | Myoviridae |
| 130 | NC_026923.1 | Synechococcus phage ACG-2014d | 2 | 0.05 | Myoviridae |
| 131 | NC_021298.1 | Streptomyces phage Lika | 2 | 0.05 | Siphoviridae |
| 132 | NC_018285.1 | Streptococcus phage YMC-2011 | 2 | 0.05 | Siphoviridae |
| 133 | NC_000872.1 | Streptococcus phage Sfi21 | 2 | 0.05 | Siphoviridae |
| 134 | NC_021868.1 | Streptococcus phage SP-QS1 | 2 | 0.05 | Siphoviridae |
| 135 | NC_002072.2 | Streptococcus phage DT1 | 2 | 0.05 | Siphoviridae |
| 136 | NC_010353.1 | Streptococcus phage 858 | 2 | 0.05 | Siphoviridae |
| 137 | NC_023503.1 | Streptococcus phage 20617 | 2 | 0.05 | Siphoviridae |
| 138 | NC_029000.1 | Stenotrophomonas phage IME13 | 2 | 0.05 | Myoviridae |
| 139 | NC_009503.1 | Spodoptera litura granulovirus | 2 | 0.05 | Baculoviridae |
| 140 | NC_023589.1 | Shigella phage pSb-1 | 2 | 0.05 | Podoviridae |
| 141 | NC_018464.1 | Shamonda virus | 2 | 0.05 | Peribunyaviridae |
| 142 | NC_010495.1 | Salmonella phage E1 | 2 | 0.05 | Siphoviridae |
| 143 | NC_020866.1 | Rhodovulum phage RS1 | 2 | 0.05 | Siphoviridae |
| 144 | NC_025115.1 | Ralstonia phage RSY1 | 2 | 0.05 | Myoviridae |
| 145 | NC_028999.1 | Pseudomonas phage PhiPA3 | 2 | 0.05 | Myoviridae |
| 146 | NC_011373.1 | Pseudomonas phage PAJU2 | 2 | 0.05 | Siphoviridae |
| 147 | NC_031091.1 | Pseudomonas phage MD8 | 2 | 0.05 | Siphoviridae |
| 148 | NC_027367.1 | Propionibacterium phage PHL132N00 | 2 | 0.05 | Siphoviridae |
| 149 | NC_027401.1 | Propionibacterium phage PHL095N00 | 2 | 0.05 | Siphoviridae |
| 150 | NC_009551.1 | Phormidium phage Pf-WMP3 | 2 | 0.05 | Podoviridae |
| 151 | NC_032001.1 | Only Syngen Nebraska virus 5 | 2 | 0.05 | Phycodnaviridae |
| 152 | NC_023705.1 | Mycobacterium phage Liefie | 2 | 0.05 | Siphoviridae |
| 153 | NC_009993.2 | Mycobacterium phage Giles | 2 | 0.05 | Siphoviridae |
| 154 | NC_031020.1 | Morganella phage vB_MmoM_MP1 | 2 | 0.05 | Myoviridae |
| 155 | NC_002667.1 | Lactococcus prophage bIL286 | 2 | 0.05 | Siphoviridae |
| 156 | NC_005822.1 | Lactococcus phage phiLC3 | 2 | 0.05 | Siphoviridae |
| 157 | NC_019489.1 | Lactobacillus phage Sha1 | 2 | 0.05 | Siphoviridae |
| 158 | NC_001716.2 | Human herpesvirus 7 | 2 | 0.05 | Herpesviridae |
| 159 | NC_010342.1 | Halomonas phage phiHAP-1 | 2 | 0.05 | Myoviridae |
| 160 | NC_031097.1 | Gordonia phage Zirinka | 2 | 0.05 | Siphoviridae |
| 161 | NC_031061.1 | Gordonia phage Nymphadora | 2 | 0.05 | Siphoviridae |
| 162 | NC_030902.1 | Gordonia phage GMA1 | 2 | 0.05 | Siphoviridae |
| 163 | NC_027125.1 | Flavobacterium phage FCL-2 | 2 | 0.05 | Myoviridae |
| 164 | NC_021867.1 | Flavobacterium phage 6H | 2 | 0.05 | unclassified dsDNA phages |
| 165 | NC_031926.1 | Flavobacterium phage 2A | 2 | 0.05 | Podoviridae |
| 166 | NC_019445.1 | Escherichia phage TL-2011b | 2 | 0.05 | Podoviridae |
| 167 | NC_027994.1 | Escherichia phage K1H | 2 | 0.05 | Siphoviridae |
| 168 | NC_031010.1 | Erwinia phage vB_EamM_Kwan | 2 | 0.05 | Myoviridae |
| 169 | NC_001650.2 | Equid herpesvirus 2 | 2 | 0.05 | Herpesviridae |
| 170 | NC_029016.1 | Enterococcus phage vB_EfaS_IME198 | 2 | 0.05 | Siphoviridae |
| 171 | NC_009904.1 | Enterococcus phage phiEF24C | 2 | 0.05 | Myoviridae |
| 172 | NC_024212.1 | Enterococcus phage VD13 | 2 | 0.05 | Siphoviridae |
| 173 | NC_019720.1 | Enterobacterial phage mEp213 | 2 | 0.05 | Siphoviridae |
| 174 | NC_019517.1 | Enterobacteria phage vB_EcoM-FV3 | 2 | 0.05 | Myoviridae |
| 175 | NC_003356.1 | Enterobacteria phage phiP27 | 2 | 0.05 | Myoviridae |
| 176 | NC_019716.1 | Enterobacteria phage mEp460 | 2 | 0.05 | Siphoviridae |
| 177 | NC_029028.1 | Enterobacteria phage JenP1 | 2 | 0.05 | Siphoviridae |
| 178 | NC_029021.1 | Enterobacteria phage JenK1 | 2 | 0.05 | Siphoviridae |
| 179 | NC_010463.1 | Enterobacteria phage Fels-2 | 2 | 0.05 | Myoviridae |
| 180 | NC_019423.1 | Enterobacter phage IME11 | 2 | 0.05 | Podoviridae |
| 181 | NC_021531.1 | Cronobacter phage CR5 | 2 | 0.05 | Myoviridae |
| 182 | NC_021325.1 | Clostridium phage vB_CpeS-CP51 | 2 | 0.05 | Siphoviridae |
| 183 | NC_028857.1 | Citrobacter phage Merlin | 2 | 0.05 | Myoviridae |
| 184 | NC_021792.1 | Cellulophaga phage phi46:3 | 2 | 0.05 | Podoviridae |
| 185 | NC_021790.1 | Cellulophaga phage phi18:1 | 2 | 0.05 | Siphoviridae |
| 186 | NC_015273.1 | Burkholderia phage KS14 | 2 | 0.05 | Myoviridae |
| 187 | NC_005887.1 | Burkholderia phage BcepC6B | 2 | 0.05 | Podoviridae |
| 188 | NC_005262.3 | Burkholderia cepacia phage Bcep22 | 2 | 0.05 | Podoviridae |
| 189 | NC_005357.1 | Bordetella phage BPP-1 | 2 | 0.05 | Podoviridae |
| 190 | NC_004167.1 | Bacillus phage phi105 | 2 | 0.05 | Siphoviridae |
| 191 | NC_019502.1 | Bacillus phage phIS3501 | 2 | 0.05 | Siphoviridae |
| 192 | NC_028890.1 | Bacillus phage TsarBomba | 2 | 0.05 | Myoviridae |
| 193 | NC_029069.1 | Bacillus phage BM5 | 2 | 0.05 | Myoviridae |
| 194 | NC_001623.1 | Autographa californica nucleopolyhedrovirus | 2 | 0.05 | Baculoviridae |
| 195 | NC_019543.1 | Aeromonas phage Aes508 | 2 | 0.05 | Myoviridae |
| 196 | NC_005885.1 | Actinoplanes phage phiAsp2 | 2 | 0.05 | Siphoviridae |
| 197 | NC_028908.2 | Achromobacter phage phiAxp-3 | 2 | 0.05 | Podoviridae |
| 198 | NC_028104.1 | Yellowstone lake mimivirus | 1 | 0.02 | Mimiviridae |
| 199 | NC_019529.1 | Vibrio phage pVp-1 | 1 | 0.02 | Siphoviridae |
| 200 | NC_014594.1 | Tomato leaf curl Bangladesh betasatellite | 1 | 0.02 | Tolecusatellitidae |
| 201 | NC_015287.1 | Synechococcus phage S-SSM7 | 1 | 0.02 | Myoviridae |
| 202 | NC_023584.1 | Synechococcus phage S-MbCM100 | 1 | 0.02 | Myoviridae |
| 203 | NC_015463.1 | Synechococcus phage S-CBS2 | 1 | 0.02 | Siphoviridae |
| 204 | NC_031903.1 | Synechococcus phage S-CAM22 | 1 | 0.02 | Myoviridae |
| 205 | NC_026927.1 | Synechococcus phage ACG-2014f | 1 | 0.02 | Myoviridae |
| 206 | NC_018836.1 | Streptomyces phage phiHau3 | 1 | 0.02 | Siphoviridae |
| 207 | NC_028827.1 | Streptomyces phage Lannister | 1 | 0.02 | Siphoviridae |
| 208 | NC_029098.1 | Streptomyces phage Jay2Jay | 1 | 0.02 | Siphoviridae |
| 209 | NC_028976.1 | Streptomyces phage Izzy | 1 | 0.02 | Siphoviridae |
| 210 | NC_030946.1 | Streptococcus phage phiARI0923 | 1 | 0.02 | Siphoviridae |
| 211 | NC_018277.1 | Staphylococcus phage SpaA1 | 1 | 0.02 | Siphoviridae |
| 212 | NC_027991.1 | Staphylococcus phage SA1 | 1 | 0.02 | Myoviridae |
| 213 | NC_029025.1 | Staphylococcus phage IME-SA4 | 1 | 0.02 | Siphoviridae |
| 214 | NC_008722.1 | Staphylococcus phage CNPH82 | 1 | 0.02 | Siphoviridae |
| 215 | NC_025829.1 | Shigella phage pSs-1 | 1 | 0.02 | Myoviridae |
| 216 | NC_031011.1 | Shigella phage SHFML-26 | 1 | 0.02 | Myoviridae |
| 217 | NC_027329.1 | Salmonella phage vB_SPuM_SP116 | 1 | 0.02 | Myoviridae |
| 218 | NC_031924.1 | Salmonella phage IME207 | 1 | 0.02 | Siphoviridae |
| 219 | NC_021782.1 | Salmonella phage FSL SP-076 | 1 | 0.02 | Podoviridae |
| 220 | NC_028250.1 | Rosellinia necatrix partitivirus 6 | 1 | 0.02 | Partitiviridae |
| 221 | NC_016654.1 | Rhodococcus phage REQ3 | 1 | 0.02 | Siphoviridae |
| 222 | NC_034248.1 | Rhizobium phage RHEph10 | 1 | 0.02 | Caudovirales; unclassified Caudovirales |
| 223 | NC_026599.1 | Pseudomonas phage vB_PaeP_C2-10_Ab22 | 1 | 0.02 | Podoviridae |
| 224 | NC_003278.1 | Pseudomonas phage phiCTX | 1 | 0.02 | Myoviridae |
| 225 | NC_001331.1 | Pseudomonas phage Pf1 | 1 | 0.02 | Inoviridae |
| 226 | NC_028931.1 | Pseudomonas phage PaMx28 | 1 | 0.02 | Siphoviridae |
| 227 | NC_007808.1 | Pseudomonas phage PA11 | 1 | 0.02 | unclassified dsDNA phages |
| 228 | NC_029017.1 | Pseudomonas phage KPP21 | 1 | 0.02 | Podoviridae |
| 229 | NC_019923.1 | Pseudomonas phage AF | 1 | 0.02 | Podoviridae |
| 230 | NC_031005.1 | Propionibacterium phage QueenBey | 1 | 0.02 | Siphoviridae |
| 231 | NC_027359.1 | Propionibacterium phage PHL082M00 | 1 | 0.02 | Siphoviridae |
| 232 | NC_022336.1 | Propionibacterium phage PHL010M04 | 1 | 0.02 | Siphoviridae |
| 233 | NC_027336.1 | Propionibacterium phage PHL009M11 | 1 | 0.02 | Siphoviridae |
| 234 | NC_009541.1 | Propionibacterium phage PA6 | 1 | 0.02 | Siphoviridae |
| 235 | NC_018849.1 | Propionibacterium phage P105 | 1 | 0.02 | Siphoviridae |
| 236 | NC_006884.2 | Prochlorococcus phage P-SSM4 | 1 | 0.02 | Myoviridae |
| 237 | NC_006883.2 | Prochlorococcus phage P-SSM2 | 1 | 0.02 | Myoviridae |
| 238 | NC_020878.1 | Prochlorococcus phage P-GSP1 | 1 | 0.02 | Podoviridae |
| 239 | NC_028998.1 | Phormidium phage MIS-PhV1B | 1 | 0.02 | unclassified dsDNA phages |
| 240 | NC_018837.1 | Pectobacterium phage My1 | 1 | 0.02 | Siphoviridae |
| 241 | NC_021858.1 | Pandoravirus dulcis | 1 | 0.02 | unclassified dsDNA viruses; Pandoravirus |
| 242 | NC_022067.1 | Mycobacterium phage Wanda | 1 | 0.02 | Siphoviridae |
| 243 | NC_029077.1 | Mycobacterium phage VohminGhazi | 1 | 0.02 | Siphoviridae |
| 244 | NC_026591.1 | Mycobacterium phage Trike | 1 | 0.02 | Siphoviridae |
| 245 | NC_028960.2 | Mycobacterium phage Theia | 1 | 0.02 | Siphoviridae |
| 246 | NC_023725.1 | Mycobacterium phage Nappy | 1 | 0.02 | Myoviridae |
| 247 | NC_028759.1 | Mycobacterium phage Mufasa | 1 | 0.02 | Siphoviridae |
| 248 | NC_004685.1 | Mycobacterium phage Corndog | 1 | 0.02 | Siphoviridae |
| 249 | NC_004680.1 | Mycobacterium phage Che8 | 1 | 0.02 | Siphoviridae |
| 250 | NC_028742.1 | Mycobacterium phage Baee | 1 | 0.02 | Siphoviridae |
| 251 | NC_022068.1 | Mycobacteriophage Daenerys | 1 | 0.02 | Siphoviridae |
| 252 | NC_009811.2 | Listeria phage A511 | 1 | 0.02 | Myoviridae |
| 253 | NC_000896.1 | Lactobacillus prophage phiadh | 1 | 0.02 | Siphoviridae |
| 254 | NC_005857.1 | Klebsiella phage phiKO2 | 1 | 0.02 | Siphoviridae |
| 255 | NC_023613.1 | Invertebrate iridovirus 25 | 1 | 0.02 | Iridoviridae |
| 256 | NC_023611.1 | Invertebrate iridescent virus 30 | 1 | 0.02 | Iridoviridae |
| 257 | NC_031052.1 | Gordonia phage Twister6 | 1 | 0.02 | Siphoviridae |
| 258 | NC_030917.1 | Gordonia phage OneUp | 1 | 0.02 | Siphoviridae |
| 259 | NC_030914.1 | Gordonia phage Kvothe | 1 | 0.02 | Siphoviridae |
| 260 | NC_031074.1 | Gordonia phage Bantam | 1 | 0.02 | Siphoviridae |
| 261 | NC_030936.1 | Gordonia phage Bachita | 1 | 0.02 | Siphoviridae |
| 262 | NC_025447.1 | Escherichia phage 121Q | 1 | 0.02 | Myoviridae |
| 263 | NC_028671.1 | Enterococcus phage vB_EfaS_IME197 | 1 | 0.02 | Siphoviridae |
| 264 | NC_021190.1 | Enterobacteria phage phi80 | 1 | 0.02 | Siphoviridae |
| 265 | NC_019708.1 | Enterobacteria phage mEp235 | 1 | 0.02 | Siphoviridae |
| 266 | NC_005066.1 | Enterobacteria phage RB49 | 1 | 0.02 | Myoviridae |
| 267 | NC_025419.1 | Enterobacteria phage RB3 | 1 | 0.02 | Myoviridae |
| 268 | NC_012740.1 | Enterobacteria phage JSE | 1 | 0.02 | Myoviridae |
| 269 | NC_001479.1 | Encephalomyocarditis virus | 1 | 0.02 | Picornaviridae |
| 270 | NC_021342.2 | Edwardsiella phage PEi21 | 1 | 0.02 | Myoviridae |
| 271 | NC_028702.1 | Delftia phage IME-DE1 | 1 | 0.02 | Podoviridae |
| 272 | NC_028663.1 | Cyanophage P-TIM40 | 1 | 0.02 | Myoviridae |
| 273 | NC_021071.1 | Cyanophage P-RSM1 | 1 | 0.02 | Myoviridae |
| 274 | NC_006659.1 | Cotesia congregata virus | 1 | 0.02 | Polydnaviridae |
| 275 | NC_011318.1 | Clostridium phage 39-O | 1 | 0.02 | Siphoviridae |
| 276 | NC_023153.1 | Citrus endogenous pararetrovirus | 1 | 0.02 | Retroviridae |
| 277 | NC_027988.2 | Citrobacter phage CVT22 | 1 | 0.02 | Podoviridae |
| 278 | NC_021788.1 | Cellulophaga phage phi4:1 | 1 | 0.02 | Podoviridae |
| 279 | NC_024709.1 | Ball python nidovirus | 1 | 0.02 | Coronaviridae |
| 280 | NC_030904.1 | Bacillus phage vB_BhaS-171 | 1 | 0.02 | Siphoviridae |
| 281 | NC_031039.1 | Bacillus phage AR9 | 1 | 0.02 | Myoviridae |
| 282 | NC_022972.2 | Arthrobacter phage vB_ArS-ArV2 | 1 | 0.02 | Siphoviridae |
| 283 | NC_031098.1 | Acinetobacter phage vB_AbaS_TRS1 | 1 | 0.02 | Siphoviridae |
| 284 | NC_019541.1 | Acinetobacter phage YMC/09/02/B1251_ABA_BP | 1 | 0.02 | Siphoviridae |
| 285 | NC_031117.1 | Acinetobacter phage LZ35 | 1 | 0.02 | Myoviridae |

**Table S3**. Relative distribution of reads belongs to identified phages from different bacterial group/genus

| Phage Group | Phylum | Family | Genus | Hits |
| --- | --- | --- | --- | --- |
| Microcystis | Cyanobacteria | Microcystaceae | Microcystis | 782 |
| Haemophilus | Proteobacteria | Pasteurellaceae | Haemophilus | 403 |
| Synechococcus | Cyanobacteria | Synechococcaceae | Synechococcus | 386 |
| Pseudomonas | Proteobacteria | Pseudomonadaceae | Pseudomonas | 279 |
| Enterococcus |  | Enterococcaceae | Enterococcus | 232 |
| Bacillus | Firmicutes | Bacillaceae | Bacillus | 196 |
| Rhodococcus | Actinobacteria | Nocardiaceae | Rhodococcus | 166 |
| Caulobacter | Proteobacteria |  | Caulobacter | 163 |
| Salmonella | Eubacteria | Enterobacteriaceae | Salmonella | 146 |
| Enterobacteria | Proteobacteria | Enterobacteriaceae |  | 143 |
| Mycobacterium | Actinobacteria | Mycobacteriaceae | Mycobacterium | 128 |
| Propionibacterium | Actinobacteria | Propionibacteriaceae | Propionibacterium | 71 |
| Erwinia | Proteobacteria | Enterobacteriaceae | Erwinia | 70 |
| Ralstonia | Proteobacteria | Ralstoniaceae/Burkholderiaceae | Ralstonia | 56 |
| Cyanophage | Cyanobacteria |  |  | 37 |
| Achromobacter | Proteobacteria | Alcaligenaceae | Achromobacter | 32 |
| Streptococcus | Firmicutes | Streptococcaceae | Streptococcus | 28 |
| Delftia | Proteobacteria | Comamonadaceae | Delftia | 26 |
| Lactobacillus | Firmicutes | Lactobacillaceae | Lactobacillus | 22 |
| Gordonia | Actinobacteria | Gordoniaceae | Gordonia | 21 |
| Aeromonas | Proteobacteria | Aeromonadaceae | Aeromonas | 19 |
| Burkholderia | Proteobacteria | Burkholderiaceae | Burkholderia | 17 |
| Vibrio |  | Vibrionaceae | Vibrio | 14 |
| Campylobacter |  | Campylobacteraceae | Campylobacter | 13 |
| Escherichia | Proteobacteria | Enterobacteriaceae | Escherichia | 12 |
| Cronobacter | Proteobacteria | Enterobacteriaceae | Cronobacter | 12 |
| Staphylococcus | Firmicutes | Staphylococcaceae | Staphylococcus | 10 |
| Shigella | Proteobacteria | Enterobacteriaceae | Shigella | 10 |
| Clostridium | Firmicutes | Clostridiaceae | Clostridium | 10 |
| Acinetobacter | Proteobacteria | Moraxellaceae | Acinetobacter | 10 |
| Streptomyces | Actinobacteria | Streptomycetaceae | Streptomyces | 9 |
| Microbacterium | Actinobacteria | Microbacteriaceae | Microbacterium | 7 |
| Aurantimonas | Proteobacteria | Aurantimonadaceae | Aurantimonas | 7 |
| Klebsiella | Proteobacteria | Enterobacteriaceae | Klebsiella | 6 |
| Flavobacterium | Bacteroidetes | Flavobacteriaceae | Flavobacterium | 6 |
| Enterobacter | Proteobacteria | Enterobacteriaceae | Enterobacter | 6 |
| Vibriophage |  |  | Vibrio | 5 |
| Cellulophaga | Bacteroidetes | Flavobacteriaceae | Cellulophaga | 5 |
| Lactococcus | Firmicutes | Streptococcaceae | Lactococcus | 4 |
| Prochlorococcus | Cyanobacteria | Prochloraceae/Synechococcaceae | Prochlorococcus | 3 |
| Phormidium | Cyanobacteria | Oscillatoriophycideae/  Oscillatoriaceae | Phormidium | 3 |
| Citrobacter | Proteobacteria | Enterobacteriaceae | Citrobacter | 3 |
| Yersinia |  | Enterobacteriaceae | Yersinia | 2 |
| crAssphage |  |  |  | 2 |
| Tsukamurella | Actinobacteria | Tsukamurellaceae | Tsukamurella | 2 |
| Stenotrophomonas | Proteobacteria | Xanthomonadaceae | Stenotrophomonas | 2 |
| Rhodovulum | Proteobacteria | Rhodobacteraceae | Rhodovulum | 2 |
| Morganella |  |  | Morganella | 2 |
| Halomonas | Proteobacteria | Halomonadaceae | Halomonas | 2 |
| Bordetella | Proteobacteria | Alcaligenaceae | Bordetella | 2 |
| Actinoplanes | Actinobacteria | Micromonosporaceae | Actinoplanes | 2 |
| Rhizobium | Proteobacteria | ‎Rhizobiaceae | ‎Rhizobium | 1 |
| Pectobacterium | Proteobacteria | Enterobacteriaceae | Pectobacterium | 1 |
| Mycobacteriophage |  |  | Mycobacterium | 1 |
| Listeria |  | Listeriaceae | Listeria | 1 |
| Edwardsiella | Proteobacteria | Enterobacteriaceae | Edwardsiella | 1 |
| Arthrobacter | Actinobacteria | Micrococcaceae | Arthrobacter | 1 |

**Table S4**. Diversity and abundance of microbial community in the Ganges sediment

| Clade name | Relative abundance (%) | Estimated number of reads |
| --- | --- | --- |
| Bacteria | 94.415 | 942294 |
| Archaea | 5.585 | 31286 |
| Bacteria-Proteobacteria | 69.291 | 708514 |
| Bacteria-Actinobacteria | 15.079 | 158831 |
| Archaea-Euryarchaeota | 5.096 | 32472 |
| Bacteria-Cyanobacteria | 3.928 | 66805 |
| Bacteria-Bacteroidetes | 2.449 | 33814 |
| Bacteria-Firmicutes | 2.411 | 20802 |
| Bacteria-Nitrospirae | 1.205 | 11090 |
| Archaea-Thaumarchaeota | 0.489 | 3028 |
| Bacteria-Spirochaetes | 0.052 | 574 |
| Bacteria-Proteobacteria-Alphaproteobacteria | 32.690 | 393680 |
| Bacteria-Proteobacteria-Betaproteobacteria | 25.314 | 252605 |
| Bacteria-Actinobacteria-Actinobacteria | 15.079 | 158831 |
| Bacteria-Proteobacteria-Gammaproteobacteria | 10.741 | 129102 |
| Archaea-Euryarchaeota-Methanomicrobia | 4.531 | 39636 |
| Bacteria-Cyanobacteria-Cyanobacteria_noname | 3.928 | 73506 |
| Bacteria-Bacteroidetes-Sphingobacteriia | 2.201 | 44248 |
| Bacteria-Firmicutes-Bacilli | 2.111 | 18645 |
| Bacteria-Nitrospirae-Nitrospira | 1.205 | 11090 |
| Archaea-Euryarchaeota-Methanobacteria | 0.565 | 3056 |
| Bacteria-Proteobacteria-Deltaproteobacteria | 0.546 | 7697 |
| Archaea-Thaumarchaeota-Thaumarchaeota_noname | 0.489 | 3028 |
| Bacteria-Firmicutes-Clostridia | 0.301 | 2818 |
| Bacteria-Bacteroidetes-Flavobacteriia | 0.191 | 1770 |
| Bacteria-Spirochaetes-Spirochaetia | 0.052 | 574 |
| Bacteria-Bacteroidetes-Bacteroidia | 0.047 | 576 |
| Bacteria-Bacteroidetes-Cytophagia | 0.010 | 178 |
| Bacteria-Proteobacteria-Alphaproteobacteria-Caulobacterales | 28.100 | 341952 |
| Bacteria-Proteobacteria-Betaproteobacteria-Burkholderiales | 13.873 | 181075 |
| Bacteria-Actinobacteria-Actinobacteria-Actinomycetales | 13.682 | 216866 |
| Bacteria-Proteobacteria-Gammaproteobacteria-Pseudomonadales | 7.482 | 100680 |
| Bacteria-Proteobacteria-Betaproteobacteria-Rhodocyclales | 6.932 | 92424 |
| Bacteria-Cyanobacteria-Cyanobacteria_noname-Chroococcales | 3.849 | 56582 |
| Bacteria-Proteobacteria-Betaproteobacteria-Hydrogenophilales | 3.398 | 35244 |
| Bacteria-Proteobacteria-Alphaproteobacteria-Rhizobiales | 3.365 | 46426 |
| Archaea-Euryarchaeota-Methanomicrobia-Methanosarcinales | 3.310 | 28274 |
| Bacteria-Proteobacteria-Gammaproteobacteria-Alteromonadales | 2.233 | 32786 |
| Bacteria-Bacteroidetes-Sphingobacteriia-Sphingobacteriales | 2.201 | 44248 |
| Bacteria-Firmicutes-Bacilli-Bacillales | 2.111 | 23454 |
| Bacteria-Actinobacteria-Actinobacteria-Solirubrobacterales | 1.397 | 25380 |
| Archaea-Euryarchaeota-Methanomicrobia-Methanomicrobiales | 1.220 | 10557 |
| Bacteria-Nitrospirae-Nitrospira-Nitrospirales | 1.205 | 11090 |
| Bacteria-Proteobacteria-Betaproteobacteria-Methylophilales | 1.111 | 7595 |
| Bacteria-Proteobacteria-Alphaproteobacteria-Sphingomonadales | 0.758 | 8927 |
| Archaea-Euryarchaeota-Methanobacteria-Methanobacteriales | 0.565 | 3056 |
| Bacteria-Proteobacteria-Deltaproteobacteria-Myxococcales | 0.467 | 15967 |
| Bacteria-Proteobacteria-Gammaproteobacteria-Xanthomonadales | 0.457 | 6290 |
| Archaea-Thaumarchaeota-Thaumarchaeota_noname-Nitrososphaerales | 0.431 | 3984 |
| Bacteria-Proteobacteria-Alphaproteobacteria-Rhodobacterales | 0.332 | 5158 |
| Bacteria-Firmicutes-Clostridia-Clostridiales | 0.301 | 3065 |
| Bacteria-Proteobacteria-Gammaproteobacteria-Methylococcales | 0.268 | 3642 |
| Bacteria-Bacteroidetes-Flavobacteriia-Flavobacteriales | 0.191 | 1546 |
| Bacteria-Proteobacteria-Gammaproteobacteria-Oceanospirillales | 0.143 | 2109 |
| Bacteria-Proteobacteria-Alphaproteobacteria-Rhodospirillales | 0.124 | 1784 |
| Bacteria-Cyanobacteria-Cyanobacteria_noname-Oscillatoriales | 0.079 | 1726 |
| Bacteria-Proteobacteria-Gammaproteobacteria-Enterobacteriales | 0.060 | 793 |
| Archaea-Thaumarchaeota-Thaumarchaeota_noname-Nitrosopumilales | 0.058 | 351 |
| Bacteria-Proteobacteria-Deltaproteobacteria-Syntrophobacterales | 0.053 | 777 |
| Bacteria-Spirochaetes-Spirochaetia-Spirochaetales | 0.052 | 574 |
| Bacteria-Proteobacteria-Gammaproteobacteria-Aeromonadales | 0.052 | 654 |
| Bacteria-Bacteroidetes-Bacteroidia-Bacteroidales | 0.047 | 576 |
| Bacteria-Proteobacteria-Gammaproteobacteria-Chromatiales | 0.046 | 541 |
| Bacteria-Proteobacteria-Deltaproteobacteria-Desulfuromonadales | 0.026 | 345 |
| Bacteria-Proteobacteria-Alphaproteobacteria-Rickettsiales | 0.010 | 56 |
| Bacteria-Bacteroidetes-Cytophagia-Cytophagales | 0.010 | 178 |
| Bacteria-Proteobacteria-Alphaproteobacteria-Caulobacterales-Caulobacteraceae | 28.100 | 341952 |
| Bacteria-Proteobacteria-Betaproteobacteria-Burkholderiales-Oxalobacteraceae | 7.908 | 112273 |
| Bacteria-Actinobacteria-Actinobacteria-Actinomycetales-Micrococcaceae | 7.071 | 69311 |
| Bacteria-Proteobacteria-Betaproteobacteria-Rhodocyclales-Rhodocyclaceae | 6.932 | 92424 |
| Bacteria-Proteobacteria-Gammaproteobacteria-Pseudomonadales-Pseudomonadaceae | 6.757 | 119743 |
| Bacteria-Cyanobacteria-Cyanobacteria_noname-Chroococcales-Chroococcales_noname | 3.849 | 56582 |
| Bacteria-Proteobacteria-Betaproteobacteria-Burkholderiales-Burkholderiales_noname | 3.437 | 45218 |
| Bacteria-Proteobacteria-Betaproteobacteria-Hydrogenophilales-Hydrogenophilaceae | 3.398 | 35244 |
| Bacteria-Proteobacteria-Alphaproteobacteria-Rhizobiales-Hyphomicrobiaceae | 2.869 | 37910 |
| Archaea-Euryarchaeota-Methanomicrobia-Methanosarcinales-Methanosarcinaceae | 2.746 | 24676 |
| Bacteria-Proteobacteria-Betaproteobacteria-Burkholderiales-Comamonadaceae | 2.528 | 39786 |
| Bacteria-Bacteroidetes-Sphingobacteriia-Sphingobacteriales-Sphingobacteriaceae | 2.201 | 41523 |
| Bacteria-Proteobacteria-Gammaproteobacteria-Alteromonadales-Alteromonadaceae | 2.158 | 31763 |
| Bacteria-Actinobacteria-Actinobacteria-Actinomycetales-Nocardiaceae | 1.755 | 36769 |
| Bacteria-Firmicutes-Bacilli-Bacillales-Planococcaceae | 1.640 | 20820 |
| Bacteria-Actinobacteria-Actinobacteria-Solirubrobacterales-Solirubrobacterales_unclassified | 1.397 | 25380 |
| Bacteria-Nitrospirae-Nitrospira-Nitrospirales-Nitrospiraceae | 1.205 | 11090 |
| Archaea-Euryarchaeota-Methanomicrobia-Methanomicrobiales-Methanoregulaceae | 1.138 | 9462 |
| Bacteria-Proteobacteria-Betaproteobacteria-Methylophilales-Methylophilaceae | 1.111 | 10463 |
| Bacteria-Actinobacteria-Actinobacteria-Actinomycetales-Dermatophilaceae | 0.810 | 10513 |
| Bacteria-Actinobacteria-Actinobacteria-Actinomycetales-Microbacteriaceae | 0.750 | 8235 |
| Bacteria-Proteobacteria-Gammaproteobacteria-Pseudomonadales-Moraxellaceae | 0.726 | 6668 |
| Bacteria-Actinobacteria-Actinobacteria-Actinomycetales-Nocardioidaceae | 0.711 | 14437 |
| Bacteria-Proteobacteria-Alphaproteobacteria-Sphingomonadales-Sphingomonadaceae | 0.682 | 8190 |
| Bacteria-Actinobacteria-Actinobacteria-Actinomycetales-Micromonosporaceae | 0.621 | 14548 |
| Archaea-Euryarchaeota-Methanobacteria-Methanobacteriales-Methanobacteriaceae | 0.565 | 3821 |
| Archaea-Euryarchaeota-Methanomicrobia-Methanosarcinales-Methanosaetaceae | 0.565 | 4572 |
| Archaea-Thaumarchaeota-Thaumarchaeota_noname-Nitrososphaerales-Nitrososphaeraceae | 0.431 | 3984 |
| Bacteria-Proteobacteria-Deltaproteobacteria-Myxococcales-Myxococcaceae | 0.415 | 11158 |
| Bacteria-Actinobacteria-Actinobacteria-Actinomycetales-Streptosporangiaceae | 0.397 | 12521 |
| Bacteria-Proteobacteria-Alphaproteobacteria-Rhodobacterales-Rhodobacteraceae | 0.332 | 5460 |
| Bacteria-Proteobacteria-Gammaproteobacteria-Xanthomonadales-Xanthomonadaceae | 0.331 | 4131 |
| Bacteria-Firmicutes-Bacilli-Bacillales-Bacillales_noname | 0.304 | 2650 |
| Bacteria-Proteobacteria-Alphaproteobacteria-Rhizobiales-Bradyrhizobiaceae | 0.301 | 5039 |
| Bacteria-Proteobacteria-Gammaproteobacteria-Methylococcales-Methylococcaceae | 0.268 | 3642 |
| Bacteria-Firmicutes-Clostridia-Clostridiales-Peptostreptococcaceae | 0.247 | 1939 |
| Bacteria-Actinobacteria-Actinobacteria-Actinomycetales-Geodermatophilaceae | 0.236 | 4051 |
| Bacteria-Actinobacteria-Actinobacteria-Actinomycetales-Thermomonosporaceae | 0.190 | 4115 |
| Bacteria-Actinobacteria-Actinobacteria-Actinomycetales-Streptomycetaceae | 0.185 | 5227 |
| Bacteria-Actinobacteria-Actinobacteria-Actinomycetales-Cellulomonadaceae | 0.181 | 2259 |
| Bacteria-Actinobacteria-Actinobacteria-Actinomycetales-Dietziaceae | 0.174 | 2050 |
| Bacteria-Firmicutes-Bacilli-Bacillales-Bacillaceae | 0.166 | 1984 |
| Bacteria-Actinobacteria-Actinobacteria-Actinomycetales-Pseudonocardiaceae | 0.153 | 3571 |
| Bacteria-Bacteroidetes-Flavobacteriia-Flavobacteriales-Flavobacteriales_noname | 0.151 | 623 |
| Bacteria-Proteobacteria-Gammaproteobacteria-Oceanospirillales-Halomonadaceae | 0.143 | 1398 |
| Bacteria-Proteobacteria-Gammaproteobacteria-Xanthomonadales-Sinobacteraceae | 0.126 | 1898 |
| Bacteria-Proteobacteria-Alphaproteobacteria-Rhodospirillales-Rhodospirillaceae | 0.124 | 1580 |
| Bacteria-Actinobacteria-Actinobacteria-Actinomycetales-Intrasporangiaceae | 0.119 | 1635 |
| Bacteria-Actinobacteria-Actinobacteria-Actinomycetales-Propionibacteriaceae | 0.113 | 1251 |
| Bacteria-Actinobacteria-Actinobacteria-Actinomycetales-Brevibacteriaceae | 0.111 | 1180 |
| Bacteria-Proteobacteria-Alphaproteobacteria-Rhizobiales-Aurantimonadaceae | 0.108 | 2037 |
| Archaea-Euryarchaeota-Methanomicrobia-Methanomicrobiales-Methanospirillaceae | 0.083 | 958 |
| Bacteria-Actinobacteria-Actinobacteria-Actinomycetales-Promicromonosporaceae | 0.080 | 1158 |
| Bacteria-Cyanobacteria-Cyanobacteria_noname-Oscillatoriales-Oscillatoriales_noname | 0.079 | 1726 |
| Bacteria-Proteobacteria-Alphaproteobacteria-Sphingomonadales-Erythrobacteraceae | 0.076 | 876 |
| Bacteria-Proteobacteria-Gammaproteobacteria-Alteromonadales-Idiomarinaceae | 0.075 | 828 |
| Bacteria-Proteobacteria-Alphaproteobacteria-Rhizobiales-Phyllobacteriaceae | 0.066 | 1107 |
| Bacteria-Proteobacteria-Gammaproteobacteria-Enterobacteriales-Enterobacteriaceae | 0.060 | 793 |
| Archaea-Thaumarchaeota-Thaumarchaeota_noname-Nitrosopumilales-Nitrosopumilaceae | 0.058 | 351 |
| Bacteria-Firmicutes-Clostridia-Clostridiales-Clostridiaceae | 0.053 | 583 |
| Bacteria-Proteobacteria-Deltaproteobacteria-Syntrophobacterales-Syntrophobacteraceae | 0.053 | 866 |
| Bacteria-Spirochaetes-Spirochaetia-Spirochaetales-Leptospiraceae | 0.052 | 748 |
| Bacteria-Proteobacteria-Gammaproteobacteria-Aeromonadales-Aeromonadaceae | 0.052 | 638 |
| Bacteria-Proteobacteria-Deltaproteobacteria-Myxococcales-Polyangiaceae | 0.052 | 2345 |
| Bacteria-Bacteroidetes-Bacteroidia-Bacteroidales-Porphyromonadaceae | 0.047 | 609 |
| Bacteria-Proteobacteria-Gammaproteobacteria-Chromatiales-Chromatiaceae | 0.046 | 711 |
| Bacteria-Bacteroidetes-Flavobacteriia-Flavobacteriales-Flavobacteriaceae | 0.041 | 490 |
| Bacteria-Proteobacteria-Deltaproteobacteria-Desulfuromonadales-Geobacteraceae | 0.026 | 375 |
| Bacteria-Actinobacteria-Actinobacteria-Actinomycetales-Mycobacteriaceae | 0.023 | 385 |
| Bacteria-Proteobacteria-Alphaproteobacteria-Rhizobiales-Rhizobiaceae | 0.021 | 340 |
| Bacteria-Proteobacteria-Alphaproteobacteria-Rickettsiales-Holosporaceae | 0.010 | 47 |
| Bacteria-Bacteroidetes-Cytophagia-Cytophagales-Cytophagaceae | 0.010 | 193 |
| Bacteria-Proteobacteria-Alphaproteobacteria-Caulobacterales-Caulobacteraceae-Brevundimonas | 24.690 | 270199 |
| Bacteria-Actinobacteria-Actinobacteria-Actinomycetales-Micrococcaceae-Arthrobacter | 7.015 | 100468 |
| Bacteria-Proteobacteria-Gammaproteobacteria-Pseudomonadales-Pseudomonadaceae-Pseudomonas | 6.708 | 136630 |
| Bacteria-Proteobacteria-Betaproteobacteria-Burkholderiales-Oxalobacteraceae-Massilia | 5.922 | 122932 |
| Bacteria-Proteobacteria-Betaproteobacteria-Rhodocyclales-Rhodocyclaceae-Thauera | 4.860 | 80035 |
| Bacteria-Proteobacteria-Betaproteobacteria-Burkholderiales-Burkholderiales_noname-Thiomonas | 3.434 | 42562 |
| Bacteria-Proteobacteria-Alphaproteobacteria-Rhizobiales-Hyphomicrobiaceae-Hyphomicrobiaceae_unclassified | 2.830 | 37397 |
| Archaea-Euryarchaeota-Methanomicrobia-Methanosarcinales-Methanosarcinaceae-Methanosarcina | 2.666 | 41592 |
| Bacteria-Proteobacteria-Alphaproteobacteria-Caulobacterales-Caulobacteraceae-Caulobacter | 2.527 | 28854 |
| Bacteria-Cyanobacteria-Cyanobacteria_noname-Chroococcales-Chroococcales_noname-Microcystis | 2.488 | 39555 |
| Bacteria-Proteobacteria-Betaproteobacteria-Hydrogenophilales-Hydrogenophilaceae-Hydrogenophilaceae_unclassified | 2.126 | 22048 |
| Bacteria-Bacteroidetes-Sphingobacteriia-Sphingobacteriales-Sphingobacteriaceae-Pedobacter | 2.099 | 34170 |
| Bacteria-Proteobacteria-Betaproteobacteria-Burkholderiales-Oxalobacteraceae-Janthinobacterium | 1.970 | 36094 |
| Bacteria-Actinobacteria-Actinobacteria-Actinomycetales-Nocardiaceae-Rhodococcus | 1.755 | 38394 |
| Bacteria-Proteobacteria-Gammaproteobacteria-Alteromonadales-Alteromonadaceae-Marinobacter | 1.732 | 24921 |
| Bacteria-Firmicutes-Bacilli-Bacillales-Planococcaceae-Paenisporosarcina | 1.640 | 20434 |
| Bacteria-Proteobacteria-Betaproteobacteria-Burkholderiales-Comamonadaceae-Polaromonas | 1.633 | 25948 |
| Bacteria-Proteobacteria-Betaproteobacteria-Rhodocyclales-Rhodocyclaceae-Azoarcus | 1.555 | 25995 |
| Bacteria-Proteobacteria-Betaproteobacteria-Hydrogenophilales-Hydrogenophilaceae-Thiobacillus | 1.252 | 13185 |
| Bacteria-Nitrospirae-Nitrospira-Nitrospirales-Nitrospiraceae-Nitrospira | 1.205 | 16957 |
| Bacteria-Cyanobacteria-Cyanobacteria_noname-Chroococcales-Chroococcales_noname-Cyanobium | 1.128 | 11359 |
| Bacteria-Proteobacteria-Betaproteobacteria-Methylophilales-Methylophilaceae-Methylotenera | 1.111 | 9843 |
| Archaea-Euryarchaeota-Methanomicrobia-Methanomicrobiales-Methanoregulaceae-Methanoregulaceae_unclassified | 1.081 | 8991 |
| Bacteria-Proteobacteria-Alphaproteobacteria-Caulobacterales-Caulobacteraceae-Asticcacaulis | 0.848 | 13001 |
| Bacteria-Actinobacteria-Actinobacteria-Actinomycetales-Dermatophilaceae-Dermatophilaceae_unclassified | 0.810 | 10513 |
| Bacteria-Actinobacteria-Actinobacteria-Actinomycetales-Nocardioidaceae-Nocardioides | 0.711 | 10534 |
| Archaea-Euryarchaeota-Methanomicrobia-Methanosarcinales-Methanosaetaceae-Methanosaeta | 0.565 | 4572 |
| Bacteria-Proteobacteria-Betaproteobacteria-Rhodocyclales-Rhodocyclaceae-Methyloversatilis | 0.505 | 6877 |
| Bacteria-Actinobacteria-Actinobacteria-Actinomycetales-Microbacteriaceae-Agromyces | 0.471 | 6172 |
| Archaea-Thaumarchaeota-Thaumarchaeota_noname-Nitrososphaerales-Nitrososphaeraceae-Nitrososphaera | 0.431 | 3984 |
| Bacteria-Proteobacteria-Gammaproteobacteria-Alteromonadales-Alteromonadaceae-Alishewanella | 0.426 | 5056 |
| Bacteria-Actinobacteria-Actinobacteria-Actinomycetales-Streptosporangiaceae-Streptosporangiaceae_unclassified | 0.397 | 12521 |
| Bacteria-Proteobacteria-Deltaproteobacteria-Myxococcales-Myxococcaceae-Anaeromyxobacter | 0.389 | 6487 |
| Bacteria-Proteobacteria-Gammaproteobacteria-Pseudomonadales-Moraxellaceae-Acinetobacter | 0.368 | 4469 |
| Bacteria-Proteobacteria-Gammaproteobacteria-Pseudomonadales-Moraxellaceae-Perlucidibaca | 0.357 | 2823 |
| Bacteria-Actinobacteria-Actinobacteria-Actinomycetales-Micromonosporaceae-Actinoplanes | 0.348 | 10813 |
| Bacteria-Proteobacteria-Alphaproteobacteria-Sphingomonadales-Sphingomonadaceae-Sandarakinorhabdus | 0.336 | 3146 |
| Archaea-Euryarchaeota-Methanobacteria-Methanobacteriales-Methanobacteriaceae-Methanobrevibacter | 0.334 | 2478 |
| Bacteria-Proteobacteria-Alphaproteobacteria-Rhodobacterales-Rhodobacteraceae-Paracoccus | 0.332 | 4011 |
| Bacteria-Firmicutes-Bacilli-Bacillales-Bacillales_noname-Exiguobacterium | 0.304 | 3005 |
| Bacteria-Proteobacteria-Alphaproteobacteria-Rhizobiales-Bradyrhizobiaceae-Rhodopseudomonas | 0.301 | 5360 |
| Bacteria-Proteobacteria-Betaproteobacteria-Burkholderiales-Comamonadaceae-Variovorax | 0.288 | 6110 |
| Bacteria-Proteobacteria-Gammaproteobacteria-Xanthomonadales-Xanthomonadaceae-Pseudoxanthomonas | 0.284 | 3151 |
| Bacteria-Proteobacteria-Gammaproteobacteria-Methylococcales-Methylococcaceae-Methylomonas | 0.256 | 4298 |
| Bacteria-Firmicutes-Clostridia-Clostridiales-Peptostreptococcaceae-Peptostreptococcaceae_noname | 0.247 | 2629 |
| Bacteria-Actinobacteria-Actinobacteria-Actinomycetales-Geodermatophilaceae-Geodermatophilaceae_unclassified | 0.236 | 4051 |
| Bacteria-Proteobacteria-Betaproteobacteria-Burkholderiales-Comamonadaceae-Alicycliphilus | 0.236 | 3006 |
| Bacteria-Cyanobacteria-Cyanobacteria_noname-Chroococcales-Chroococcales_noname-Cyanobacterium | 0.233 | 2760 |
| Archaea-Euryarchaeota-Methanobacteria-Methanobacteriales-Methanobacteriaceae-Methanobacterium | 0.231 | 1926 |
| Bacteria-Actinobacteria-Actinobacteria-Actinomycetales-Micromonosporaceae-Salinispora | 0.225 | 4026 |
| Bacteria-Proteobacteria-Betaproteobacteria-Burkholderiales-Comamonadaceae-Limnohabitans | 0.219 | 2295 |
| Bacteria-Proteobacteria-Alphaproteobacteria-Sphingomonadales-Sphingomonadaceae-Sphingobium | 0.210 | 3167 |
| Bacteria-Actinobacteria-Actinobacteria-Actinomycetales-Thermomonosporaceae-Thermomonosporaceae_unclassified | 0.190 | 4115 |
| Bacteria-Actinobacteria-Actinobacteria-Actinomycetales-Streptomycetaceae-Streptomyces | 0.185 | 5148 |
| Bacteria-Actinobacteria-Actinobacteria-Actinomycetales-Cellulomonadaceae-Cellulomonas | 0.181 | 2259 |
| Bacteria-Actinobacteria-Actinobacteria-Actinomycetales-Dietziaceae-Dietzia | 0.174 | 2050 |
| Bacteria-Firmicutes-Bacilli-Bacillales-Bacillaceae-Bacillus | 0.166 | 2532 |
| Bacteria-Actinobacteria-Actinobacteria-Actinomycetales-Pseudonocardiaceae-Saccharomonospora | 0.153 | 2336 |
| Bacteria-Bacteroidetes-Flavobacteriia-Flavobacteriales-Flavobacteriales_noname-Flavobacteriales_noname | 0.151 | 1129 |
| Bacteria-Actinobacteria-Actinobacteria-Actinomycetales-Microbacteriaceae-Leifsonia | 0.143 | 1597 |
| Bacteria-Proteobacteria-Gammaproteobacteria-Oceanospirillales-Halomonadaceae-Halomonas | 0.143 | 1968 |
| Bacteria-Actinobacteria-Actinobacteria-Actinomycetales-Microbacteriaceae-Leucobacter | 0.136 | 1458 |
| Bacteria-Proteobacteria-Gammaproteobacteria-Xanthomonadales-Sinobacteraceae-Sinobacteraceae_unclassified | 0.126 | 1898 |
| Bacteria-Actinobacteria-Actinobacteria-Actinomycetales-Propionibacteriaceae-Propionibacterium | 0.113 | 962 |
| Bacteria-Proteobacteria-Alphaproteobacteria-Sphingomonadales-Sphingomonadaceae-Sphingopyxis | 0.112 | 1221 |
| Bacteria-Actinobacteria-Actinobacteria-Actinomycetales-Brevibacteriaceae-Brevibacterium | 0.111 | 1180 |
| Bacteria-Proteobacteria-Alphaproteobacteria-Rhodospirillales-Rhodospirillaceae-Rhodospirillum | 0.109 | 1489 |
| Bacteria-Proteobacteria-Alphaproteobacteria-Rhizobiales-Aurantimonadaceae-Aurantimonadaceae_unclassified | 0.108 | 2037 |
| Bacteria-Bacteroidetes-Sphingobacteriia-Sphingobacteriales-Sphingobacteriaceae-Sphingobacterium | 0.102 | 1878 |
| Archaea-Euryarchaeota-Methanomicrobia-Methanomicrobiales-Methanospirillaceae-Methanospirillum | 0.083 | 958 |
| Bacteria-Actinobacteria-Actinobacteria-Actinomycetales-Promicromonosporaceae-Promicromonosporaceae_unclassified | 0.080 | 1158 |
| Archaea-Euryarchaeota-Methanomicrobia-Methanosarcinales-Methanosarcinaceae-Methanomethylovorans | 0.079 | 628 |
| Bacteria-Proteobacteria-Alphaproteobacteria-Sphingomonadales-Erythrobacteraceae-Erythrobacteraceae_unclassified | 0.076 | 876 |
| Bacteria-Proteobacteria-Gammaproteobacteria-Alteromonadales-Idiomarinaceae-Idiomarina | 0.075 | 828 |
| Bacteria-Proteobacteria-Betaproteobacteria-Burkholderiales-Comamonadaceae-Acidovorax | 0.073 | 1231 |
| Bacteria-Proteobacteria-Alphaproteobacteria-Rhizobiales-Phyllobacteriaceae-Mesorhizobium | 0.066 | 1474 |
| Bacteria-Actinobacteria-Actinobacteria-Actinomycetales-Intrasporangiaceae-Ornithinimicrobium | 0.065 | 814 |
| Bacteria-Proteobacteria-Gammaproteobacteria-Enterobacteriales-Enterobacteriaceae-Klebsiella | 0.060 | 1105 |
| Bacteria-Actinobacteria-Actinobacteria-Actinomycetales-Micrococcaceae-Kocuria | 0.056 | 590 |
| Bacteria-Actinobacteria-Actinobacteria-Actinomycetales-Intrasporangiaceae-Serinicoccus | 0.055 | 608 |
| Bacteria-Firmicutes-Clostridia-Clostridiales-Clostridiaceae-Youngiibacter | 0.053 | 683 |
| Bacteria-Proteobacteria-Deltaproteobacteria-Syntrophobacterales-Syntrophobacteraceae-Syntrophobacter | 0.053 | 866 |
| Bacteria-Spirochaetes-Spirochaetia-Spirochaetales-Leptospiraceae-Leptonema | 0.052 | 772 |
| Bacteria-Proteobacteria-Gammaproteobacteria-Aeromonadales-Aeromonadaceae-Aeromonas | 0.052 | 749 |
| Bacteria-Proteobacteria-Deltaproteobacteria-Myxococcales-Polyangiaceae-Sorangium | 0.052 | 2345 |
| Archaea-Thaumarchaeota-Thaumarchaeota_noname-Nitrosopumilales-Nitrosopumilaceae-Candidatus_Nitrosoarchaeum | 0.049 | 277 |
| Bacteria-Proteobacteria-Gammaproteobacteria-Pseudomonadales-Pseudomonadaceae-Cellvibrio | 0.049 | 751 |
| Bacteria-Actinobacteria-Actinobacteria-Actinomycetales-Micromonosporaceae-Micromonospora | 0.048 | 1075 |
| Bacteria-Proteobacteria-Gammaproteobacteria-Xanthomonadales-Xanthomonadaceae-Stenotrophomonas | 0.047 | 740 |
| Bacteria-Bacteroidetes-Bacteroidia-Bacteroidales-Porphyromonadaceae-Parabacteroides | 0.047 | 843 |
| Bacteria-Proteobacteria-Gammaproteobacteria-Chromatiales-Chromatiaceae-Rheinheimera | 0.046 | 624 |
| Bacteria-Proteobacteria-Betaproteobacteria-Burkholderiales-Comamonadaceae-Verminephrobacter | 0.042 | 706 |
| Bacteria-Cyanobacteria-Cyanobacteria_noname-Oscillatoriales-Oscillatoriales_noname-Oscillatoria | 0.041 | 992 |
| Bacteria-Bacteroidetes-Flavobacteriia-Flavobacteriales-Flavobacteriaceae-Myroides | 0.041 | 523 |
| Archaea-Euryarchaeota-Methanomicrobia-Methanomicrobiales-Methanoregulaceae-Methanosphaerula | 0.040 | 381 |
| Bacteria-Proteobacteria-Alphaproteobacteria-Rhizobiales-Hyphomicrobiaceae-Hyphomicrobium | 0.039 | 531 |
| Bacteria-Cyanobacteria-Cyanobacteria_noname-Oscillatoriales-Oscillatoriales_noname-Microcoleus | 0.039 | 895 |
| Bacteria-Proteobacteria-Betaproteobacteria-Burkholderiales-Comamonadaceae-Comamonas | 0.037 | 899 |
| Bacteria-Proteobacteria-Alphaproteobacteria-Caulobacterales-Caulobacteraceae-Phenylobacterium | 0.034 | 441 |
| Bacteria-Proteobacteria-Deltaproteobacteria-Desulfuromonadales-Geobacteraceae-Geobacter | 0.026 | 384 |
| Bacteria-Actinobacteria-Actinobacteria-Actinomycetales-Mycobacteriaceae-Mycobacterium | 0.023 | 416 |
| Bacteria-Proteobacteria-Alphaproteobacteria-Sphingomonadales-Sphingomonadaceae-Citromicrobium | 0.022 | 227 |
| Bacteria-Proteobacteria-Betaproteobacteria-Hydrogenophilales-Hydrogenophilaceae-Sulfuricella | 0.021 | 211 |
| Bacteria-Proteobacteria-Deltaproteobacteria-Myxococcales-Myxococcaceae-Myxococcus | 0.019 | 595 |
| Archaea-Euryarchaeota-Methanomicrobia-Methanomicrobiales-Methanoregulaceae-Methanoregula | 0.017 | 146 |
| Bacteria-Proteobacteria-Alphaproteobacteria-Rhodospirillales-Rhodospirillaceae-Azospirillum | 0.015 | 208 |
| Bacteria-Proteobacteria-Betaproteobacteria-Burkholderiales-Oxalobacteraceae-Herbaspirillum | 0.015 | 254 |
| Bacteria-Proteobacteria-Alphaproteobacteria-Rhizobiales-Rhizobiaceae-Sinorhizobium | 0.014 | 252 |
| Bacteria-Proteobacteria-Betaproteobacteria-Rhodocyclales-Rhodocyclaceae-Azospira | 0.012 | 147 |
| Bacteria-Proteobacteria-Gammaproteobacteria-Methylococcales-Methylococcaceae-Methylobacter | 0.011 | 106 |
| Bacteria-Proteobacteria-Alphaproteobacteria-Rickettsiales-Holosporaceae-Holospora | 0.010 | 47 |
| Archaea-Thaumarchaeota-Thaumarchaeota_noname-Nitrosopumilales-Nitrosopumilaceae-Nitrosopumilaceae_unclassified | 0.008 | 51 |
| Bacteria-Proteobacteria-Deltaproteobacteria-Myxococcales-Myxococcaceae-Corallococcus | 0.007 | 245 |
| Bacteria-Bacteroidetes-Cytophagia-Cytophagales-Cytophagaceae-Hymenobacter | 0.007 | 113 |
| Bacteria-Proteobacteria-Alphaproteobacteria-Rhizobiales-Rhizobiaceae-Kaistia | 0.004 | 58 |
| Bacteria-Proteobacteria-Alphaproteobacteria-Rhizobiales-Rhizobiaceae-Agrobacterium | 0.003 | 58 |
| Bacteria-Bacteroidetes-Cytophagia-Cytophagales-Cytophagaceae-Pontibacter | 0.003 | 44 |
| Bacteria-Proteobacteria-Betaproteobacteria-Burkholderiales-Burkholderiales_noname-Burkholderiales_noname | 0.003 | 38 |
| Bacteria-Proteobacteria-Alphaproteobacteria-Sphingomonadales-Sphingomonadaceae-Blastomonas | 0.003 | 35 |
| Bacteria-Proteobacteria-Alphaproteobacteria-Caulobacterales-Caulobacteraceae-Brevundimonas-Brevundimonas_unclassified | 22.890 | 250493 |
| Bacteria-Actinobacteria-Actinobacteria-Actinomycetales-Micrococcaceae-Arthrobacter-Arthrobacter_phenanthrenivorans | 7.015 | 97229 |
| Bacteria-Proteobacteria-Gammaproteobacteria-Pseudomonadales-Pseudomonadaceae-Pseudomonas-Pseudomonas_unclassified | 5.969 | 121571 |
| Bacteria-Proteobacteria-Betaproteobacteria-Burkholderiales-Oxalobacteraceae-Massilia-Massilia_unclassified | 5.922 | 122932 |
| Bacteria-Proteobacteria-Betaproteobacteria-Rhodocyclales-Rhodocyclaceae-Thauera-Thauera_unclassified | 4.510 | 74270 |
| Bacteria-Proteobacteria-Betaproteobacteria-Burkholderiales-Burkholderiales_noname-Thiomonas-Thiomonas_unclassified | 3.434 | 42562 |
| Bacteria-Proteobacteria-Alphaproteobacteria-Caulobacterales-Caulobacteraceae-Caulobacter-Caulobacter_unclassified | 2.375 | 27113 |
| Bacteria-Bacteroidetes-Sphingobacteriia-Sphingobacteriales-Sphingobacteriaceae-Pedobacter-Pedobacter_unclassified | 2.099 | 34170 |
| Bacteria-Proteobacteria-Betaproteobacteria-Burkholderiales-Oxalobacteraceae-Janthinobacterium-Janthinobacterium_unclassified | 1.970 | 36094 |
| Archaea-Euryarchaeota-Methanomicrobia-Methanosarcinales-Methanosarcinaceae-Methanosarcina-Methanosarcina_unclassified | 1.806 | 28175 |
| Bacteria-Proteobacteria-Alphaproteobacteria-Caulobacterales-Caulobacteraceae-Brevundimonas-Brevundimonas_diminuta | 1.801 | 19692 |
| Bacteria-Cyanobacteria-Cyanobacteria_noname-Chroococcales-Chroococcales_noname-Microcystis-Microcystis_aeruginosa | 1.768 | 29145 |
| Bacteria-Proteobacteria-Gammaproteobacteria-Alteromonadales-Alteromonadaceae-Marinobacter-Marinobacter_unclassified | 1.642 | 23636 |
| Bacteria-Firmicutes-Bacilli-Bacillales-Planococcaceae-Paenisporosarcina-Paenisporosarcina_unclassified | 1.640 | 20434 |
| Bacteria-Proteobacteria-Betaproteobacteria-Burkholderiales-Comamonadaceae-Polaromonas-Polaromonas_unclassified | 1.633 | 25948 |
| Bacteria-Proteobacteria-Betaproteobacteria-Rhodocyclales-Rhodocyclaceae-Azoarcus-Azoarcus_unclassified | 1.555 | 25995 |
| Bacteria-Proteobacteria-Betaproteobacteria-Hydrogenophilales-Hydrogenophilaceae-Thiobacillus-Thiobacillus_denitrificans | 1.252 | 13305 |
| Bacteria-Nitrospirae-Nitrospira-Nitrospirales-Nitrospiraceae-Nitrospira-Candidatus_Nitrospira_defluvii | 1.205 | 16957 |
| Bacteria-Cyanobacteria-Cyanobacteria_noname-Chroococcales-Chroococcales_noname-Cyanobium-Cyanobium_unclassified | 1.128 | 11359 |
| Bacteria-Proteobacteria-Betaproteobacteria-Methylophilales-Methylophilaceae-Methylotenera-Methylotenera_unclassified | 1.111 | 9843 |
| Bacteria-Proteobacteria-Alphaproteobacteria-Caulobacterales-Caulobacteraceae-Asticcacaulis-Asticcacaulis_unclassified | 0.848 | 13001 |
| Archaea-Euryarchaeota-Methanomicrobia-Methanosarcinales-Methanosarcinaceae-Methanosarcina-Methanosarcina_mazei | 0.766 | 9398 |
| Bacteria-Cyanobacteria-Cyanobacteria_noname-Chroococcales-Chroococcales_noname-Microcystis-Microcystis_unclassified | 0.720 | 11452 |
| Bacteria-Actinobacteria-Actinobacteria-Actinomycetales-Nocardioidaceae-Nocardioides-Nocardioides_unclassified | 0.711 | 10534 |
| Bacteria-Actinobacteria-Actinobacteria-Actinomycetales-Nocardiaceae-Rhodococcus-Rhodococcus_qingshengii | 0.637 | 13719 |
| Bacteria-Actinobacteria-Actinobacteria-Actinomycetales-Nocardiaceae-Rhodococcus-Rhodococcus_erythropolis | 0.620 | 13488 |
| Bacteria-Proteobacteria-Gammaproteobacteria-Pseudomonadales-Pseudomonadaceae-Pseudomonas-Pseudomonas_stutzeri | 0.563 | 8518 |
| Bacteria-Proteobacteria-Betaproteobacteria-Rhodocyclales-Rhodocyclaceae-Methyloversatilis-Methyloversatilis_unclassified | 0.505 | 6877 |
| Bacteria-Actinobacteria-Actinobacteria-Actinomycetales-Microbacteriaceae-Agromyces-Agromyces_unclassified | 0.471 | 6172 |
| Archaea-Thaumarchaeota-Thaumarchaeota_noname-Nitrososphaerales-Nitrososphaeraceae-Nitrososphaera-Candidatus_Nitrososphaera_gargensis | 0.431 | 3984 |
| Bacteria-Proteobacteria-Gammaproteobacteria-Alteromonadales-Alteromonadaceae-Alishewanella-Alishewanella_agri | 0.426 | 4854 |
| Bacteria-Proteobacteria-Deltaproteobacteria-Myxococcales-Myxococcaceae-Anaeromyxobacter-Anaeromyxobacter_unclassified | 0.389 | 6487 |
| Archaea-Euryarchaeota-Methanomicrobia-Methanosarcinales-Methanosaetaceae-Methanosaeta-Methanosaeta_unclassified | 0.380 | 3074 |
| Bacteria-Proteobacteria-Gammaproteobacteria-Pseudomonadales-Moraxellaceae-Perlucidibaca-Perlucidibaca_piscinae | 0.357 | 2823 |
| Bacteria-Actinobacteria-Actinobacteria-Actinomycetales-Nocardiaceae-Rhodococcus-Rhodococcus_ruber | 0.349 | 6553 |
| Bacteria-Actinobacteria-Actinobacteria-Actinomycetales-Micromonosporaceae-Actinoplanes-Actinoplanes_unclassified | 0.348 | 10813 |
| Bacteria-Proteobacteria-Alphaproteobacteria-Sphingomonadales-Sphingomonadaceae-Sandarakinorhabdus-Sandarakinorhabdus_unclassified | 0.336 | 3146 |
| Archaea-Euryarchaeota-Methanobacteria-Methanobacteriales-Methanobacteriaceae-Methanobrevibacter-Methanobrevibacter_unclassified | 0.334 | 2478 |
| Bacteria-Proteobacteria-Alphaproteobacteria-Rhodobacterales-Rhodobacteraceae-Paracoccus-Paracoccus_unclassified | 0.332 | 4011 |
| Bacteria-Proteobacteria-Alphaproteobacteria-Rhizobiales-Bradyrhizobiaceae-Rhodopseudomonas-Rhodopseudomonas_palustris | 0.301 | 5306 |
| Bacteria-Proteobacteria-Betaproteobacteria-Burkholderiales-Comamonadaceae-Variovorax-Variovorax_unclassified | 0.288 | 6110 |
| Bacteria-Proteobacteria-Gammaproteobacteria-Xanthomonadales-Xanthomonadaceae-Pseudoxanthomonas-Pseudoxanthomonas_unclassified | 0.284 | 3151 |
| Bacteria-Proteobacteria-Gammaproteobacteria-Pseudomonadales-Moraxellaceae-Acinetobacter-Acinetobacter_unclassified | 0.273 | 3315 |
| Bacteria-Proteobacteria-Gammaproteobacteria-Methylococcales-Methylococcaceae-Methylomonas-Methylomonas_unclassified | 0.256 | 4298 |
| Bacteria-Proteobacteria-Betaproteobacteria-Burkholderiales-Comamonadaceae-Alicycliphilus-Alicycliphilus_unclassified | 0.236 | 3006 |
| Bacteria-Cyanobacteria-Cyanobacteria_noname-Chroococcales-Chroococcales_noname-Cyanobacterium-Cyanobacterium_stanieri | 0.233 | 2399 |
| Archaea-Euryarchaeota-Methanobacteria-Methanobacteriales-Methanobacteriaceae-Methanobacterium-Methanobacterium_unclassified | 0.231 | 1926 |
| Bacteria-Actinobacteria-Actinobacteria-Actinomycetales-Micromonosporaceae-Salinispora-Salinispora_unclassified | 0.225 | 4026 |
| Bacteria-Firmicutes-Bacilli-Bacillales-Bacillales_noname-Exiguobacterium-Exiguobacterium_unclassified | 0.223 | 2202 |
| Bacteria-Proteobacteria-Betaproteobacteria-Burkholderiales-Comamonadaceae-Limnohabitans-Limnohabitans_unclassified | 0.219 | 2295 |
| Bacteria-Firmicutes-Clostridia-Clostridiales-Peptostreptococcaceae-Peptostreptococcaceae_noname-Peptostreptococcaceae_noname_unclassified | 0.212 | 2256 |
| Bacteria-Proteobacteria-Alphaproteobacteria-Sphingomonadales-Sphingomonadaceae-Sphingobium-Sphingobium_unclassified | 0.210 | 3167 |
| Bacteria-Proteobacteria-Betaproteobacteria-Rhodocyclales-Rhodocyclaceae-Thauera-Thauera_aminoaromatica | 0.192 | 2678 |
| Bacteria-Actinobacteria-Actinobacteria-Actinomycetales-Streptomycetaceae-Streptomyces-Streptomyces_chartreusis | 0.185 | 5496 |
| Archaea-Euryarchaeota-Methanomicrobia-Methanosarcinales-Methanosaetaceae-Methanosaeta-Methanosaeta_concilii | 0.185 | 1815 |
| Bacteria-Actinobacteria-Actinobacteria-Actinomycetales-Cellulomonadaceae-Cellulomonas-Cellulomonas_unclassified | 0.181 | 2259 |
| Bacteria-Actinobacteria-Actinobacteria-Actinomycetales-Dietziaceae-Dietzia-Dietzia_unclassified | 0.169 | 1995 |
| Bacteria-Firmicutes-Bacilli-Bacillales-Bacillaceae-Bacillus-Bacillus_megaterium | 0.166 | 2780 |
| Bacteria-Proteobacteria-Betaproteobacteria-Rhodocyclales-Rhodocyclaceae-Thauera-Thauera_phenylacetica | 0.158 | 2591 |
| Bacteria-Actinobacteria-Actinobacteria-Actinomycetales-Pseudonocardiaceae-Saccharomonospora-Saccharomonospora_unclassified | 0.153 | 2336 |
| Bacteria-Proteobacteria-Alphaproteobacteria-Caulobacterales-Caulobacteraceae-Caulobacter-Caulobacter_vibrioides | 0.153 | 2107 |
| Bacteria-Bacteroidetes-Flavobacteriia-Flavobacteriales-Flavobacteriales_noname-Flavobacteriales_noname-Flavobacteria_bacterium_BAL38 | 0.151 | 1379 |
| Bacteria-Actinobacteria-Actinobacteria-Actinomycetales-Microbacteriaceae-Leifsonia-Leifsonia_unclassified | 0.143 | 1597 |
| Bacteria-Proteobacteria-Gammaproteobacteria-Oceanospirillales-Halomonadaceae-Halomonas-Halomonas_unclassified | 0.143 | 1968 |
| Bacteria-Actinobacteria-Actinobacteria-Actinomycetales-Microbacteriaceae-Leucobacter-Leucobacter_unclassified | 0.136 | 1458 |
| Bacteria-Actinobacteria-Actinobacteria-Actinomycetales-Propionibacteriaceae-Propionibacterium-Propionibacterium_acnes | 0.113 | 925 |
| Bacteria-Proteobacteria-Alphaproteobacteria-Sphingomonadales-Sphingomonadaceae-Sphingopyxis-Sphingopyxis_unclassified | 0.112 | 1221 |
| Bacteria-Actinobacteria-Actinobacteria-Actinomycetales-Brevibacteriaceae-Brevibacterium-Brevibacterium_unclassified | 0.111 | 1180 |
| Bacteria-Proteobacteria-Alphaproteobacteria-Rhodospirillales-Rhodospirillaceae-Rhodospirillum-Rhodospirillum_unclassified | 0.109 | 1489 |
| Bacteria-Proteobacteria-Gammaproteobacteria-Pseudomonadales-Pseudomonadaceae-Pseudomonas-Pseudomonas_mandelii | 0.105 | 3555 |
| Bacteria-Bacteroidetes-Sphingobacteriia-Sphingobacteriales-Sphingobacteriaceae-Sphingobacterium-Sphingobacterium_unclassified | 0.102 | 1878 |
| Bacteria-Proteobacteria-Gammaproteobacteria-Pseudomonadales-Moraxellaceae-Acinetobacter-Acinetobacter_indicus | 0.095 | 992 |
| Bacteria-Proteobacteria-Gammaproteobacteria-Alteromonadales-Alteromonadaceae-Marinobacter-Marinobacter_hydrocarbonoclasticus | 0.089 | 1211 |
| Archaea-Euryarchaeota-Methanomicrobia-Methanomicrobiales-Methanospirillaceae-Methanospirillum-Methanospirillum_hungatei | 0.083 | 958 |
| Bacteria-Firmicutes-Bacilli-Bacillales-Bacillales_noname-Exiguobacterium-Exiguobacterium_pavilionensis | 0.081 | 801 |
| Archaea-Euryarchaeota-Methanomicrobia-Methanosarcinales-Methanosarcinaceae-Methanomethylovorans-Methanomethylovorans_hollandica | 0.079 | 628 |
| Archaea-Euryarchaeota-Methanomicrobia-Methanosarcinales-Methanosarcinaceae-Methanosarcina-Methanosarcina_barkeri | 0.078 | 1227 |
| Bacteria-Proteobacteria-Gammaproteobacteria-Alteromonadales-Idiomarinaceae-Idiomarina-Idiomarina_unclassified | 0.075 | 828 |
| Bacteria-Actinobacteria-Actinobacteria-Actinomycetales-Nocardiaceae-Rhodococcus-Rhodococcus_rhodochrous | 0.075 | 1520 |
| Bacteria-Proteobacteria-Betaproteobacteria-Burkholderiales-Comamonadaceae-Acidovorax-Acidovorax_radicis | 0.073 | 1325 |
| Bacteria-Proteobacteria-Gammaproteobacteria-Pseudomonadales-Pseudomonadaceae-Pseudomonas-Pseudomonas_mendocina | 0.072 | 1223 |
| Bacteria-Proteobacteria-Alphaproteobacteria-Rhizobiales-Phyllobacteriaceae-Mesorhizobium-Mesorhizobium_unclassified | 0.066 | 1474 |
| Bacteria-Actinobacteria-Actinobacteria-Actinomycetales-Intrasporangiaceae-Ornithinimicrobium-Ornithinimicrobium_pekingense | 0.065 | 814 |
| Bacteria-Actinobacteria-Actinobacteria-Actinomycetales-Micrococcaceae-Kocuria-Kocuria_unclassified | 0.056 | 590 |
| Bacteria-Actinobacteria-Actinobacteria-Actinomycetales-Intrasporangiaceae-Serinicoccus-Serinicoccus_unclassified | 0.055 | 608 |
| Bacteria-Firmicutes-Clostridia-Clostridiales-Clostridiaceae-Youngiibacter-Youngiibacter_fragilis | 0.053 | 683 |
| Bacteria-Proteobacteria-Deltaproteobacteria-Syntrophobacterales-Syntrophobacteraceae-Syntrophobacter-Syntrophobacter_fumaroxidans | 0.053 | 866 |
| Bacteria-Spirochaetes-Spirochaetia-Spirochaetales-Leptospiraceae-Leptonema-Leptonema_illini | 0.052 | 772 |
| Bacteria-Proteobacteria-Gammaproteobacteria-Aeromonadales-Aeromonadaceae-Aeromonas-Aeromonas_unclassified | 0.052 | 749 |
| Bacteria-Proteobacteria-Deltaproteobacteria-Myxococcales-Polyangiaceae-Sorangium-Sorangium_cellulosum | 0.052 | 2345 |
| Archaea-Thaumarchaeota-Thaumarchaeota_noname-Nitrosopumilales-Nitrosopumilaceae-Candidatus_Nitrosoarchaeum-Candidatus_Nitrosoarchaeum_unclassified | 0.049 | 277 |
| Bacteria-Proteobacteria-Gammaproteobacteria-Pseudomonadales-Pseudomonadaceae-Cellvibrio-Cellvibrio_unclassified | 0.049 | 751 |
| Bacteria-Actinobacteria-Actinobacteria-Actinomycetales-Micromonosporaceae-Micromonospora-Micromonospora_unclassified | 0.048 | 1075 |
| Bacteria-Actinobacteria-Actinobacteria-Actinomycetales-Nocardiaceae-Rhodococcus-Rhodococcus_pyridinivorans | 0.047 | 809 |
| Bacteria-Proteobacteria-Gammaproteobacteria-Xanthomonadales-Xanthomonadaceae-Stenotrophomonas-Stenotrophomonas_maltophilia | 0.047 | 706 |
| Bacteria-Bacteroidetes-Bacteroidia-Bacteroidales-Porphyromonadaceae-Parabacteroides-Parabacteroides_unclassified | 0.047 | 843 |
| Bacteria-Proteobacteria-Gammaproteobacteria-Chromatiales-Chromatiaceae-Rheinheimera-Rheinheimera_nanhaiensis | 0.046 | 596 |
| Bacteria-Proteobacteria-Gammaproteobacteria-Enterobacteriales-Enterobacteriaceae-Klebsiella-Klebsiella_pneumoniae | 0.045 | 810 |
| Bacteria-Proteobacteria-Betaproteobacteria-Burkholderiales-Comamonadaceae-Verminephrobacter-Verminephrobacter_unclassified | 0.042 | 706 |
| Bacteria-Cyanobacteria-Cyanobacteria_noname-Oscillatoriales-Oscillatoriales_noname-Oscillatoria-Oscillatoria_acuminata | 0.041 | 1021 |
| Bacteria-Bacteroidetes-Flavobacteriia-Flavobacteriales-Flavobacteriaceae-Myroides-Myroides_unclassified | 0.041 | 523 |
| Archaea-Euryarchaeota-Methanomicrobia-Methanomicrobiales-Methanoregulaceae-Methanosphaerula-Methanosphaerula_palustris | 0.040 | 381 |
| Bacteria-Proteobacteria-Alphaproteobacteria-Rhizobiales-Hyphomicrobiaceae-Hyphomicrobium-Hyphomicrobium_unclassified | 0.039 | 531 |
| Bacteria-Cyanobacteria-Cyanobacteria_noname-Oscillatoriales-Oscillatoriales_noname-Microcoleus-Microcoleus_unclassified | 0.039 | 895 |
| Bacteria-Proteobacteria-Betaproteobacteria-Burkholderiales-Comamonadaceae-Comamonas-Comamonas_unclassified | 0.037 | 899 |
| Bacteria-Proteobacteria-Alphaproteobacteria-Caulobacterales-Caulobacteraceae-Phenylobacterium-Phenylobacterium_zucineum | 0.034 | 441 |
| Bacteria-Actinobacteria-Actinobacteria-Actinomycetales-Nocardiaceae-Rhodococcus-Rhodococcus_equi | 0.027 | 456 |
| Bacteria-Proteobacteria-Deltaproteobacteria-Desulfuromonadales-Geobacteraceae-Geobacter-Geobacter_metallireducens | 0.026 | 335 |
| Bacteria-Firmicutes-Clostridia-Clostridiales-Peptostreptococcaceae-Peptostreptococcaceae_noname-Clostridium_sticklandii | 0.025 | 220 |
| Bacteria-Actinobacteria-Actinobacteria-Actinomycetales-Mycobacteriaceae-Mycobacterium-Mycobacterium_vanbaalenii | 0.023 | 484 |
| Bacteria-Proteobacteria-Alphaproteobacteria-Sphingomonadales-Sphingomonadaceae-Citromicrobium-Citromicrobium_unclassified | 0.022 | 227 |
| Bacteria-Proteobacteria-Betaproteobacteria-Hydrogenophilales-Hydrogenophilaceae-Sulfuricella-Sulfuricella_denitrificans | 0.021 | 211 |
| Bacteria-Proteobacteria-Deltaproteobacteria-Myxococcales-Myxococcaceae-Myxococcus-Myxococcus_unclassified | 0.019 | 595 |
| Archaea-Euryarchaeota-Methanomicrobia-Methanomicrobiales-Methanoregulaceae-Methanoregula-Methanoregula_formicica | 0.017 | 153 |
| Archaea-Euryarchaeota-Methanomicrobia-Methanosarcinales-Methanosarcinaceae-Methanosarcina-Methanosarcina_acetivorans | 0.016 | 305 |
| Bacteria-Proteobacteria-Alphaproteobacteria-Rhodospirillales-Rhodospirillaceae-Azospirillum-Azospirillum_unclassified | 0.015 | 208 |
| Bacteria-Proteobacteria-Betaproteobacteria-Burkholderiales-Oxalobacteraceae-Herbaspirillum-Herbaspirillum_unclassified | 0.015 | 254 |
| Bacteria-Proteobacteria-Gammaproteobacteria-Enterobacteriales-Enterobacteriaceae-Klebsiella-Klebsiella_unclassified | 0.015 | 275 |
| Bacteria-Proteobacteria-Alphaproteobacteria-Rhizobiales-Rhizobiaceae-Sinorhizobium-Sinorhizobium_unclassified | 0.014 | 252 |
| Bacteria-Proteobacteria-Betaproteobacteria-Rhodocyclales-Rhodocyclaceae-Azospira-Azospira_oryzae | 0.012 | 147 |
| Bacteria-Proteobacteria-Gammaproteobacteria-Methylococcales-Methylococcaceae-Methylobacter-Methylobacter_unclassified | 0.011 | 106 |
| Bacteria-Proteobacteria-Alphaproteobacteria-Rickettsiales-Holosporaceae-Holospora-Holospora_unclassified | 0.010 | 47 |
| Bacteria-Firmicutes-Clostridia-Clostridiales-Peptostreptococcaceae-Peptostreptococcaceae_noname-Clostridium_bifermentans | 0.010 | 119 |
| Bacteria-Proteobacteria-Deltaproteobacteria-Myxococcales-Myxococcaceae-Corallococcus-Corallococcus_coralloides | 0.007 | 245 |
| Bacteria-Bacteroidetes-Cytophagia-Cytophagales-Cytophagaceae-Hymenobacter-Hymenobacter_unclassified | 0.007 | 113 |
| Bacteria-Actinobacteria-Actinobacteria-Actinomycetales-Dietziaceae-Dietzia-Dietzia_cinnamea | 0.005 | 54 |
| Bacteria-Proteobacteria-Alphaproteobacteria-Rhizobiales-Rhizobiaceae-Kaistia-Kaistia_granuli | 0.004 | 58 |
| Bacteria-Proteobacteria-Alphaproteobacteria-Rhizobiales-Rhizobiaceae-Agrobacterium-Agrobacterium_albertimagni | 0.003 | 56 |
| Bacteria-Bacteroidetes-Cytophagia-Cytophagales-Cytophagaceae-Pontibacter-Pontibacter_unclassified | 0.003 | 44 |
| Bacteria-Proteobacteria-Betaproteobacteria-Burkholderiales-Burkholderiales_noname-Burkholderiales_noname-Burkholderiales_bacterium_JOSHI_001 | 0.003 | 52 |
| Bacteria-Proteobacteria-Alphaproteobacteria-Sphingomonadales-Sphingomonadaceae-Blastomonas-Blastomonas_unclassified | 0.003 | 35 |

**Table S5**. Table showing number of reads and relative abundance assigned to 20 different ARG classes (types) discovered in the Ganges sediments

| **ARG Classes (ARG types)** | **No. of Reads**  **(21869)** | **Abundance**  **(%)** | **Relative abundance (%)** |
| --- | --- | --- | --- |
| Multidrug resistance (MDR) | 12041 | 0.003824 | 55.06 |
| Bacitracins | 3202 | 0.001017 | 14.64 |
| Macrolide-lincosamide-streptogramin (MLS) | 1744 | 0.000554 | 7.97 |
| Fosmidomycins | 990 | 0.000314 | 4.53 |
| Unclassified | 777 | 0.000247 | 3.55 |
| Vancomycins | 684 | 0.000217 | 3.13 |
| Tetracyclines | 376 | 0.000119 | 1.72 |
| Quinolones | 364 | 0.000116 | 1.66 |
| Aminoglycoside | 355 | 0.000113 | 1.62 |
| Rifamycins | 300 | 0.000095 | 1.37 |
| Trimethoprim | 255 | 0.000081 | 1.17 |
| Betalactams | 254 | 0.000081 | 1.16 |
| Sulfonamide | 204 | 0.000065 | 0.93 |
| Chloramphenicol | 148 | 0.000047 | 0.68 |
| Polymyxin | 78 | 0.000025 | 0.36 |
| Fosfomycin | 45 | 0.000014 | 0.21 |
| Tetracenomycin_C | 34 | 0.000011 | 0.16 |
| Kasugamycin | 6 | 0.000002 | 0.03 |
| Puromycin | 6 | 0.000002 | 0.03 |
| Carbomycin | 6 | 0.000002 | 0.03 |

**Table S6.** Alphabetical list of all the identified ARG classes (types) and genes (subtypes) along with corresponding abundance and number of reads

| Classes (Types) | Genes (Subtypes) | Abundance (ppm) | Number of Reads | Relative abundance (%) |
| --- | --- | --- | --- | --- |
| Aminoglycoside (22) | aadA | 0.311 | 98 | 0.448 |
|  | aph(6)-I | 0.171 | 54 | 0.247 |
|  | aph(3')-I | 0.137 | 43 | 0.197 |
|  | aac(3)-I | 0.124 | 39 | 0.178 |
|  | aac(3)-IV | 0.057 | 18 | 0.082 |
|  | aac(6')-II | 0.051 | 16 | 0.073 |
|  | aadE | 0.051 | 16 | 0.073 |
|  | ant(9)-I | 0.051 | 16 | 0.073 |
|  | aac(3)-IIIa | 0.032 | 10 | 0.046 |
|  | aac(6')-I | 0.032 | 10 | 0.046 |
|  | aac(2')-I | 0.022 | 7 | 0.032 |
|  | aph(3''')-III | 0.016 | 5 | 0.023 |
|  | aph(3')-V | 0.013 | 4 | 0.018 |
|  | aac(3)-VII | 0.010 | 3 | 0.014 |
|  | aac(3)-X | 0.010 | 3 | 0.014 |
|  | aac(6')-31 | 0.010 | 3 | 0.014 |
|  | ant(2'')-I | 0.010 | 3 | 0.014 |
|  | aph(2'')-III | 0.010 | 3 | 0.014 |
|  | aac(3)-II | 0.003 | 1 | 0.005 |
|  | aac(3)-VIII | 0.003 | 1 | 0.005 |
|  | aad(6) | 0.003 | 1 | 0.005 |
|  | tunicamycin resistance protein | 0.003 | 1 | 0.005 |
| Bacitracin (2) | bacA | 9.988 | 3145 | 14.381 |
|  | bcrA | 0.181 | 57 | 0.261 |
| Betalactams (59) | THIN-B | 0.076 | 24 | 0.110 |
|  | OXA-209 | 0.051 | 16 | 0.073 |
|  | OXA-36 | 0.041 | 13 | 0.059 |
|  | class A beta-lactamase | 0.041 | 13 | 0.059 |
|  | OXA-53 | 0.038 | 12 | 0.055 |
|  | CMY-19 | 0.032 | 10 | 0.046 |
|  | LCR-1 | 0.032 | 10 | 0.046 |
|  | OXA-119 | 0.032 | 10 | 0.046 |
|  | OXA-205 | 0.032 | 10 | 0.046 |
|  | NPS-1 | 0.029 | 9 | 0.041 |
|  | ampC | 0.029 | 9 | 0.041 |
|  | OXA-21 | 0.025 | 8 | 0.037 |
|  | MOX-4 | 0.022 | 7 | 0.032 |
|  | OXA-118 | 0.022 | 7 | 0.032 |
|  | OXA-46 | 0.022 | 7 | 0.032 |
|  | GES-15 | 0.019 | 6 | 0.027 |
|  | FEZ-1 | 0.013 | 4 | 0.018 |
|  | OXA-142 | 0.013 | 4 | 0.018 |
|  | OXA-251 | 0.013 | 4 | 0.018 |
|  | OXA-34 | 0.013 | 4 | 0.018 |
|  | OXA-37 | 0.013 | 4 | 0.018 |
|  | TEM-117 | 0.013 | 4 | 0.018 |
|  | MOX-3 | 0.010 | 3 | 0.014 |
|  | OXA-10 | 0.010 | 3 | 0.014 |
|  | OXA-2 | 0.010 | 3 | 0.014 |
|  | CMY-9 | 0.006 | 2 | 0.009 |
|  | GES-14 | 0.006 | 2 | 0.009 |
|  | MOX-2 | 0.006 | 2 | 0.009 |
|  | OXA-129 | 0.006 | 2 | 0.009 |
|  | OXA-141 | 0.006 | 2 | 0.009 |
|  | OXA-147 | 0.006 | 2 | 0.009 |
|  | OXA-20 | 0.006 | 2 | 0.009 |
|  | OXA-226 | 0.006 | 2 | 0.009 |
|  | OXA-3 | 0.006 | 2 | 0.009 |
|  | PBP-1A | 0.006 | 2 | 0.009 |
|  | TEM-171 | 0.006 | 2 | 0.009 |
|  | VEB-1 | 0.006 | 2 | 0.009 |
|  | VEB-6 | 0.006 | 2 | 0.009 |
|  | class B beta-lactamase | 0.006 | 2 | 0.009 |
|  | class C beta-lactamase | 0.006 | 2 | 0.009 |
|  | metallo-beta-lactamase | 0.006 | 2 | 0.009 |
|  | CMY-1 | 0.003 | 1 | 0.005 |
|  | GES-13 | 0.003 | 1 | 0.005 |
|  | GES-4 | 0.003 | 1 | 0.005 |
|  | LRA-12 | 0.003 | 1 | 0.005 |
|  | LRA-19 | 0.003 | 1 | 0.005 |
|  | LRA-5 | 0.003 | 1 | 0.005 |
|  | LRA-9 | 0.003 | 1 | 0.005 |
|  | OCH-8 | 0.003 | 1 | 0.005 |
|  | OXA-18 | 0.003 | 1 | 0.005 |
|  | OXA-183 | 0.003 | 1 | 0.005 |
|  | OXA-5 | 0.003 | 1 | 0.005 |
|  | OXA-7 | 0.003 | 1 | 0.005 |
|  | OXA-74 | 0.003 | 1 | 0.005 |
|  | TEM-1 | 0.003 | 1 | 0.005 |
|  | TEM-87 | 0.003 | 1 | 0.005 |
|  | VEB-2 | 0.003 | 1 | 0.005 |
|  | VEB-7 | 0.003 | 1 | 0.005 |
|  | penA | 0.003 | 1 | 0.005 |
| Carbomycin (1) | carA | 0.019 | 6 | 0.027 |
| Chloramphenicol (8) | catB | 0.133 | 42 | 0.192 |
|  | cat_chloramphenicol acetyltransferase | 0.133 | 42 | 0.192 |
|  | chloramphenicol exporter | 0.092 | 29 | 0.133 |
|  | catQ | 0.044 | 14 | 0.064 |
|  | cmlA | 0.032 | 10 | 0.046 |
|  | cmrA | 0.025 | 8 | 0.037 |
|  | catA | 0.006 | 2 | 0.009 |
|  | chloramphenicol and florfenicol exporter | 0.003 | 1 | 0.005 |
| Fosfomycin (2) | fosB | 0.130 | 41 | 0.187 |
|  | fosX | 0.013 | 4 | 0.018 |
| Fosmidomycin (2) | rosA | 0.632 | 199 | 0.910 |
|  | rosB | 2.512 | 791 | 3.617 |
| kasugamycin (1) | kasugamycin resistance protein ksgA | 0.019 | 6 | 0.027 |
| Macrolide-Lincosamide-Streptogramin (MLS) (28) | macB | 4.513 | 1421 | 6.498 |
|  | mefA | 0.521 | 164 | 0.750 |
|  | ermF | 0.127 | 40 | 0.183 |
|  | ereA | 0.067 | 21 | 0.096 |
|  | macA | 0.054 | 17 | 0.078 |
|  | lnuA | 0.038 | 12 | 0.055 |
|  | vatB | 0.032 | 10 | 0.046 |
|  | lsa | 0.022 | 7 | 0.032 |
|  | lmrB | 0.019 | 6 | 0.027 |
|  | lnuB | 0.016 | 5 | 0.023 |
|  | mphA | 0.016 | 5 | 0.023 |
|  | oleD | 0.016 | 5 | 0.023 |
|  | tlcC | 0.016 | 5 | 0.023 |
|  | srmB | 0.013 | 4 | 0.018 |
|  | erm(35) | 0.010 | 3 | 0.014 |
|  | ereB | 0.006 | 2 | 0.009 |
|  | ermA | 0.006 | 2 | 0.009 |
|  | ermO | 0.006 | 2 | 0.009 |
|  | ermX | 0.006 | 2 | 0.009 |
|  | mgtA | 0.006 | 2 | 0.009 |
|  | vatG | 0.006 | 2 | 0.009 |
|  | erm(33) | 0.003 | 1 | 0.005 |
|  | erm(38) | 0.003 | 1 | 0.005 |
|  | erm(39) | 0.003 | 1 | 0.005 |
|  | erm(TR) | 0.003 | 1 | 0.005 |
|  | ermC | 0.003 | 1 | 0.005 |
|  | vatD | 0.003 | 1 | 0.005 |
|  | vgbA | 0.003 | 1 | 0.005 |
| Multidrug (MDR) (59) | mexF | 7.752 | 2441 | 11.162 |
|  | acrB | 5.189 | 1634 | 7.472 |
|  | multidrug_ABC_transporter | 3.389 | 1067 | 4.879 |
|  | mdtB | 3.074 | 968 | 4.426 |
|  | multidrug_transporter | 2.699 | 850 | 3.887 |
|  | mexW | 2.131 | 671 | 3.068 |
|  | ceoB | 1.956 | 616 | 2.817 |
|  | mdtC | 1.839 | 579 | 2.648 |
|  | smeE | 1.083 | 341 | 1.559 |
|  | mexI | 0.934 | 294 | 1.344 |
|  | mexB | 0.921 | 290 | 1.326 |
|  | amrB | 0.848 | 267 | 1.221 |
|  | mexD | 0.673 | 212 | 0.969 |
|  | ompR | 0.635 | 200 | 0.915 |
|  | acrF | 0.540 | 170 | 0.777 |
|  | oprC | 0.511 | 161 | 0.736 |
|  | mexY | 0.413 | 130 | 0.594 |
|  | major_facilitator_superfamily_transporter | 0.406 | 128 | 0.585 |
|  | bpeF | 0.403 | 127 | 0.581 |
|  | mdtF | 0.289 | 91 | 0.416 |
|  | adeB | 0.245 | 77 | 0.352 |
|  | mexE | 0.232 | 73 | 0.334 |
|  | mexC | 0.216 | 68 | 0.311 |
|  | oprM | 0.203 | 64 | 0.293 |
|  | adeJ | 0.191 | 60 | 0.274 |
|  | mexT | 0.187 | 59 | 0.270 |
|  | smeB | 0.178 | 56 | 0.256 |
|  | smeD | 0.162 | 51 | 0.233 |
|  | cmeB | 0.152 | 48 | 0.219 |
|  | oprN | 0.121 | 38 | 0.174 |
|  | emrB | 0.089 | 28 | 0.128 |
|  | mexA | 0.067 | 21 | 0.096 |
|  | mexX | 0.054 | 17 | 0.078 |
|  | EmrB-QacA family major facilitator transporter | 0.048 | 15 | 0.069 |
|  | smeF | 0.048 | 15 | 0.069 |
|  | acrA | 0.044 | 14 | 0.064 |
|  | oprA | 0.044 | 14 | 0.064 |
|  | oprJ | 0.035 | 11 | 0.050 |
|  | mdtK | 0.029 | 9 | 0.041 |
|  | sdeY | 0.029 | 9 | 0.041 |
|  | adeC | 0.022 | 7 | 0.032 |
|  | TolC | 0.019 | 6 | 0.027 |
|  | mdfA | 0.016 | 5 | 0.023 |
|  | adeK | 0.013 | 4 | 0.018 |
|  | bicyclomycin-multidrug_efflux_protein_bcr | 0.013 | 4 | 0.018 |
|  | emrA | 0.013 | 4 | 0.018 |
|  | ompF | 0.013 | 4 | 0.018 |
|  | emrD | 0.010 | 3 | 0.014 |
|  | mdtA | 0.010 | 3 | 0.014 |
|  | abeS | 0.006 | 2 | 0.009 |
|  | mdtD | 0.006 | 2 | 0.009 |
|  | mdtE | 0.006 | 2 | 0.009 |
|  | mdtG | 0.006 | 2 | 0.009 |
|  | mdtH | 0.006 | 2 | 0.009 |
|  | mdtL | 0.006 | 2 | 0.009 |
|  | omp36 | 0.006 | 2 | 0.009 |
|  | mexH | 0.003 | 1 | 0.005 |
|  | opcM | 0.003 | 1 | 0.005 |
|  | smeC | 0.003 | 1 | 0.005 |
| Polymyxin (1) | arnA | 0.248 | 78 | 0.357 |
| Puromycin (1) | puromycin resistance protein | 0.019 | 6 | 0.027 |
| Quinolone (3) | norB | 0.546 | 172 | 0.787 |
|  | qepA | 0.603 | 190 | 0.869 |
|  | qnrS | 0.006 | 2 | 0.009 |
| Rifamycin (2) | ADP-ribosylating transferase_arr | 0.680 | 214 | 0.979 |
|  | rifampin monooxygenase | 0.273 | 86 | 0.393 |
| Sulfonamide (3) | sul1 | 0.400 | 126 | 0.576 |
|  | sul2 | 0.245 | 77 | 0.352 |
|  | sul3 | 0.003 | 1 | 0.005 |
| Tetracenomycin_C (1) | tcmA | 0.108 | 34 | 0.155 |
| Tetracycline (20) | tetP | 0.441 | 139 | 0.636 |
|  | otrA | 0.184 | 58 | 0.265 |
|  | tetM | 0.114 | 36 | 0.165 |
|  | tet35 | 0.111 | 35 | 0.160 |
|  | tetA | 0.079 | 25 | 0.114 |
|  | tetV | 0.048 | 15 | 0.069 |
|  | tetW | 0.048 | 15 | 0.069 |
|  | tetC | 0.038 | 12 | 0.055 |
|  | tetG | 0.029 | 9 | 0.041 |
|  | tetR | 0.022 | 7 | 0.032 |
|  | tet41 | 0.016 | 5 | 0.023 |
|  | tetracycline_resistance_protein | 0.013 | 4 | 0.018 |
|  | tetQ | 0.010 | 3 | 0.014 |
|  | tet32 | 0.006 | 2 | 0.009 |
|  | tet36 | 0.006 | 2 | 0.009 |
|  | tet39 | 0.006 | 2 | 0.009 |
|  | tet40 | 0.006 | 2 | 0.009 |
|  | tet44 | 0.006 | 2 | 0.009 |
|  | tetO | 0.006 | 2 | 0.009 |
|  | tetT | 0.003 | 1 | 0.005 |
| Trimethoprim (8) | dfrA12 | 0.718 | 226 | 1.033 |
|  | dfrA23 | 0.038 | 12 | 0.055 |
|  | dfrA1 | 0.022 | 7 | 0.032 |
|  | dfrA13 | 0.016 | 5 | 0.023 |
|  | dfrB3 | 0.006 | 2 | 0.009 |
|  | dfrA22 | 0.003 | 1 | 0.005 |
|  | dfrA5 | 0.003 | 1 | 0.005 |
|  | dfrB1 | 0.003 | 1 | 0.005 |
| Unclassified (4) | transcriptional regulatory protein CpxR cpxR | 1.458 | 459 | 2.099 |
|  | truncated putative response regulator ArlR | 0.692 | 218 | 0.997 |
|  | cAMP-regulatory protein | 0.181 | 57 | 0.261 |
|  | histidine kinase | 0.137 | 43 | 0.197 |
| Vancomycin (13) | vanR | 1.420 | 447 | 2.044 |
|  | vanS | 0.489 | 154 | 0.704 |
|  | vanH | 0.057 | 18 | 0.082 |
|  | vanA | 0.044 | 14 | 0.064 |
|  | vanD | 0.041 | 13 | 0.059 |
|  | vanX | 0.035 | 11 | 0.050 |
|  | vanC | 0.032 | 10 | 0.046 |
|  | vanE | 0.016 | 5 | 0.023 |
|  | vanW | 0.013 | 4 | 0.018 |
|  | vanG | 0.010 | 3 | 0.014 |
|  | vanB | 0.006 | 2 | 0.009 |
|  | vanY | 0.006 | 2 | 0.009 |
|  | vanU | 0.003 | 1 | 0.005 |

**Table S7.** Abundance wise catalogue of antibiotic resistance genes

| **Genes (Subtypes)** | **Class (Types)** | **Abundance (ppm)** | **Number of Reads** | **Relative abundance (%)** |
| --- | --- | --- | --- | --- |
| bacA | Bacitracin | 9.988 | 3145 | 14.381 |
| mexF | MDR | 7.752 | 2441 | 11.162 |
| acrB | MDR | 5.189 | 1634 | 7.472 |
| macB | MLS | 4.513 | 1421 | 6.498 |
| multidrug_ABC_transporter | MDR | 3.389 | 1067 | 4.879 |
| mdtB | MDR | 3.074 | 968 | 4.426 |
| multidrug_transporter | MDR | 2.699 | 850 | 3.887 |
| rosB | Fosmidomycin | 2.512 | 791 | 3.617 |
| mexW | MDR | 2.131 | 671 | 3.068 |
| ceoB | MDR | 1.956 | 616 | 2.817 |
| mdtC | MDR | 1.839 | 579 | 2.648 |
| transcriptional regulatory protein CpxR cpxR | Unclassified | 1.458 | 459 | 2.099 |
| vanR | Vancomycin | 1.420 | 447 | 2.044 |
| smeE | MDR | 1.083 | 341 | 1.559 |
| mexI | MDR | 0.934 | 294 | 1.344 |
| mexB | MDR | 0.921 | 290 | 1.326 |
| amrB | MDR | 0.848 | 267 | 1.221 |
| dfrA12 | Trimethoprim | 0.718 | 226 | 1.033 |
| truncated putative response regulator ArlR | Unclassified | 0.692 | 218 | 0.997 |
| ADP-ribosylating transferase_arr | Rifamycin | 0.680 | 214 | 0.979 |
| mexD | MDR | 0.673 | 212 | 0.969 |
| ompR | MDR | 0.635 | 200 | 0.915 |
| rosA | Fosmidomycin | 0.632 | 199 | 0.910 |
| qepA | Quinolone | 0.603 | 190 | 0.869 |
| norB | Quinolone | 0.546 | 172 | 0.787 |
| acrF | MDR | 0.540 | 170 | 0.777 |
| mefA | MLS | 0.521 | 164 | 0.750 |
| oprC | MDR | 0.511 | 161 | 0.736 |
| vanS | Vancomycin | 0.489 | 154 | 0.704 |
| tetP | Tetracycline | 0.441 | 139 | 0.636 |
| mexY | MDR | 0.413 | 130 | 0.594 |
| major_facilitator_superfamily_transporter (MFS) | MDR | 0.406 | 128 | 0.585 |
| bpeF | MDR | 0.403 | 127 | 0.581 |
| sul1 | Sulfonamide | 0.400 | 126 | 0.576 |
| aadA | Aminoglycoside | 0.311 | 98 | 0.448 |
| mdtF | MDR | 0.289 | 91 | 0.416 |
| rifampin monooxygenase | Rifamycin | 0.273 | 86 | 0.393 |
| arnA | Polymyxin | 0.248 | 78 | 0.357 |
| adeB | MDR | 0.245 | 77 | 0.352 |
| sul2 | Sulfonamide | 0.245 | 77 | 0.352 |
| mexE | MDR | 0.232 | 73 | 0.334 |
| mexC | MDR | 0.216 | 68 | 0.311 |
| oprM | MDR | 0.203 | 64 | 0.293 |
| adeJ | MDR | 0.191 | 60 | 0.274 |
| mexT | MDR | 0.187 | 59 | 0.270 |
| otrA | Tetracycline | 0.184 | 58 | 0.265 |
| bcrA | Bacitracin | 0.181 | 57 | 0.261 |
| cAMP-regulatory protein | Unclassified | 0.181 | 57 | 0.261 |
| smeB | MDR | 0.178 | 56 | 0.256 |
| aph(6)-I | Aminoglycoside | 0.171 | 54 | 0.247 |
| smeD | MDR | 0.162 | 51 | 0.233 |
| cmeB | MDR | 0.152 | 48 | 0.219 |
| aph(3')-I | Aminoglycoside | 0.137 | 43 | 0.197 |
| histidine kinase | Unclassified | 0.137 | 43 | 0.197 |
| catB | Chloramphenicol | 0.133 | 42 | 0.192 |
| cat_chloramphenicol acetyltransferase | Chloramphenicol | 0.133 | 42 | 0.192 |
| fosB | Fosfomycin | 0.130 | 41 | 0.187 |
| ermF | MLS | 0.127 | 40 | 0.183 |
| aac(3)-I | Aminoglycoside | 0.124 | 39 | 0.178 |
| oprN | MDR | 0.121 | 38 | 0.174 |
| tetM | Tetracycline | 0.114 | 36 | 0.165 |
| tet35 | Tetracycline | 0.111 | 35 | 0.160 |
| tcmA | Tetracenomycin_C | 0.108 | 34 | 0.155 |
| chloramphenicol exporter | Chloramphenicol | 0.092 | 29 | 0.133 |
| emrB | MDR | 0.089 | 28 | 0.128 |
| tetA | Tetracycline | 0.079 | 25 | 0.114 |
| THIN-B | Betalactams | 0.076 | 24 | 0.110 |
| ereA | MLS | 0.067 | 21 | 0.096 |
| mexA | MDR | 0.067 | 21 | 0.096 |
| aac(3)-IV | Aminoglycoside | 0.057 | 18 | 0.082 |
| vanH | Vancomycin | 0.057 | 18 | 0.082 |
| macA | MLS | 0.054 | 17 | 0.078 |
| mexX | MDR | 0.054 | 17 | 0.078 |
| aac(6')-II | Aminoglycoside | 0.051 | 16 | 0.073 |
| aadE | Aminoglycoside | 0.051 | 16 | 0.073 |
| ant(9)-I | Aminoglycoside | 0.051 | 16 | 0.073 |
| OXA-209 | Betalactams | 0.051 | 16 | 0.073 |
| EmrB-QacA family major facilitator transporter | MDR | 0.048 | 15 | 0.069 |
| smeF | MDR | 0.048 | 15 | 0.069 |
| tetV | Tetracycline | 0.048 | 15 | 0.069 |
| tetW | Tetracycline | 0.048 | 15 | 0.069 |
| catQ | Chloramphenicol | 0.044 | 14 | 0.064 |
| acrA | MDR | 0.044 | 14 | 0.064 |
| oprA | MDR | 0.044 | 14 | 0.064 |
| vanA | Vancomycin | 0.044 | 14 | 0.064 |
| OXA-36 | Betalactams | 0.041 | 13 | 0.059 |
| class A beta-lactamase | Betalactams | 0.041 | 13 | 0.059 |
| vanD | Vancomycin | 0.041 | 13 | 0.059 |
| OXA-53 | Betalactams | 0.038 | 12 | 0.055 |
| lnuA | MLS | 0.038 | 12 | 0.055 |
| tetC | Tetracycline | 0.038 | 12 | 0.055 |
| dfrA23 | Trimethoprim | 0.038 | 12 | 0.055 |
| oprJ | MDR | 0.035 | 11 | 0.050 |
| vanX | Vancomycin | 0.035 | 11 | 0.050 |
| aac(3)-IIIa | Aminoglycoside | 0.032 | 10 | 0.046 |
| aac(6')-I | Aminoglycoside | 0.032 | 10 | 0.046 |
| CMY-19 | Betalactams | 0.032 | 10 | 0.046 |
| LCR-1 | Betalactams | 0.032 | 10 | 0.046 |
| OXA-119 | Betalactams | 0.032 | 10 | 0.046 |
| OXA-205 | Betalactams | 0.032 | 10 | 0.046 |
| cmlA | Chloramphenicol | 0.032 | 10 | 0.046 |
| vatB | MLS | 0.032 | 10 | 0.046 |
| vanC | Vancomycin | 0.032 | 10 | 0.046 |
| NPS-1 | Betalactams | 0.029 | 9 | 0.041 |
| ampC | Betalactams | 0.029 | 9 | 0.041 |
| mdtK | MDR | 0.029 | 9 | 0.041 |
| sdeY | MDR | 0.029 | 9 | 0.041 |
| tetG | Tetracycline | 0.029 | 9 | 0.041 |
| OXA-21 | Betalactams | 0.025 | 8 | 0.037 |
| cmrA | Chloramphenicol | 0.025 | 8 | 0.037 |
| aac(2')-I | Aminoglycoside | 0.022 | 7 | 0.032 |
| MOX-4 | Betalactams | 0.022 | 7 | 0.032 |
| OXA-118 | Betalactams | 0.022 | 7 | 0.032 |
| OXA-46 | Betalactams | 0.022 | 7 | 0.032 |
| lsa | MLS | 0.022 | 7 | 0.032 |
| adeC | MDR | 0.022 | 7 | 0.032 |
| tetR | Tetracycline | 0.022 | 7 | 0.032 |
| dfrA1 | Trimethoprim | 0.022 | 7 | 0.032 |
| GES-15 | Betalactams | 0.019 | 6 | 0.027 |
| carA | Carbomycin | 0.019 | 6 | 0.027 |
| kasugamycin resistance protein ksgA | kasugamycin | 0.019 | 6 | 0.027 |
| lmrB | MLS | 0.019 | 6 | 0.027 |
| TolC | MDR | 0.019 | 6 | 0.027 |
| puromycin resistance protein | Puromycin | 0.019 | 6 | 0.027 |
| aph(3''')-III | Aminoglycoside | 0.016 | 5 | 0.023 |
| lnuB | MLS | 0.016 | 5 | 0.023 |
| mphA | MLS | 0.016 | 5 | 0.023 |
| oleD | MLS | 0.016 | 5 | 0.023 |
| tlcC | MLS | 0.016 | 5 | 0.023 |
| mdfA | MDR | 0.016 | 5 | 0.023 |
| tet41 | Tetracycline | 0.016 | 5 | 0.023 |
| dfrA13 | Trimethoprim | 0.016 | 5 | 0.023 |
| vanE | Vancomycin | 0.016 | 5 | 0.023 |
| aph(3')-V | Aminoglycoside | 0.013 | 4 | 0.018 |
| FEZ-1 | Betalactams | 0.013 | 4 | 0.018 |
| OXA-142 | Betalactams | 0.013 | 4 | 0.018 |
| OXA-251 | Betalactams | 0.013 | 4 | 0.018 |
| OXA-34 | Betalactams | 0.013 | 4 | 0.018 |
| OXA-37 | Betalactams | 0.013 | 4 | 0.018 |
| TEM-117 | Betalactams | 0.013 | 4 | 0.018 |
| fosX | Fosfomycin | 0.013 | 4 | 0.018 |
| srmB | MLS | 0.013 | 4 | 0.018 |
| adeK | MDR | 0.013 | 4 | 0.018 |
| bicyclomycin-multidrug_efflux_protein_bcr | MDR | 0.013 | 4 | 0.018 |
| emrA | MDR | 0.013 | 4 | 0.018 |
| ompF | MDR | 0.013 | 4 | 0.018 |
| tetracycline_resistance_protein | Tetracycline | 0.013 | 4 | 0.018 |
| vanW | Vancomycin | 0.013 | 4 | 0.018 |
| aac(3)-VII | Aminoglycoside | 0.010 | 3 | 0.014 |
| aac(3)-X | Aminoglycoside | 0.010 | 3 | 0.014 |
| aac(6')-31 | Aminoglycoside | 0.010 | 3 | 0.014 |
| ant(2'')-I | Aminoglycoside | 0.010 | 3 | 0.014 |
| aph(2'')-III | Aminoglycoside | 0.010 | 3 | 0.014 |
| MOX-3 | Betalactams | 0.010 | 3 | 0.014 |
| OXA-10 | Betalactams | 0.010 | 3 | 0.014 |
| OXA-2 | Betalactams | 0.010 | 3 | 0.014 |
| erm(35) | MLS | 0.010 | 3 | 0.014 |
| emrD | MDR | 0.010 | 3 | 0.014 |
| mdtA | MDR | 0.010 | 3 | 0.014 |
| tetQ | Tetracycline | 0.010 | 3 | 0.014 |
| vanG | Vancomycin | 0.010 | 3 | 0.014 |
| CMY-9 | Betalactams | 0.006 | 2 | 0.009 |
| GES-14 | Betalactams | 0.006 | 2 | 0.009 |
| MOX-2 | Betalactams | 0.006 | 2 | 0.009 |
| OXA-129 | Betalactams | 0.006 | 2 | 0.009 |
| OXA-141 | Betalactams | 0.006 | 2 | 0.009 |
| OXA-147 | Betalactams | 0.006 | 2 | 0.009 |
| OXA-20 | Betalactams | 0.006 | 2 | 0.009 |
| OXA-226 | Betalactams | 0.006 | 2 | 0.009 |
| OXA-3 | Betalactams | 0.006 | 2 | 0.009 |
| PBP-1A | Betalactams | 0.006 | 2 | 0.009 |
| TEM-171 | Betalactams | 0.006 | 2 | 0.009 |
| VEB-1 | Betalactams | 0.006 | 2 | 0.009 |
| VEB-6 | Betalactams | 0.006 | 2 | 0.009 |
| class B beta-lactamase | Betalactams | 0.006 | 2 | 0.009 |
| class C beta-lactamase | Betalactams | 0.006 | 2 | 0.009 |
| metallo-beta-lactamase | Betalactams | 0.006 | 2 | 0.009 |
| catA | Chloramphenicol | 0.006 | 2 | 0.009 |
| ereB | MLS | 0.006 | 2 | 0.009 |
| ermA | MLS | 0.006 | 2 | 0.009 |
| ermO | MLS | 0.006 | 2 | 0.009 |
| ermX | MLS | 0.006 | 2 | 0.009 |
| mgtA | MLS | 0.006 | 2 | 0.009 |
| vatG | MLS | 0.006 | 2 | 0.009 |
| abeS | MDR | 0.006 | 2 | 0.009 |
| mdtD | MDR | 0.006 | 2 | 0.009 |
| mdtE | MDR | 0.006 | 2 | 0.009 |
| mdtG | MDR | 0.006 | 2 | 0.009 |
| mdtH | MDR | 0.006 | 2 | 0.009 |
| mdtL | MDR | 0.006 | 2 | 0.009 |
| omp36 | MDR | 0.006 | 2 | 0.009 |
| qnrS | Quinolone | 0.006 | 2 | 0.009 |
| tet32 | Tetracycline | 0.006 | 2 | 0.009 |
| tet36 | Tetracycline | 0.006 | 2 | 0.009 |
| tet39 | Tetracycline | 0.006 | 2 | 0.009 |
| tet40 | Tetracycline | 0.006 | 2 | 0.009 |
| tet44 | Tetracycline | 0.006 | 2 | 0.009 |
| tetO | Tetracycline | 0.006 | 2 | 0.009 |
| dfrB3 | Trimethoprim | 0.006 | 2 | 0.009 |
| vanB | Vancomycin | 0.006 | 2 | 0.009 |
| vanY | Vancomycin | 0.006 | 2 | 0.009 |
| aac(3)-II | Aminoglycoside | 0.003 | 1 | 0.005 |
| aac(3)-VIII | Aminoglycoside | 0.003 | 1 | 0.005 |
| aad(6) | Aminoglycoside | 0.003 | 1 | 0.005 |
| tunicamycin resistance protein | Aminoglycoside | 0.003 | 1 | 0.005 |
| CMY-1 | Betalactams | 0.003 | 1 | 0.005 |
| GES-13 | Betalactams | 0.003 | 1 | 0.005 |
| GES-4 | Betalactams | 0.003 | 1 | 0.005 |
| LRA-12 | Betalactams | 0.003 | 1 | 0.005 |
| LRA-19 | Betalactams | 0.003 | 1 | 0.005 |
| LRA-5 | Betalactams | 0.003 | 1 | 0.005 |
| LRA-9 | Betalactams | 0.003 | 1 | 0.005 |
| OCH-8 | Betalactams | 0.003 | 1 | 0.005 |
| OXA-18 | Betalactams | 0.003 | 1 | 0.005 |
| OXA-183 | Betalactams | 0.003 | 1 | 0.005 |
| OXA-5 | Betalactams | 0.003 | 1 | 0.005 |
| OXA-7 | Betalactams | 0.003 | 1 | 0.005 |
| OXA-74 | Betalactams | 0.003 | 1 | 0.005 |
| TEM-1 | Betalactams | 0.003 | 1 | 0.005 |
| TEM-87 | Betalactams | 0.003 | 1 | 0.005 |
| VEB-2 | Betalactams | 0.003 | 1 | 0.005 |
| VEB-7 | Betalactams | 0.003 | 1 | 0.005 |
| penA | Betalactams | 0.003 | 1 | 0.005 |
| chloramphenicol and florfenicol exporte | Chloramphenicol | 0.003 | 1 | 0.005 |
| erm(33) | MLS | 0.003 | 1 | 0.005 |
| erm(38) | MLS | 0.003 | 1 | 0.005 |
| erm(39) | MLS | 0.003 | 1 | 0.005 |
| erm(TR) | MLS | 0.003 | 1 | 0.005 |
| ermC | MLS | 0.003 | 1 | 0.005 |
| vatD | MLS | 0.003 | 1 | 0.005 |
| vgbA | MLS | 0.003 | 1 | 0.005 |
| mexH | MDR | 0.003 | 1 | 0.005 |
| opcM | MDR | 0.003 | 1 | 0.005 |
| smeC | MDR | 0.003 | 1 | 0.005 |
| sul3 | Sulfonamide | 0.003 | 1 | 0.005 |
| tetT | Tetracycline | 0.003 | 1 | 0.005 |
| dfrA22 | Trimethoprim | 0.003 | 1 | 0.005 |
| dfrA5 | Trimethoprim | 0.003 | 1 | 0.005 |
| dfrB1 | Trimethoprim | 0.003 | 1 | 0.005 |
| vanU | Vancomycin | 0.003 | 1 | 0.005 |

MDR, Multidrug resistance; MLS, Macrolide-Lincosamide-Streptogramin

**Table S8.** Distribution and abundance of the antibiotic resistance mechanisms

| **Mechanism** | **Genes (Subtypes)** | **Abundance (ppm)** | **Number of Reads** | **Relative abundance (%)** |
| --- | --- | --- | --- | --- |
| Antibiotic efflux | mexF-MDR | 7.752 | 2441 | 11.162 |
|  | acrB-MDR | 5.189 | 1634 | 7.472 |
|  | macB-MLS | 4.513 | 1421 | 6.498 |
|  | multidrug_ABC_transporter-MDR | 3.389 | 1067 | 4.879 |
|  | mdtB-MDR | 3.074 | 968 | 4.426 |
|  | multidrug_transporter-MDR | 2.699 | 850 | 3.887 |
|  | rosB-Fosmidomycin | 2.512 | 791 | 3.617 |
|  | mexW-MDR | 2.131 | 671 | 3.068 |
|  | ceoB-MDR | 1.956 | 616 | 2.817 |
|  | mdtC-MDR | 1.839 | 579 | 2.648 |
|  | transcriptional regulatory protein CpxR cpxR-Unclassified | 1.458 | 459 | 2.099 |
|  | smeE-MDR | 1.083 | 341 | 1.559 |
|  | mexI-MDR | 0.934 | 294 | 1.344 |
|  | mexB-MDR | 0.921 | 290 | 1.326 |
|  | amrB-MDR | 0.848 | 267 | 1.221 |
|  | truncated putative response regulator ArlR-Unclassified | 0.692 | 218 | 0.997 |
|  | mexD-MDR | 0.673 | 212 | 0.969 |
|  | rosA-Fosmidomycin | 0.632 | 199 | 0.910 |
|  | qepA-Quinolone | 0.603 | 190 | 0.869 |
|  | norB-Quinolone | 0.546 | 172 | 0.787 |
|  | acrF-MDR | 0.540 | 170 | 0.777 |
|  | mefA-MLS | 0.521 | 164 | 0.750 |
|  | oprC-MDR | 0.511 | 161 | 0.736 |
|  | tetP-Tetracycline | 0.441 | 139 | 0.636 |
|  | mexY-MDR | 0.413 | 130 | 0.594 |
|  | major_facilitator_superfamily_transporter (MFS)-MDR | 0.406 | 128 | 0.585 |
|  | bpeF-MDR | 0.403 | 127 | 0.581 |
|  | mdtF-MDR | 0.289 | 91 | 0.416 |
|  | adeB-MDR | 0.245 | 77 | 0.352 |
|  | mexE-MDR | 0.232 | 73 | 0.334 |
|  | mexC-MDR | 0.216 | 68 | 0.311 |
|  | oprM-MDR | 0.203 | 64 | 0.293 |
|  | adeJ-MDR | 0.191 | 60 | 0.274 |
|  | mexT-MDR | 0.187 | 59 | 0.270 |
|  | bcrA-Bacitracin | 0.181 | 57 | 0.261 |
|  | cAMP-regulatory protein-Unclassified | 0.181 | 57 | 0.261 |
|  | smeB-MDR | 0.178 | 56 | 0.256 |
|  | smeD-MDR | 0.162 | 51 | 0.233 |
|  | cmeB-MDR | 0.152 | 48 | 0.219 |
|  | oprN-MDR | 0.121 | 38 | 0.174 |
|  | tet35-Tetracycline | 0.111 | 35 | 0.160 |
|  | tcmA-Tetracenomycin_C | 0.108 | 34 | 0.155 |
|  | chloramphenicol exporter-Chloramphenicol | 0.092 | 29 | 0.133 |
|  | emrB-MDR | 0.089 | 28 | 0.128 |
|  | tetA-Tetracycline | 0.079 | 25 | 0.114 |
|  | mexA-MDR | 0.067 | 21 | 0.096 |
|  | macA-MLS | 0.054 | 17 | 0.078 |
|  | mexX-MDR | 0.054 | 17 | 0.078 |
|  | EmrB-QacA family major facilitator transporter-MDR | 0.048 | 15 | 0.069 |
|  | smeF-MDR | 0.048 | 15 | 0.069 |
|  | tetV-Tetracycline | 0.048 | 15 | 0.069 |
|  | acrA-MDR | 0.044 | 14 | 0.064 |
|  | oprA-MDR | 0.044 | 14 | 0.064 |
|  | tetC-Tetracycline | 0.038 | 12 | 0.055 |
|  | oprJ-MDR | 0.035 | 11 | 0.050 |
|  | cmlA-Chloramphenicol | 0.032 | 10 | 0.046 |
|  | mdtK-MDR | 0.029 | 9 | 0.041 |
|  | sdeY-MDR | 0.029 | 9 | 0.041 |
|  | tetG-Tetracycline | 0.029 | 9 | 0.041 |
|  | cmrA-Chloramphenicol | 0.025 | 8 | 0.037 |
|  | adeC-MDR | 0.022 | 7 | 0.032 |
|  | tetR-Tetracycline | 0.022 | 7 | 0.032 |
|  | lmrB-MLS | 0.019 | 6 | 0.027 |
|  | TolC-MDR | 0.019 | 6 | 0.027 |
|  | mdfA-MDR | 0.016 | 5 | 0.023 |
|  | tet41-Tetracycline | 0.016 | 5 | 0.023 |
|  | adeK-MDR | 0.013 | 4 | 0.018 |
|  | bicyclomycin-multidrug_efflux_protein_bcr-MDR | 0.013 | 4 | 0.018 |
|  | emrA-MDR | 0.013 | 4 | 0.018 |
|  | emrD-MDR | 0.010 | 3 | 0.014 |
|  | mdtA-MDR | 0.010 | 3 | 0.014 |
|  | abeS-MDR | 0.006 | 2 | 0.009 |
|  | mdtD-MDR | 0.006 | 2 | 0.009 |
|  | mdtE-MDR | 0.006 | 2 | 0.009 |
|  | mdtG-MDR | 0.006 | 2 | 0.009 |
|  | mdtH-MDR | 0.006 | 2 | 0.009 |
|  | mdtL-MDR | 0.006 | 2 | 0.009 |
|  | tet39-Tetracycline | 0.006 | 2 | 0.009 |
|  | tet40-Tetracycline | 0.006 | 2 | 0.009 |
|  | chloramphenicol and florfenicol exporte-Chloramphenicol | 0.003 | 1 | 0.005 |
|  | mexH-MDR | 0.003 | 1 | 0.005 |
|  | opcM-MDR | 0.003 | 1 | 0.005 |
|  | smeC-MDR | 0.003 | 1 | 0.005 |
| Antibiotic inactivation | ADP-ribosylating transferase_arr-Rifamycin | 0.680 | 214 | 0.979 |
|  | aadA-Aminoglycoside | 0.311 | 98 | 0.448 |
|  | rifampin monooxygenase-Rifamycin | 0.273 | 86 | 0.393 |
|  | aph(6)-I-Aminoglycoside | 0.171 | 54 | 0.247 |
|  | aph(3')-I-Aminoglycoside | 0.137 | 43 | 0.197 |
|  | catB-Chloramphenicol | 0.133 | 42 | 0.192 |
|  | cat_chloramphenicol acetyltransferase-Chloramphenicol | 0.133 | 42 | 0.192 |
|  | fosB-Fosfomycin | 0.130 | 41 | 0.187 |
|  | aac(3)-I-Aminoglycoside | 0.124 | 39 | 0.178 |
|  | THIN-B-Betalactams | 0.076 | 24 | 0.110 |
|  | ereA-MLS | 0.067 | 21 | 0.096 |
|  | aac(3)-IV-Aminoglycoside | 0.057 | 18 | 0.082 |
|  | aac(6')-II-Aminoglycoside | 0.051 | 16 | 0.073 |
|  | aadE-Aminoglycoside | 0.051 | 16 | 0.073 |
|  | ant(9)-I-Aminoglycoside | 0.051 | 16 | 0.073 |
|  | OXA-209-Betalactams | 0.051 | 16 | 0.073 |
|  | catQ-Chloramphenicol | 0.044 | 14 | 0.064 |
|  | OXA-36-Betalactams | 0.041 | 13 | 0.059 |
|  | class A beta-lactamase-Betalactams | 0.041 | 13 | 0.059 |
|  | OXA-53-Betalactams | 0.038 | 12 | 0.055 |
|  | lnuA-MLS | 0.038 | 12 | 0.055 |
|  | aac(3)-IIIa-Aminoglycoside | 0.032 | 10 | 0.046 |
|  | aac(6')-I-Aminoglycoside | 0.032 | 10 | 0.046 |
|  | CMY-19-Betalactams | 0.032 | 10 | 0.046 |
|  | LCR-1-Betalactams | 0.032 | 10 | 0.046 |
|  | OXA-119-Betalactams | 0.032 | 10 | 0.046 |
|  | OXA-205-Betalactams | 0.032 | 10 | 0.046 |
|  | vatB-MLS | 0.032 | 10 | 0.046 |
|  | NPS-1-Betalactams | 0.029 | 9 | 0.041 |
|  | ampC-Betalactams | 0.029 | 9 | 0.041 |
|  | OXA-21-Betalactams | 0.025 | 8 | 0.037 |
|  | aac(2')-I-Aminoglycoside | 0.022 | 7 | 0.032 |
|  | MOX-4-Betalactams | 0.022 | 7 | 0.032 |
|  | OXA-118-Betalactams | 0.022 | 7 | 0.032 |
|  | OXA-46-Betalactams | 0.022 | 7 | 0.032 |
|  | GES-15-Betalactams | 0.019 | 6 | 0.027 |
|  | kasugamycin resistance protein ksgA-kasugamycin | 0.019 | 6 | 0.027 |
|  | aph(3''')-III-Aminoglycoside | 0.016 | 5 | 0.023 |
|  | lnuB-MLS | 0.016 | 5 | 0.023 |
|  | mphA-MLS | 0.016 | 5 | 0.023 |
|  | oleD-MLS | 0.016 | 5 | 0.023 |
|  | aph(3')-V-Aminoglycoside | 0.013 | 4 | 0.018 |
|  | FEZ-1-Betalactams | 0.013 | 4 | 0.018 |
|  | OXA-142-Betalactams | 0.013 | 4 | 0.018 |
|  | OXA-251-Betalactams | 0.013 | 4 | 0.018 |
|  | OXA-34-Betalactams | 0.013 | 4 | 0.018 |
|  | OXA-37-Betalactams | 0.013 | 4 | 0.018 |
|  | TEM-117-Betalactams | 0.013 | 4 | 0.018 |
|  | fosX-Fosfomycin | 0.013 | 4 | 0.018 |
|  | aac(3)-VII-Aminoglycoside | 0.010 | 3 | 0.014 |
|  | aac(3)-X-Aminoglycoside | 0.010 | 3 | 0.014 |
|  | aac(6')-31-Aminoglycoside | 0.010 | 3 | 0.014 |
|  | ant(2'')-I-Aminoglycoside | 0.010 | 3 | 0.014 |
|  | aph(2'')-III-Aminoglycoside | 0.010 | 3 | 0.014 |
|  | MOX-3-Betalactams | 0.010 | 3 | 0.014 |
|  | OXA-10-Betalactams | 0.010 | 3 | 0.014 |
|  | OXA-2-Betalactams | 0.010 | 3 | 0.014 |
|  | CMY-9-Betalactams | 0.006 | 2 | 0.009 |
|  | GES-14-Betalactams | 0.006 | 2 | 0.009 |
|  | MOX-2-Betalactams | 0.006 | 2 | 0.009 |
|  | OXA-129-Betalactams | 0.006 | 2 | 0.009 |
|  | OXA-141-Betalactams | 0.006 | 2 | 0.009 |
|  | OXA-147-Betalactams | 0.006 | 2 | 0.009 |
|  | OXA-20-Betalactams | 0.006 | 2 | 0.009 |
|  | OXA-226-Betalactams | 0.006 | 2 | 0.009 |
|  | OXA-3-Betalactams | 0.006 | 2 | 0.009 |
|  | TEM-171-Betalactams | 0.006 | 2 | 0.009 |
|  | VEB-1-Betalactams | 0.006 | 2 | 0.009 |
|  | VEB-6-Betalactams | 0.006 | 2 | 0.009 |
|  | class B beta-lactamase-Betalactams | 0.006 | 2 | 0.009 |
|  | class C beta-lactamase-Betalactams | 0.006 | 2 | 0.009 |
|  | metallo-beta-lactamase-Betalactams | 0.006 | 2 | 0.009 |
|  | catA-Chloramphenicol | 0.006 | 2 | 0.009 |
|  | ereB-MLS | 0.006 | 2 | 0.009 |
|  | mgtA-MLS | 0.006 | 2 | 0.009 |
|  | vatG-MLS | 0.006 | 2 | 0.009 |
|  | aac(3)-II-Aminoglycoside | 0.003 | 1 | 0.005 |
|  | aac(3)-VIII-Aminoglycoside | 0.003 | 1 | 0.005 |
|  | aad(6)-Aminoglycoside | 0.003 | 1 | 0.005 |
|  | CMY-1-Betalactams | 0.003 | 1 | 0.005 |
|  | GES-13-Betalactams | 0.003 | 1 | 0.005 |
|  | GES-4-Betalactams | 0.003 | 1 | 0.005 |
|  | LRA-12-Betalactams | 0.003 | 1 | 0.005 |
|  | LRA-19-Betalactams | 0.003 | 1 | 0.005 |
|  | LRA-5-Betalactams | 0.003 | 1 | 0.005 |
|  | LRA-9-Betalactams | 0.003 | 1 | 0.005 |
|  | OCH-8-Betalactams | 0.003 | 1 | 0.005 |
|  | OXA-18-Betalactams | 0.003 | 1 | 0.005 |
|  | OXA-183-Betalactams | 0.003 | 1 | 0.005 |
|  | OXA-5-Betalactams | 0.003 | 1 | 0.005 |
|  | OXA-7-Betalactams | 0.003 | 1 | 0.005 |
|  | OXA-74-Betalactams | 0.003 | 1 | 0.005 |
|  | TEM-1-Betalactams | 0.003 | 1 | 0.005 |
|  | TEM-87-Betalactams | 0.003 | 1 | 0.005 |
|  | VEB-2-Betalactams | 0.003 | 1 | 0.005 |
|  | VEB-7-Betalactams | 0.003 | 1 | 0.005 |
|  | penA-Betalactams | 0.003 | 1 | 0.005 |
|  | vatD-MLS | 0.003 | 1 | 0.005 |
|  | vgbA-MLS | 0.003 | 1 | 0.005 |
| Antibiotic target alteration | bacA-Bacitracin | 9.988 | 3145 | 14.381 |
|  | vanR-Vancomycin | 1.420 | 447 | 2.044 |
|  | vanS-Vancomycin | 0.489 | 154 | 0.704 |
|  | arnA-Polymyxin | 0.248 | 78 | 0.357 |
|  | histidine kinase-Unclassified | 0.137 | 43 | 0.197 |
|  | ermF-MLS | 0.127 | 40 | 0.183 |
|  | vanH-Vancomycin | 0.057 | 18 | 0.082 |
|  | vanA-Vancomycin | 0.044 | 14 | 0.064 |
|  | vanD-Vancomycin | 0.041 | 13 | 0.059 |
|  | vanX-Vancomycin | 0.035 | 11 | 0.050 |
|  | vanC-Vancomycin | 0.032 | 10 | 0.046 |
|  | vanE-Vancomycin | 0.016 | 5 | 0.023 |
|  | vanW-Vancomycin | 0.013 | 4 | 0.018 |
|  | erm(35)-MLS | 0.010 | 3 | 0.014 |
|  | vanG-Vancomycin | 0.010 | 3 | 0.014 |
|  | PBP-1A-Betalactams | 0.006 | 2 | 0.009 |
|  | ermA-MLS | 0.006 | 2 | 0.009 |
|  | ermO-MLS | 0.006 | 2 | 0.009 |
|  | ermX-MLS | 0.006 | 2 | 0.009 |
|  | vanB-Vancomycin | 0.006 | 2 | 0.009 |
|  | vanY-Vancomycin | 0.006 | 2 | 0.009 |
|  | erm(33)-MLS | 0.003 | 1 | 0.005 |
|  | erm(38)-MLS | 0.003 | 1 | 0.005 |
|  | erm(39)-MLS | 0.003 | 1 | 0.005 |
|  | erm(TR)-MLS | 0.003 | 1 | 0.005 |
|  | ermC-MLS | 0.003 | 1 | 0.005 |
|  | vanU-Vancomycin | 0.003 | 1 | 0.005 |
| Antibiotic target protection | otrA-Tetracycline | 0.184 | 58 | 0.265 |
|  | tetM-Tetracycline | 0.114 | 36 | 0.165 |
|  | tetW-Tetracycline | 0.048 | 15 | 0.069 |
|  | lsa-MLS | 0.022 | 7 | 0.032 |
|  | carA-Carbomycin | 0.019 | 6 | 0.027 |
|  | srmB-MLS | 0.013 | 4 | 0.018 |
|  | tetracycline_resistance_protein-Tetracycline | 0.013 | 4 | 0.018 |
|  | tetQ-Tetracycline | 0.010 | 3 | 0.014 |
|  | qnrS-Quinolone | 0.006 | 2 | 0.009 |
|  | tet32-Tetracycline | 0.006 | 2 | 0.009 |
|  | tet36-Tetracycline | 0.006 | 2 | 0.009 |
|  | tet44-Tetracycline | 0.006 | 2 | 0.009 |
|  | tetO-Tetracycline | 0.006 | 2 | 0.009 |
|  | tetT-Tetracycline | 0.003 | 1 | 0.005 |
| Antibiotic target replacement | dfrA12-Trimethoprim | 0.718 | 226 | 1.033 |
|  | sul1-Sulfonamide | 0.400 | 126 | 0.576 |
|  | sul2-Sulfonamide | 0.245 | 77 | 0.352 |
|  | dfrA23-Trimethoprim | 0.038 | 12 | 0.055 |
|  | dfrA1-Trimethoprim | 0.022 | 7 | 0.032 |
|  | dfrA13-Trimethoprim | 0.016 | 5 | 0.023 |
|  | dfrB3-Trimethoprim | 0.006 | 2 | 0.009 |
|  | sul3-Sulfonamide | 0.003 | 1 | 0.005 |
|  | dfrA22-Trimethoprim | 0.003 | 1 | 0.005 |
|  | dfrA5-Trimethoprim | 0.003 | 1 | 0.005 |
|  | dfrB1-Trimethoprim | 0.003 | 1 | 0.005 |
| Reduced permeability to antibiotic | ompR-MDR | 0.635 | 200 | 0.915 |
|  | ompF-MDR | 0.013 | 4 | 0.018 |
|  | omp36-MDR | 0.006 | 2 | 0.009 |
|  | tunicamycin resistance protein-Aminoglycoside | 0.003 | 1 | 0.005 |
| Others | puromycin resistance protein-Puromycin | 0.019 | 6 | 0.027 |
|  | tlcC-MLS | 0.016 | 5 | 0.023 |

**Table S9**. Identified viruses/phages from the river Ganges sediment employing Nanopore sequencing

| Viruses/Phages | Hits |
| --- | --- |
| Pseudomonas phage OBP | 45 |
| Enterobacteria phage lambda | 9 |
| Pseudomonas phage B3 | 1 |
| Microcystis phage Ma-LMM01 | 1 |
| Erwinia phage PhiEaH1 | 1 |
| Caulobacter phage CcrRogue | 1 |
| Burkholderia phage phiE125 | 1 |
| BeAn 58058 virus | 1 |
